# The Impact of Stability Considerations on Genetic Fine-mapping

**DOI:** 10.1101/2023.04.11.536456

**Authors:** Alan J. Aw, Lionel Chentian Jin, Nilah M. Ioannidis, Yun S. Song

**Author notes:** **For correspondence:** (NMI); (YSS). Department of Genetics, University of Pennsylvania, United States of America (Alan J. Aw).

## Abstract

Fine-mapping methods, which aim to identify genetic variants responsible for complex traits following genetic association studies, typically assume that sufficient adjustments for confounding within the association study cohort have been made, e.g., through regressing out the top principal components (i.e., residualization). Despite its widespread use, however, residualization may not completely remove all sources of confounding. Here, we propose a complementary stability-guided approach that does not rely on residualization, which identifies consistently fine-mapped variants across different genetic backgrounds or environments. Simulations show that stability guidance neither outperforms nor underperforms residualization, but each approach picks up different variants considerably often. Critically, prioritizing variants that match between the residualization and stability-guided approaches enhances recovery of causal variants. We further demonstrate the utility of the stability approach by applying it to fine-map eQTLs in the GEUVADIS data. Using 378 different functional annotations of the human genome, including recent deep learning-based annotations (e.g., Enformer), we compare enrichments of these annotations among variants for which the stability and traditional residualization-based fine-mapping approaches agree against those for which they disagree, and find that the stability approach enhances the power of traditional fine-mapping methods in identifying variants with functional impact. Finally, in cases where the two approaches report distinct variants, our approach identifies variants comparably enriched for functional annotations. Our findings suggest that the stability principle, as a conceptually simple device, complements existing approaches to fine-mapping, reinforcing recent advocacy of evaluating cross-population and cross-environment portability of biological findings. To support visualization and interpretation of our results, we provide a Shiny app, available at: https://alan-aw.shinyapps.io/stability_v0/.

## Introduction

An important challenge faced by computational precision health research is the lack of generalizability of biological findings, which are often obtained by studying cohorts that do not include particular communities of individuals. Known as the *cross-population generalizability* or *portability problem*, the challenge persists in multiple settings, including gene expression prediction (***Keys et al., 2020***) and polygenic risk prediction (***Mostafavi et al., 2020***). Generalizable biological signals, such as the functional impact of a variant, are important, as they ensure that general conclusions drawn from cohort-specific analyses are not based on spurious discoveries. Portability problems are potentially attributable to cohort-biased discoveries being treated as generalizable true signals. Efforts to address such problems have included the use of diverse cohorts typically representing multiple population ancestries (***Márquez-Luna et al., 2017***), meta-analyses of earlier studies across diverse cohorts (***Turley et al., 2021***; ***Han and Eskin, 2012***; ***Morris, 2011***; ***Willer et al., 2010***), or focusing on biological markers that are more likely *a priori* to play a causal role (including proxy variables), e.g., rare variants (***Zaidi and Mathieson, 2020***) or the transcriptome (***Liang et al., 2022***).

In this paper, we consider an approach based on the notion of *stability* to improve generalizability. Being a pillar of veridical data science (***Yu and Kumbier, 2020***), stability refers to the robustness of statistical conclusions to *perturbations* of the data. Perturbations are not arbitrary, but instead they encode the practitioner’s beliefs about the quality of the data and the nature of relationship between variables. Well-chosen perturbation schemes will help the practitioner obtain statistical conclusions that are robust, in the sense that they are generalizable rather than spurious findings, and therefore more likely to capture the true signal. For example, in sparse linear models, prioritizing the stability of effect sizes to cross-validation “perturbations” leads to a much smaller set of selected features without reducing predictive performance (***Lim and Yu, 2016***). In another example involving the application of random forests to detect higher-order interactions between gene regulation features (***Basu et al., 2018***), interactions stable to bootstrap perturbations are also largely consistent with known physical interactions between the associated transcription factor binding or enhancer sites.

To further investigate the utility of the stability approach, we focus on a procedure known as (genetic) fine-mapping (***Schaid et al., 2018***). Fine-mapping is the task of identifying genetic variants that causally affect some trait of interest. From the viewpoint of stability, previous works focusing on cross-population stability of fine-mapped variants typically use Bayesian linear models. The linear model has allowed modeling of heterogeneous effect sizes across different user-defined populations (e.g., ethnic groups, ancestrally distinct populations, or study cohorts), and it is common to assume that causal variants share correlated effects across populations (see, e.g., eq. (24) of ***LaPierre et al., 2021*** or eq. (9) of ***Wen et al., 2015***).

Unlike the parametric approaches described above, we here apply a *non-parametric* fine-mapping method to GEUVADIS (***Lappalainen et al., 2013***), a database of gene expression traits measured across individuals of diverse genetic ancestries and from different geographical environments, whose accompanying genotypes are available through the 1000 Genomes Project (1kGP) (1000 ***Genomes Project Consortium et al., 2015***). We apply fine-mapping through two approaches, one that is commonly used in practice and another that is guided by stability. First, we perform simulations using 1kGP genotypes to evaluate the strength of each approach. We next evaluate the agreement of fine-mapped variants between the two approaches on GEUVADIS data, and then measure the functional significance of variants picked by both approaches using a much wider range of functional annotations than considered in previous studies. By performing various statistical tests on the functional annotations, we evaluate the advantages brought about by the incorporation of stability. Finally, we investigate the broader applicability of the stability approach, by implementing and analyzing a stability-guided version of the fine-mapping algorithm SuSiE on simulated data. To support visualization of results obtained from the GEUVADIS analysis at both the single gene and genome-wide levels, we provide an interactive Shiny app which is available at: https://alan-aw.shinyapps.io/stability_v0/. Our app is open-source and designed to support geneticists in interpreting our results, thereby also addressing a growing demand for accessible, *integrative* software to interpret genetic findings in the age of big data and variant annotation databases.

## Results

### Experimental Design

We apply the fine-mapping method PICS (***Taylor et al., 2021***; ***Farh et al., 2015***; see Algorithm 1 in Probabilistic Identification of Causal SNPs) to the GEUVADIS data (***Lappalainen et al., 2013***), which consists of *T* = 22, 720 gene expression traits measured across *N* = 445 individuals, with their accompanying genotypes obtained from the 1000 Genomes Project (1000 ***Genomes Project Consortium et al., 2015***). These individuals are of either European or African ancestry, with about four fifths of the cohort made up of individuals of (self-identified) European descent. In particular, these ancestrally different subpopulations have distinct linkage disequilibrium patterns and environmental exposures, which constitute potential confounders that we wish to stabilize the fine-mapping procedure against.

PICS, like many eQTL analysis methods, requires the lead variant at a locus to compute posterior probabilities, so we perform marginal regressions of each gene expression trait against variants within the fine-mapping locus. Our implementation of PICS generates three sets of variants, which represent putatively causal variants that are (marginally) highly associated, moderately associated and weakly associated with the gene expression phenotype. (See Materials and Methods for details.) As illustrated in Figure 1A, we compare two versions of the fine-mapping procedure — one that is typically performed in practice, and another that is motivated by the stability approach.

**Figure 1.**
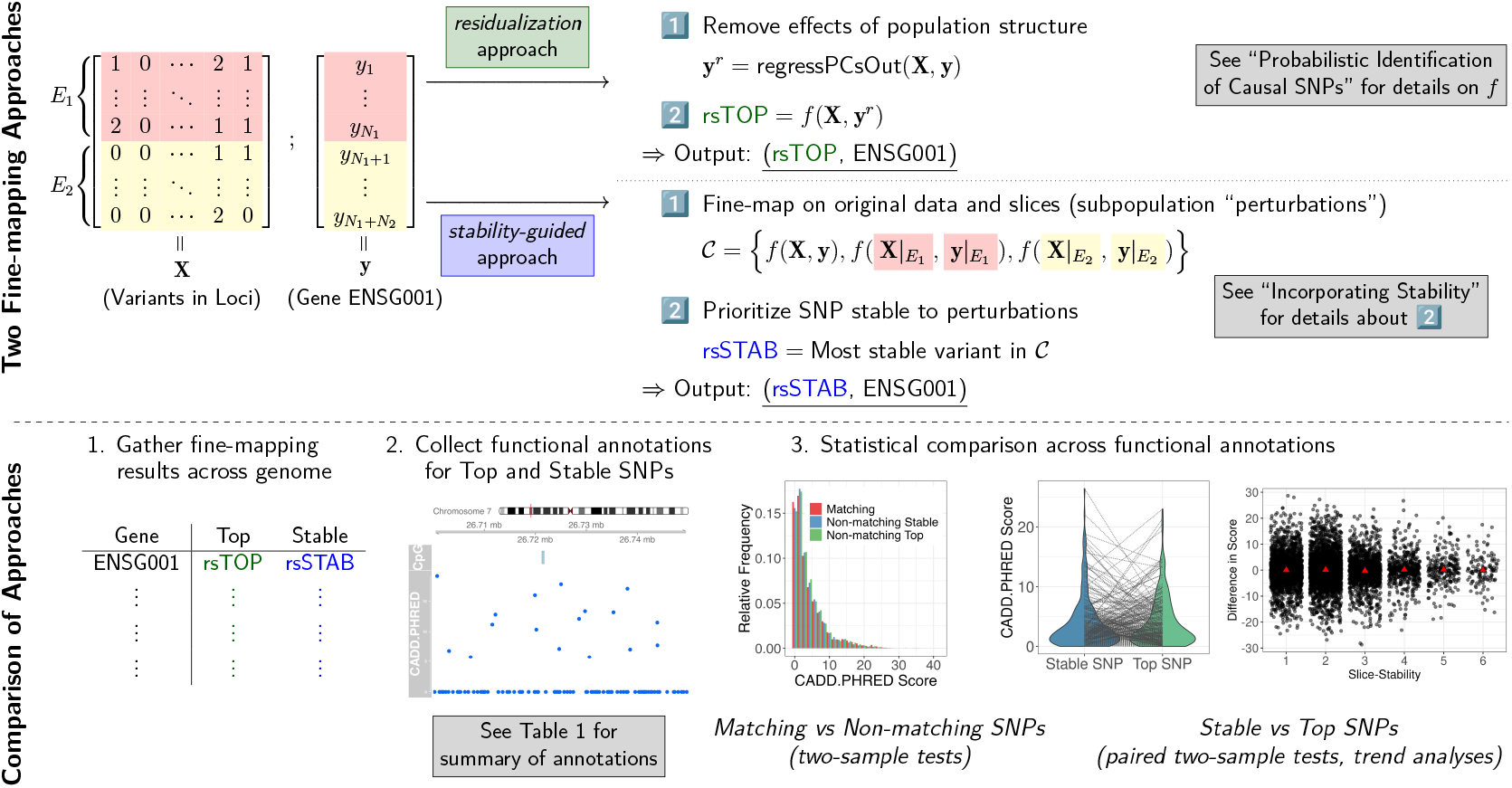
An overview of our study of the impact of stability considerations on genetic fine-mapping. **A**. The two ways in which we perform fine-mapping, the first of which (colored in green) prioritizes the stability of variant discoveries to subpopulation perturbations. The data illustrates the case where there are two distinct environments, or subpopulations (denoted *E*_1_ and *E*_2_), that split the observations. **B**. Key steps in our comparison of the stability-guided approach with the popular residualization approach.

#### Stable variant

Specifically, to incorporate stability into fine-mapping and obtain what we call the *stable variant* or stable SNP, we encode into the algorithm our belief that for a gene expression trait, a causal variant would presumably act through the same mechanism regardless of the population from which they originate. Indeed, if a set of genetic variants were causal, the same algorithm should report it, if run on subsets of the data corresponding to heterogeneous populations. This belief is consistent with recent simulation studies showing that the inclusion of GWAS variants discovered in diverse populations both mitigates false discoveries driven by linkage disequilibrium differences (***Li et al., 2022***) and improves generalizability of polygenic score construction (***Cavazos and Witte, 2021***). Hence, the stable variant is the variant with the highest probability of being causal, conditioned on being reported by PICS in the most number of subpopulations. Incorporating Stability provides a formal definition of the stable variant.

#### Top variant

The stability consideration we have described is closely related to a popular procedure known as correction for population structure, which residualizes the trait using measured confounders (e.g., top principal components computed from the genotype matrix). We call the variant returned by the residualization approach the *top variant* or top SNP. The residualization approach removes the effects of genetic ancestry and environental exposures on a trait, and is used to avoid the risk of low statistical power borne by stratified analyses. The top variant is formally defined towards the end of Probabilistic Identification of Causal SNPs. We remark that the top variant is a function of the residualized trait, as opposed to the stable variant, which is a function of the unresidualized trait (see Figure 1A).

### Simulation Study

We simulated 100 genes from GEUVADIS gene expression data, sampled proportionally across the 22 autosomes. Mirroring the approach described in Section 4 of ***Wang et al. (2020***), synthetic gene expression phenotypes were simulated based on the sampled genes, and involve two parameters: *C*, the number of causal variants, and *ϕ*, the proportion of variance in gene expression explained by variants in *cis*. Whenever available, gene expression canonical TSS was used to include only variants lying within 1 Mb upstream and downstream when simulating gene expression phenotypes. We considered all combinations of *C ∈* {1, 2, 3} and *ϕ ∈* {0.05, 0.1, 0.2, 0.4}, and simulated two replicates for each gene and for each combination of *C* and *ϕ*. Altogether, we generated 100× 3× 4× 2 = 2, 400 datasets, on which we ran PICS and SuSiE. For PICS, we ran four versions: the *stability-guided* version and *residualization* version, which return the stable variant and top variant respectively; a *combined* version, which runs stability-guided PICS on covariate-residualized phenotypes; as well as a *plain* version, where no correction for population stratification or stability guidance was applied. For SuSiE, we ran two versions: the stability-guided version and the residualization version. We also ran a separate set of simulations comprising six scenarios of environmental heterogeneity by ancestry on 10 genes from Chromosomes 1, 20, 21 and 22 (6 × 10 × 3 × 4 × 2 = 1, 440 generated datasets), on which we ran the plain and stability-guided versions of PICS.

#### Evaluation on simulated genes

Following ***Mazumder (2020***), we analyze the recovery probability of all fine-mapping algorithms as a function of the Signal-to-Noise Ratio (SNR), defined here as the ratio of true signal to background noise: SNR = var(**Xb**)/*σ*^2^(see Simulation Study and Evaluation Details in Materials and Methods for details).

### Analysis of GEUVADIS Data

We compare the results obtained by using each approach, across a range of categories of functional annotations including conservation, pathogenicity, chromatin accessibility, transcription factor binding and histone modification, and across biological assays and computational predictions. See Table 1 for full list of annotations covered. (Supplementary File 1 Section S5 contains a full description of each quantity and its interpretation.)

**Table 1.** A list of 378 functional annotations across which the biological significances of stable and top fine-mapped single nucleotide polymorphisms are compared. Annotations that report multiple scores have the total number of scores reported shown in parentheses. Scores mined from the FAVOR database (***Zhou et al., 2023***) are indicated by an asterisk. (*TSS* = Transcription Start Site, *bp* = base pair)

| Functional Annotation Type | Functional Annotation |
| --- | --- |
| Ensembl | Distance to Canonical TSS (Cunningham et al., 2022)<br>Regulatory Features (6; Cunningham et al., 2022) |
| Computational Predictions | CADD* (2; Rentzsch et al., 2019)<br>SIFTVal* (Ng and Henikoff, 2003)<br>FATHMM-XF* (Rogers et al., 2018)<br>LINSIGHT* (Huang et al., 2017)<br>Polyphen* (Adzhubei et al., 2010)<br>PhyloP* (3; Pollard et al., 2010)<br>Gerp* (2; Davydov et al., 2010)<br>B Statistic* (McVicker et al., 2009)<br>FunSeq2* (Fu et al., 2014)<br>ALoFT* (Balasubramanian et al., 2017)<br>Percent CpG in 75bp window* (Rentzsch et al., 2019)<br>Percent GC in 75bp window* (Rentzsch et al., 2019)<br>FIRE (Ioannidis et al., 2017)<br>Enformer (177 tracks × 2 scores per track; Avsec et al., 2021) |

Figure 1B summarizes the key steps of our investigation. To be clear, a first comparison is between the set of genes for which the top and stable variants match and the set of genes for which the top and stable variants disagree; comparisons are made between matching variants and one of the non-matching sets of variants (top or stable). A second comparison is restricted to the set of genes with non-matching top and stable variants, i.e., between *paired* sets of variants fine-mapped to a gene, where one set of a pair corresponds to the output of the residualization approach whereas the other corresponds to the output of the stability-guided approach. We additionally compare across three fine-mapping output sets, corresponding to variants that have a high, moderate or weak marginal association with the expression phenotype. For brevity, we term these three sets Potential Set 1, Potential Set 2 and Potential Set 3, respectively. (See Algorithm 1 for details.) For each Potential Set, we find the top and stable variants as described in Incorporating Stability.

### Simulations justify exploration of stability guidance

To better understand how stability guidance may support the discovery of causal variants across ancestrally diverse cohorts that possess confounding exogenous factors, we selected 10 genes from Chromosomes 1, 20, 21 and 22 to simulate six scenarios where environmental heterogeneity by ancestry influences gene expression. We consider four scenarios where environmental *variance* differed between ancestries and two scenarios where environmental *mean* differed between ancestries. We also vary the number of causal variants and proportion of variance in gene expression expressed in *cis*. We run Stable PICS, which returns stable variants; alongside a “plain” version of PICS (Plain PICS), which neither performs PC-residualization of phenotypes nor incorporates stability.

We find that, across these simulations, performance was not significantly different between Stable PICS and Plain PICS: both approaches recover the true causal variant in Potential Set 1 with similar frequencies in simulations involving one causal variant, and in simulations involving two or more causal variants the leading potential sets recover at least one causal variant with similar frequency (Figure 2A). These also hold for lower potential sets (Figure 2—figure supplement 1). While the two approaches agree more often than not, there is greater disagreement when considering simulations with low SNR (which may potentially reflect real gene expression data — see Stable variants frequently do not match top variants in GEUVADIS). For example, we identified 83% matching variants in Potential Set 1 across all simulations, but this dropped to 74% when restricting to simulations with *ϕ* = 0.05. We next split results by whether the variant returned by Plain PICS had a large posterior probability (PP) — defined as *>* 0.9 to reflect thresholds reported in practice — and found strong agreement between the two approaches when PP was large: for Potential Set 1 the agreement was 98%, Potential Set 2 was 90%, and Potential Set 3 was 93% (Supplementary File 2A). These observations together imply that the two approaches disagree particularly when SNR is low and the PP of variants returned by Plain PICS is not large, and Stable PICS recovers causal variants accurately in certain cases where Plain PICS fails to do so (otherwise their performances would be different).

**Figure 2.**
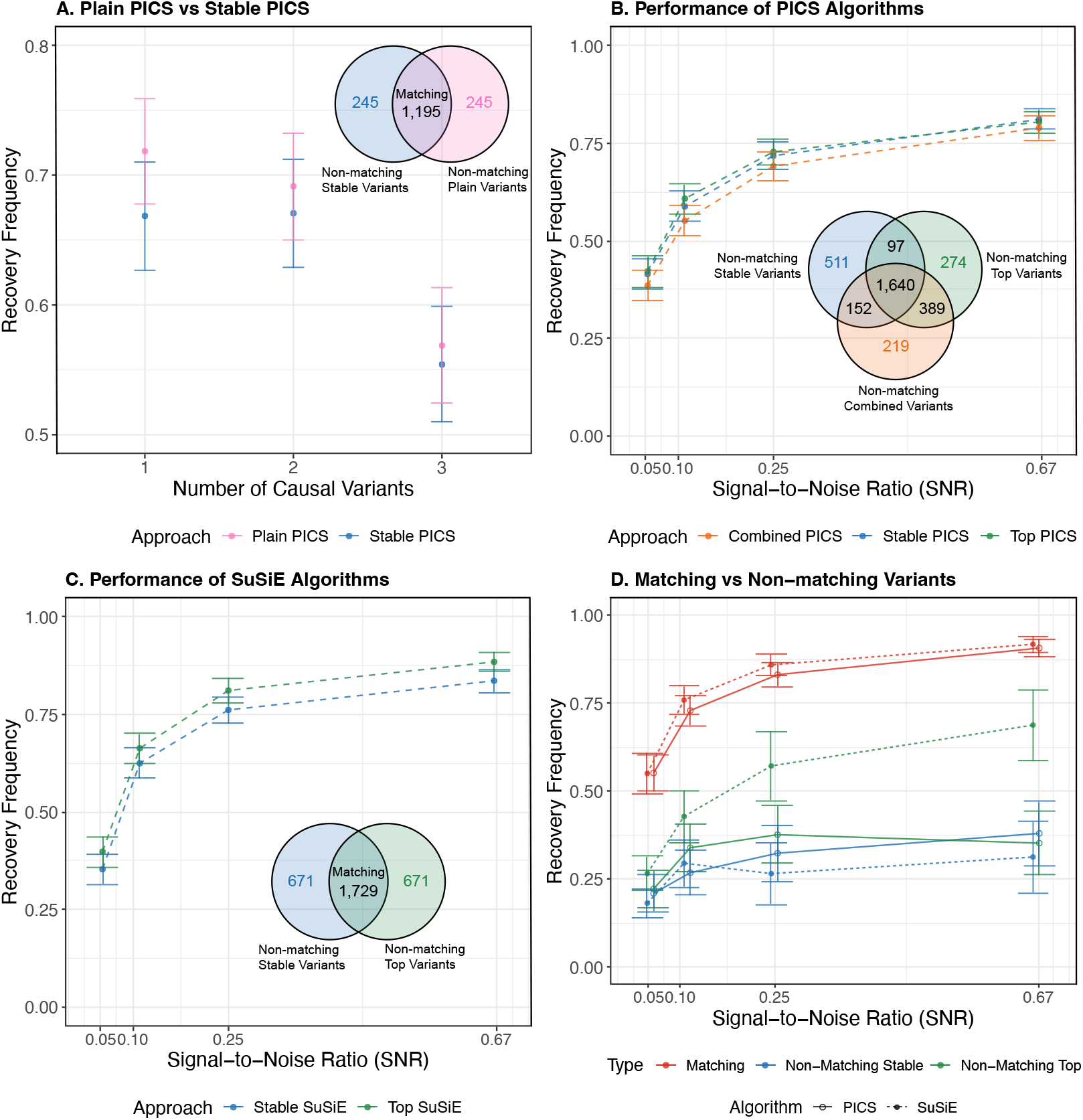
Simulation study results. **A**. The frequency with which at least one causal variant is recovered in Potential Set 1 by Plain PICS and Stable PICS, across 1, 440 simulated gene expression data that incorporate ancestry-mediated environmental heterogeneity. Recovery frequencies are stratified by simulations differing in the number of causal variants, and the Venn diagram reports the number of matching and non-matching variants in Potential Set 1 across all simulations. **B**. The frequency with which at least one causal variant is recovered in Potential Set 1 by Combined PICS, Stable PICS and Top PICS, across 2, 400 simulated gene expression data. Recovery frequencies are stratified by the SNR parameter *ϕ* used in simulations, and the Venn diagram reports the number of matching and non-matching variants in Potential Set 1 across all simulations. **C**. The frequency with which at least one causal variant is recovered in Credible Set 1 by Stable SuSiE and Top SuSiE. Venn diagram reports the number of matching and non-matching variants in Potential Set 1 across all simulations. **D**. The frequency with which matching and non-matching variants in the first credible or potential set recover a causal variant, obtained from comparing top and stable approaches to an algorithm. **Figure 2—figure supplement 1**. Plain PICS vs Stable PICS (Potential Sets 2 and 3) **Figure 2—figure supplement 2**. Performance of PICS Algorithms (Potential Sets 2 and 3) **Figure 2—figure supplement 3**. Performance of SuSiE Algorithms (Credible Sets 2 and 3) **Figure 2—figure supplement 4**. Matching vs Non-matching Variants (Potential and Credible Sets 1 and 2) **Figure 2—figure supplement 5**. Stable PICS vs Stable SuSiE (1 causal variant) **Figure 2—figure supplement 6**. Stable PICS vs Stable SuSiE (2 causal variants) **Figure 2—figure supplement 7**. Stable PICS vs Stable SuSiE (3 causal variants) **Figure 2—figure supplement 8**. Matching Top vs Stable SNP posterior probabilities **Figure 2—figure supplement 9**. Non-matching Top vs Stable SNP posterior probabilities **Figure 2—figure supplement 10**. Stable PICS vs SuSiE (1 causal variant) **Figure 2—figure supplement 11**. Stable PICS vs SuSiE (2 causal variants) **Figure 2—figure supplement 12**. Stable PICS vs SuSiE (3 causal variants) **Figure 2—figure supplement 13**. Distribution of number of variants recovered by PICS (1 causal variant) **Figure 2—figure supplement 14**. Distribution of number of variants recovered by PICS (2 causal variants) **Figure 2—figure supplement 15**. Distribution of number of variants recovered by PICS (3 causal variants) **Figure 2—figure supplement 16**. Distribution of number of variants recovered by SuSiE (1 causal variant) **Figure 2—figure supplement 17**. Distribution of number of variants recovered by SuSiE (2 causal variants) **Figure 2—figure supplement 18**. Distribution of number of variants recovered by SuSiE (3 causal variants) **Figure 2—figure supplement 19**. Variant recovery frequency of PICS and SuSiE matching and non-matching variants (1 causal variant) **Figure 2—figure supplement 20**. Distribution of number of variants recovered by PICS matching and non-matching variants (2 causal variants) **Figure 2—figure supplement 21**. Distribution of number of variants recovered by SuSiE matching and non-matching variants (2 causal variants) **Figure 2—figure supplement 22**. Distribution of number of variants recovered by PICS matching and non-matching variants (3 causal variants) **Figure 2—figure supplement 23**. Distribution of number of variants recovered by SuSiE matching and non-matching variants (3 causal variants) **Figure 2—figure supplement 24**. Plain PICS vs Stable PICS in environmental heterogeneity Simulations (1 causal variant) **Figure 2—figure supplement 25**. Plain PICS vs Stable PICS in environmental heterogeneity simulations (2 causal variants) **Figure 2—figure supplement 26**. Plain PICS vs Stable PICS in environmental heterogeneity simulations (3 causal variants) **Figure 2—figure supplement 27**. Plain PICS vs Stable PICS in “variance shift (*t* = 8)” environmental heterogeneity simulations **Figure 2—figure supplement 28**. Plain PICS vs Stable PICS in “variance shift (*t* = 16)” environmental heterogeneity simulations **Figure 2—figure supplement 29**. Plain PICS vs Stable PICS in “variance shift (*t* = 128)” environmental heterogeneity simulations **Figure 2—figure supplement 30**. Plain PICS vs Stable PICS in “variance shift (*t* = 256)” environmental heterogeneity simulations **Figure 2—figure supplement 31**. Plain PICS vs Stable PICS in “mean shift (|*i* − 3|)” environmental heterogeneity simulations **Figure 2—figure supplement 32**. Plain PICS vs Stable PICS in “mean shift (*i* = 3)” environmental heterogeneity simulations

Looking closely at each simulation scenario, we see that Stable PICS recovers a causal variant more frequently than Plain PICS in scenarios where there are multiple causal variants and environmental heterogeneity is driven by a “spiked mean shift” (Simulation Study and Evaluation Details), while Plain PICS generally outperforms when environmental heterogeneity is driven by differences in environmental variation. For example, in a set of spiked mean shift simulations with three causal variants, with low signal-to-noise ratio (*ϕ* = 0.1) and with exogenous variable drawn from *N*(2*σ, σ*^2^) for only GBR individuals (rest are drawn from *N*(0, *σ*^2^)), the frequencies at which at least one causal variant is recovered were 80% and 75% for Stable PICS and Plain PICS, respectively (see Figure 2— figure supplement 32). However, none of these differences in performance is statistically significant. In summary, these findings demonstrate that non-genetic confounding in cohorts can reduce power in methods not adjusting or accounting for ancestral confounding, but can be remedied by approaches that do so. (Performances stratified by SNR and number of potential sets included are summarized in Figure 2—figure supplements 24-26; performances stratified by each scenario are summarized in Figure 2—figure supplements 27-32)

### Simulations reveal advantages (and disadvantages) of stability guidance

We next conduct a larger set of simulations using 100 genes selected across all 22 autosomes. Similar to the earlier set of simulations, we vary both the number of causal variants and proportion of variance in gene expression explained by variants in *cis*. However, here we compare stability-guided fine-mapping against residualization fine-mapping, a commonly used approach.

#### No difference in power between Stability guidance and Residualization, although consider ably many variants do not match

Comparing stable and top variants returned by Stable PICS and PICS run via the residualization approach (Top PICS), we observed similar causal variant recovery rates. Across all 2, 400 simulated gene expression phenotypes, the stable variant in Potential Set 1 was causal with frequency 0.63, which is close to and not significantly different from the frequency at which the Potential Set 1 top variant was causal (0.64; McNemar test *p*-value = 0.38). Similarly close frequencies were observed for lower potential sets (Potential Set 2: stable = 0.12,top = 0.13; Potential Set 3: stable = 0.041,top = 0.047), as well as when results were stratified by SNR (Figure 2B). However, top and stable variants often disagreed, with Potential Set 1 having 72% matching top and stable variants, Potential Set 2 matching at 40% and Potential Set 3 matching at 25% (Figure 2—figure supplement 2). Stratifying results by simulation parameters (number of causal variants, *S*, and SNR, *ϕ*), we generally observed matching fractions increasing with SNR, at least in Potential Set 1: for example, in simulations involving one causal variant, the matching fraction is 69.5% under SNR = 0.05/(1 − 0.05) ≈ 0.053, and it increases monotonically to 89.5% under SNR = 0.4/(1 − 0.4) ≈ 0.67 (Supplementary File 2B). There was no clear relationship between matching fractions and the number of causal variants simulated.

#### Searching for matching variants between Top PICS and Stable PICS improves causal variant recovery

Given that stable and top variants frequently do not match despite achieving similar performance, we suspect that, similar to the smaller set of simulations on which Plain PICS and Stable PICS were compared, each approach can recover causal variants in scenarios where the other does not. We thus explore ways to combine the residualization and stability-driven approaches, by considering (i) combining them into a single fine-mapping algorithm (we call the resulting procedure *Combined PICS*); and (ii) prioritizing matching variants between the two algorithms. Comparing the performance of Combined PICS against both Top and Stable PICS, however, we find no significant difference in its ability to recover causal variants (Figure 2B). This conclusion held even when we analyzed performance by stratifying simulations by SNR and number of causal variants simulated (*ϕ* and *S* parameters; see Figure 2—figure supplements 13-15). On the other hand, matching variants between Top and Stable PICS are significantly more likely to be causal. Across all simulations, a matching variant in Potential Set 1 was 2.5× as likely to be causal than either a non-matching top or stable variant (Figure 2D) — a result that was qualitatively consistent even when we stratified simulations by SNR and number of causal variants simulated (Figure 2—figure supplements 19, 20 and 22). A similar trend was observed for Potential Set 2, although we do not see much higher causal variant recovery when comparing matching and non-matching variants in Potential Set 3 (Figure 2—figure supplement 4).

#### Stability guidance improves causal variant recovery in SuSiE

To explore the applicability of stability guidance beyond PICS, we developed a similar procedure that runs the fine-mapping algorithm SuSiE (***Wang et al., 2020***) on multiple slices and returns a stable variant. We call this approach *Stable SuSiE*. We then compared Stable SuSiE against SuSiE applied to residualized phenotypes (*Top SuSiE*), analogous to our comparisons for PICS earlier. First, similar to PICS, we generally observe no significance differences in performance between Stable and Top SuSiE, although empirically more causal variants are recovered by Top SuSiE in larger SNR simulation settings (Figure 2C), while more causal variants are recovered by Stable SuSiE in lower credible sets (Figure 2—figure supplement 3; see also Figure 2—figure supplements 16-18). Similar to PICS, we also observed matching fractions increasing with SNR in Potential Set 1 (Supplementary File 2C). Next, comparing matching versus non-matching variants between the approaches, we observe a boost in causal variant recovery similar to that observed in PICS (Figure 2D), which persisted even when we stratified results by simulation parameters (Figure 2—figure supplements 19, 21 and 23). However, one difference is that, unlike in PICS, this boost was unique to Credible Set 1 — for the other two credible sets improvements in causal variant recovery were observed only when SNR is high, with non-matching Stable SuSiE variants recovering at least one causal variant with the highest frequency in low SNR scenarios (Figure 2—figure supplement 4).

#### Other analyses

Our key findings from the simulation study are as follows. Stability guidance, on its own, does not significantly outperform nor underperform other standard approaches at causal variant recovery. However, prioritizing variants that agree between stability-guided fine-mapping and standard fine-mapping can significantly boost causal variant recovery. In the Supplement, we also describe findings from investigations into the impact of including more potential sets on matching frequency and causal variant recovery (Supplementary File 1 Section S2), the differences in posterior probability between non-matching variants (Supplementary File 1 Section S3), the relative performance of SuSiE and Stable PICS (Supplementary File 1 Section S4), and interpreting matching variants with very low (stable) posterior probability (Supplementary File 1 Section S11).

### Stable variants frequently do not match top variants in GEUVADIS

Having identified scenarios that favor stability guidance, as well as strategies for using stability guidance to recover causal variants, we now turn to analysis of real gene expression data from GEUVADIS. As a preliminary analysis, when interrogating if the residualization and stability-guided approaches produce the same candidate causal variants, we find that for Potential Set 1, which corresponds to the set typically reported in fine-mapping studies, 56.2% of genes had matching top and stable variants (see Figure 3). Moving down potential sets, we find less agreement (Potential Set 2: 36.2%, Potential Set 3: 25.6%), providing evidence that as the marginal association of a variant with the expression phenotype decreases, the two approaches prioritize different signals when searching for putatively causal variants. This result is most consistent with our simulations using *S* = 3 causal variants and low SNR (*ϕ* = 0.05), where we observed 56.5%, 37% and 24.5% matching top and stable variants in Potential Sets 1, 2 and 3 respectively.

**Figure 3.**
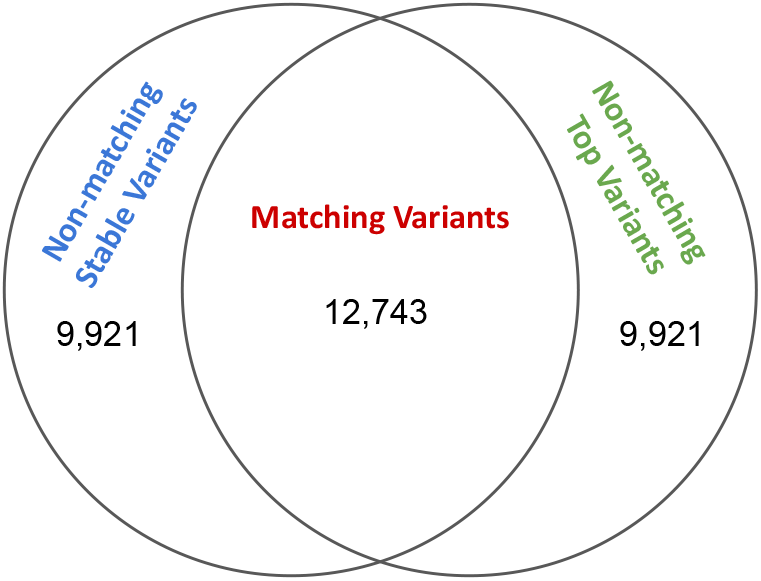
Venn diagram showing the number of matching and non-matching variants for Potential Set 1 in GEUVADIS fine-mapped variants. **Figure 3—figure supplement 1**. Matching GEUVADIS Top vs Stable SNP posterior probabilities **Figure 3—figure supplement 2**. Non-matching GEUVADIS Top vs Stable SNP posterior probabilities

Because our simulations demonstrate that prioritizing matching variants boosts causal variant recovery, we proceeded with comparing functional impact of matching stable and top variants against non-matching variants. For the rest of this Section, we report only results for Potential Set 1, given that it corresponds to the set typically reported in fine-mapping. Results for the other potential sets are provided in Supplementary File 1 Sections S7 and S8.

### Matching versus Non-Matching Variants

For each gene, our algorithm finds the top eQTL variant and the stable eQTL variant, which may not coincide. We thus run (one-sided) unpaired Wilcoxon tests on matching and non-matching sets of variants, to detect significant functional enrichment of one set of variants over the other. We find that the top variants that are also stable for the corresponding gene (*N*_match_ = 12, 743 maximum annotatable) score significantly higher in functional annotations than the top variants that are not stable (*N*_non-match_ = 9, 921 maximum annotatable). Notably, 361 out of 378 functional annotations report one-sided greater *p*-values *<* 0.05 for the matching (i.e., both top and stable) variants after correcting for multiple testing using the Benjamini-Hochberg (BH) procedure, with many of these annotations measuring magnitudes of functional impact or functional enrichment (e.g., Enformer perturbation scores, FATHMM.XF score). Amongst the 17 remaining functional annotations, none has significantly lower scores for the matching variants. (Supplementary File 1 Section S7 lists these functional annotations in detail.) We also find that empirically, the matching variants tend to have greater agreement in posterior probabilities than non-matching variants (Figure 2—figure supplements 1 and 2).

As an example of a significant functional annotation, consider raw CADD scores (***Rentzsch et al., 2019***) — a higher value of which indicates a greater likelihood of deleterious effects. Out of the 22, 559 genes for which both the top and the stable variant are annotatable, looking at the distribution of top variant scores, the one corresponding to the 12, 685 genes with matching top and stable variants stochastically dominates the one corresponding to the remaining 9, 874 genes (Figure 4A). This relationship is more pronounced when we inspect PHRED-scaled CADD scores, where we apply a sliding cutoff threshold for calling variant deleteriousness (i.e., potential pathogenicity — see ***Rentzsch et al., 2019***). We find that a greater proportion of matching variants than of either non-matching variant is classified as deleterious under the typical range of deleteriousness cutoffs (Figure 4B).

**Figure 4.**
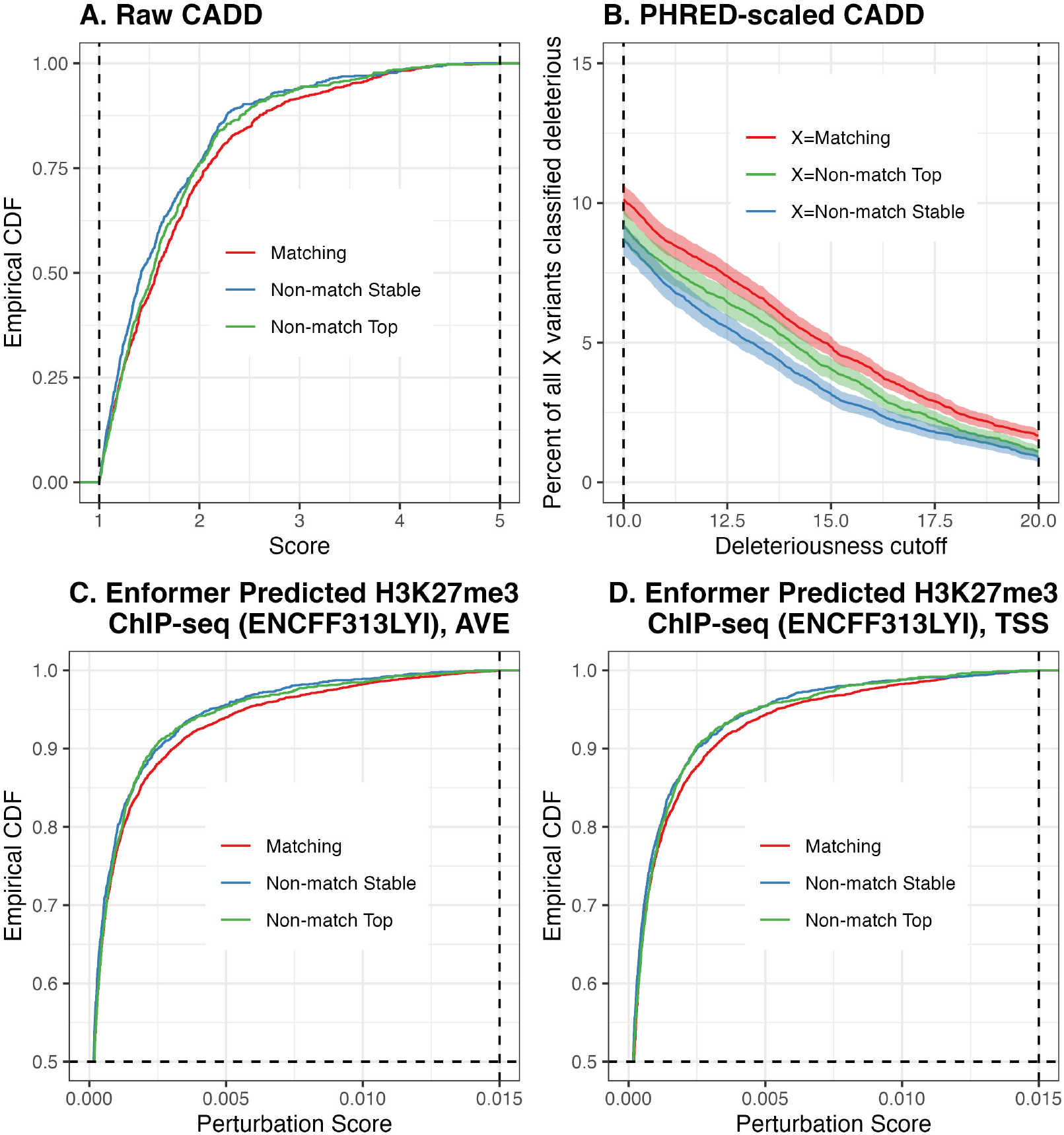
*Top Row*. CADD scores. **A**. Empirical cumulative distribution functions of raw CADD scores of matching and non-matching variants across all genes, for Potential Set 1. Non-matching variants are further divided into stable and top variants, with a score lower threshold of 1.0 and upper threshold of 5.0 used to improve visualization. **B**. For a deleteriousness cutoff, the percent of (i) all matching variants, (ii) all non-matching top variants, and (iii) all non-matching stable variants, which are classified as deleterious. We use a sliding cutoff threshold ranging from 10 to 20 as recommended by CADD authors. *Bottom Row*. Empirical cumulative distribution functions of perturbation scores of Enformer-predicted H3K27me3 ChIP-seq track. Score upper threshold of 0.015 and empirical CDF lower threshold of 0.5 used to improve visualization. **C**. Perturbation scores computed from predictions based on centering input sequences on the gene TSS as well as its two flanking positions. **D**. Perturbation scores computed from predictions based on centering input sequences on the gene TSS only.

Another example demonstrating significant functional enrichment of matching variants over non-matching variants is the perturbation scores on H3K9me3 ChIP-seq peaks, as predicted by the Enformer (***Avsec et al., 2021***). Out of the 6, 364 genes for which the distance of both the top and the stable variant to the TSS are within the Enformer input sequence length constraint, looking at the the distribution of top variant scores, the one corresponding to the 4, 491 genes with matching top and stable variants stochastically dominates the one corresponding to the remaining 1, 873 genes (Figure 4C,D). This relationship is true regardless of whether perturbation scores are calculated from an average of input sequences centered on the gene TSS and its two flanking positions (Figure 4C), or from input sequences centered on the gene TSS only (Figure 4D). (See Supplementary File 1 Section S6 for details on Enformer annotation calculation.)

Similarly, amongst all stable variants, those variants that match the top variant are significantly more enriched in functional annotations: 363 functional annotations report one-sided greater BH adjusted *p*-values *<* 0.05 for the matching variants, and none of the remaining 15 annotations present significant depletion for the matching variants set.

### Top versus Stable variants when they do not match

Focusing on the genes for which the top and stable variants are different, we run (one-sided) paired Wilcoxon tests to detect significant functional enrichment of one set of variants over the other. We find in general that for some genes stable variants can carry more functional impact than top variants, and for other genes top variants carry more functional impact — although neither of these patterns is statistically significant genome-wide after multiple testing correction using the BH procedure. For example, for raw CADD scores, out of the 9, 874 genes for which the top and the stable variants do not match, 4, 906 genes have higher scoring stable variants than top variants, whereas 4, 968 genes have higher scoring top variants than stable variants (Figure 5A). Looking at the accompanying PHRED-scaled CADD scores (CADD PHRED) for Potential Set 1, when applying a sliding cutoff threshold for calling variant deleteriousness, we find that even though a higher fraction of genes have top but not stable variant classified as deleterious, the difference is usually not significant (Figure 5B). Figure 5B also demonstrates that no matter how the cutoff is chosen, there are genes for which the top variant is not classified deleterious while the stable variant is. Taken together, these observations suggest that the stability-guided approach can sometimes be more useful at identifying variants of functional significance, and in a broad sense both fine-mapped variants should be equally prioritized for potential of carrying functional impact.

**Figure 5.**
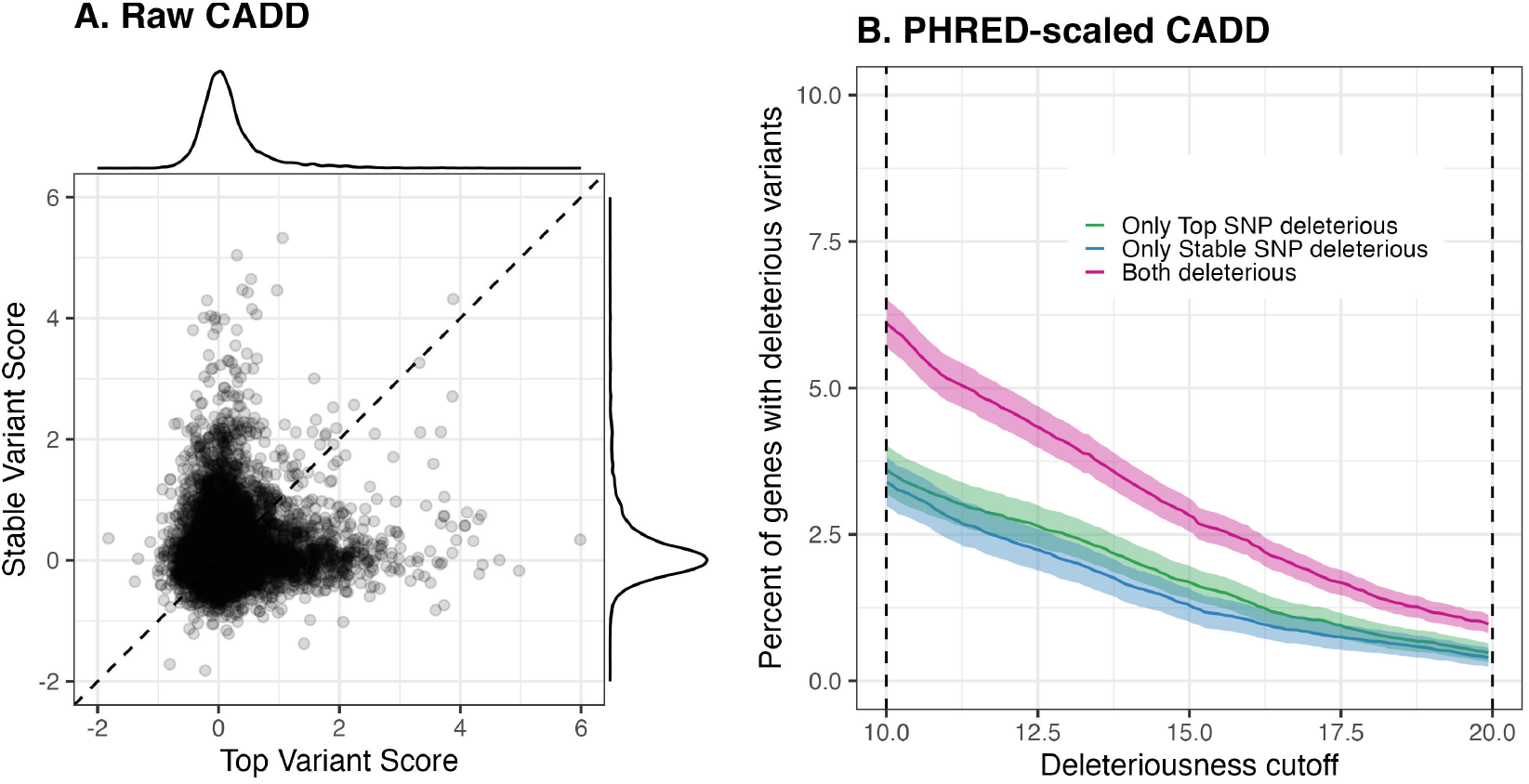
**A**. Paired scatterplot of raw CADD scores of both top and stable variant for each gene, for Potential Set 1. **B**. Percent of genes that are classified as (i) having deleterious top variant only, (ii) having deleterious stable variant only, and (iii) having both top and stable variant deleterious, using a sliding cutoff threshold ranging from 10 to 20 as recommended by CADD authors.

Because our comparison between the top and stable variants yielded no significant functional enrichment of one over the other, we investigate whether external factors — for example, the posterior probability of the variants — might moderate the relative enrichment of one variant over the other (i.e., we perform *trend analysis* — see Section for a list of all moderators). Here, we find that all but one of the moderators considered — namely, the Posterior Probability of Top Variant (see Table 2) — did not produce any significant trends for Potential Set 1. There is a small but significant positive correlation (Pearson’s *r* = 0.05) between the posterior probability of the top variant and the difference between the stable variant and top variant FIRE scores (***Ioannidis et al., 2017***), and a small but significant negative correlation (Pearson’s *r* = −0.07) between the posterior probability of the top variant and the difference between the stable variant and top variant Absolute Distance to Canonical TSS. (Details are reported in Supplementary File 1 Section S8.)

**Table 2.** List of 6 moderating factors considered.

| Moderator | Quantity/Statistic Computed |
| --- | --- |
| (1) Degree of Stability | No. subpopulations for which stable variant has positive probability |
| (2) Population Diversity | Maximum of pairwise allele frequency difference between subpopulations for which stable variant has positive posterior probability |
| (3) Population Differentiation | Maximum $F_{ST}$ between subpopulations for which stable variant has positive posterior probability |
| (4) Inclusion of Distal Subpopulations (Top) | Whether or not the top variant also had positive probability in Yoruban subpopulation when the stability-guided approach was used |
| (5) Inclusion of Distal Subpopulations (Stable) | Whether or not the stable variant had positive probability in Yoruban subpopulation when the stability-guided approach was used |
| (6) Degree of Certainty of Causality Using Residualization Approach | Posterior probability of top variant |

#### Additional comparisons

We perform various *conditional analyses* to evaluate whether additional restrictions to characteristics of fine-mapping outputs may boost the power of either approach over the other. Such characteristics include (i) the positive posterior probability support (i.e., how many variants reported a positive posterior probability from fine-mapping), and (ii) the posterior probability of the top or stable variant. Results from our conditional analysis of (i) are reported in Supplementary File 1 Section S9. As an example, for (ii) we further perform a comparison between top and stable variants, by restricting to genes where the posterior probability of the top variant or the stable variant exceeds 0.9. Such restriction of valid fine-mapped variant-gene pairs is useful in training variant effect prediction models requiring reliable positive and negative examples, as seen in ***Wang et al. (2021***). We find that for genes where the top variant posterior probability exceeds 0.9, there is no significant enrichment of the top variant over the stable variant across the 378 annotations considered (all BH-adjusted *p*-values exceed 0.05). Interestingly, when focusing on genes where the stable variant posterior probability exceeds 0.9, FIRE scores of the top variant are significantly larger than the stable variant (BH-adjusted *p*-value = 1 × 10^−6^). Detailed results are reported in Supplementary File 1 Section S10.

## Discussion

We have shown that a stability-guided approach complements existing approaches to detect biologically meaningful variants in genetic fine-mapping. Through various statistical comparisons, we have found that prioritizing the agreement between existing approaches and a stability-guided approach enhances the functional impact of the fine-mapped variant. Incorporating stability into fine-mapping also provides an adjuvant approach that helps discover variants of potential functional impact in case standard approaches fail to pick up variants of functional significance. Our findings are consistent with earlier reports of stable discoveries having the tendency to capture actual physical or mechanistic relationships, potentially making such discoveries generalizable or portable (***Basu et al., 2018***).

The link between stability and generalizability is not new. In the machine learning and statistics literature, it has been shown that stable algorithms provably lead to generalization errors that are well-controlled (***Bousquet and Elisseeff, 2002***, Theorem 17, p. 510). Furthermore, in certain classes of algorithms (e.g., sparse regression), it has been demonstrated empirically that this mathematical relationship is explained precisely by the stable algorithm removing spurious discoveries (***Lim and Yu, 2016***).

The stability-guided approach is also distinct from other subsampling approaches, such as the bootstrap (***Efron and Tibshirani, 1994***). First, the bootstrap is more often deployed as a method for calibrating uncertainty surrounding a prediction, which is not the objective of our method. Next, in settings where the bootstrap is deployed as a type of perturbation against which a prediction or estimand is expected to be stable (***Basu et al., 2018***), a stability threshold is implicitly needed and would require tuning to be chosen (e.g., in ***Basu et al., 2018*** this threshold was chosen to be 50%). Here, we leverage interpretable existing external annotations to define the perturbation against which we expect the fine-mapped variant to be stable — in other words, the user relates what is meant by the fine-mapped variant being stable to a biologically meaningful concept like “portable across environmentally and LD pattern-wise heterogeneous populations.”

The last sentence in the preceding paragraph suggests there are teleological similarities between the stability-guided approach and meta-analysis approaches (***Turley et al., 2021***). We emphasize that meta-analyses rely on already analyzed cohorts, thereby implicitly assuming that *within-cohort* heterogeneities have been sufficiently accounted for prior to the reporting of findings for that cohort. The stability-guided approach, however, is relevant to the *cohort-specific analysis itself*, where existing approaches may present methodological insufficiencies resulting in inflated false discoveries. In other words, whereas the goal of meta-analysis may be stated as identifying consistent hits across cohorts while also assuming that findings specific to each cohort are reliable, the goal of a stability-guided approach is to search for consistent signals despite the presence of potential confounders within a single cohort. Our analysis has focused on comparing the stability-guided approach against residualization, a convenient and popular approach to account for population structure, which reflects this difference in purpose.

Related to the previous point, while revising our work we came across a few recent papers that developed multi-ancestry fine-mapping algorithms to investigate causal variant heterogeneity across ancestries. ***Gao and Zhou (2024***) and ***Yuan et al. (2024***) found that complex traits have both shared and non-negligible ancestry-specific causal signals, although the proportion of trait-specific causal variants that are ancestry-specific is probably low (***Hu et al., 2025***; ***Shi et al., 2021***). On the other hand, for molecular quantitative traits, which include RNA sequencing reads that we have studied in this work, ***Lu et al. (2025***) reported highly correlated effect sizes in *cis* QTLs across ancestries, with effect heterogeneity concentrated at predicted loss-of-function-intolerant genes. These findings are all consistent with our goal of leveraging stability guidance to identify generalizable causal signals, and also imply that stability guidance would be particularly useful for molecular quantitative trait fine-mapping.

There are several limitations to our present work. First, we have chosen to focus on a non-parametric approach to fine-mapping, but multiple parametric fine-mapping approaches have been extended to incorporate cross-population heterogeneity (e.g., ***Wen et al., 2015***; ***LaPierre et al., 2021***; ***Lu et al., 2022***). While our present work is a proof-of-concept of the applicability of the stability to genetic fine-mapping, we believe that future work focusing on comparing functional impact of variants prioritized by our stability-driven approach or these parametric methods will shed more light on the efficacy of the stability principle at detecting generalizable biological signals. Our results for SuSiE on simulated gene expression data are promising in this regard. Second, our analyses assume that there is at most one *cis* eQTL prioritized by each potential set of PICS. This is owing to the implicit assumption that all other variants in high LD with the causal variant are simply tagging it, though we acknowledge the possibility of relaxing this assumption. Understanding the impact of this relaxation requires extensive investigation, hence we defer it to future work. Third, while we have relied on numerous computational and experimental annotations to evaluate the functional impact of our fine-mapped variants, some computational variant effect predictors may themselves be biased owing to the lack of ancestral diversity of training data. Recent work has found that deep learning-based gene expression predictors — despite outperforming traditional statistical models at inferring regulatory tracks — explain surprisingly little of gene expression variation across ancestries (***Huang et al., 2023***) and yield limited power at predicting individual gene expression levels more broadly (***Sasse et al., 2023***). While this complicates the use of such predictors in their present form as strong evidence of functional impact, it is more constructive to understand their shortcomings and develop strategies to fine-tune their predictions for future use as trustworthy functional impact metrics — an emerging desideratum in the era of computational biomedicine (***Katsonis et al., 2022***; ***Barbadilla-Martínez et al., 2025***; ***Benegas et al., 2025***; ***Livesey et al., 2025***).

In closing, while our work explores stability to subpopulation perturbations where subpopulations are defined by ancestry, we emphasize that our stability-guided slicing methodology is applicable to all settings where meaningful external labels are available to the data analyst. For instance, environmental or geographical variables, which are well-recognized determinants of some health outcomes (***Favé et al., 2018***; ***Abdellaoui et al., 2022***) and arguably better measure potential confounders than ancestry labels, can be the basis on which slices are defined in the stability-guided approach. As the barriers to access larger biobank-scale datasets, which contain such aforementioned variables, continue to be lowered, we expect stability-driven analyses conducted on such data and relying on carefully defined slices will help users better understand genetic drivers of complex traits. Given its utility in our present work, we believe that the stability approach in precision medicine may find uses beyond genetic fine-mapping and few other biological tasks previously studied, ultimately empowering the discovery of veridical effects not previously known.

## Materials and Methods

We use publicly available GEUVADIS B-lymphocyte RNA-seq measurements from 445 individuals (***Lappalainen et al., 2013***), whose genotypes are also available from the 1000 Genomes Project (1000 ***Genomes Project Consortium et al., 2015***). The 445 individuals come from five populations: Tuscany Italian (TSI, *N* = 91), Great British (GBR, *N* = 86), Finnish (FIN, *N* = 92), Utah White American (CEU, *N* = 89) and Yoruban (YRI, *N* = 87). The mRNA measurements are normalized for library depth, expression frequency across individuals as well as PEER factors, as reported in the GEU-VADIS databank^1^. Of the available normalized gene expression phenotypes, only those with non-empty sets of variants lying within 1Mb upstream or downstream of the canonical transcription start site were kept for (*cis*) fine-mapping. This process yielded 22, 664 genes used in all subsequent analyses involving the PICS fine-mapping algorithm.

### Probabilistic Identification of Causal SNPs

We implement Probabilistic Identification of Causal SNPs (PICS) (***Farh et al., 2015***), which is based on an earlier genome-wide analysis identifying genetic and epigenetic maps of causal autoimmune disease variants (***Farh et al., 2015***).

We assume that **X** and **y** are the *N* × *P* locus haplotype (or genotype) matrix and the *N* × 1 vector of phenotype values respectively, with **y** possibly already adjusted by relevant covariates. For consistency of exposition, let the SNPs be named *A*_1_,…, *A*_*P*_. Recall the goal of fine-mapping is to return information about which SNP(s) in the set {*A*_1_,…, *A*_*P*_} is (are) causal, given the input pair {**X, y**} and possibly other external information such as functional annotations or, more generally, prior biological knowledge.

#### Overview

Probabilistic Identification of Causal SNPs (PICS) is a Bayesian, non-parametric approach to fine-mapping. Given the observed patterns of association at a locus, and furthermore not assuming a parametric model relating the causal variants to the trait itself, one can estimate the probability that any SNP is causal by performing permutations that preserve its marginal association with the trait as well as the LD patterns at the locus.

To see how this is accomplished, suppose that *A*_1_ is the lead SNP. We are interested in 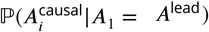, the probability that *A*_*i*_is causal, for each *i ∈* [*P* ]. By Bayes’ theorem,

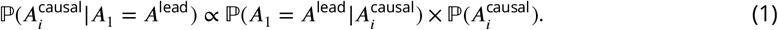

Focusing on the focal SNP *i*, permute the rows of **X** such that the association between the focal SNP and the trait **y** is invariant; see Supplementary File 1 Section S1 for a concrete mathematical description of this set of constrained permutations. Then the first term on RHS of eq. (1), 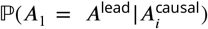, is estimated by the proportion of all permutations where *A*_1_ emerges as the lead SNP. The second term, 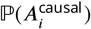, is the prior probability of the focal SNP being causal, which the user can choose based on prior knowledge. The default setting in PICS is 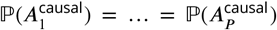. Figure 6 provides a visual summary of the method.

**Figure 6.**
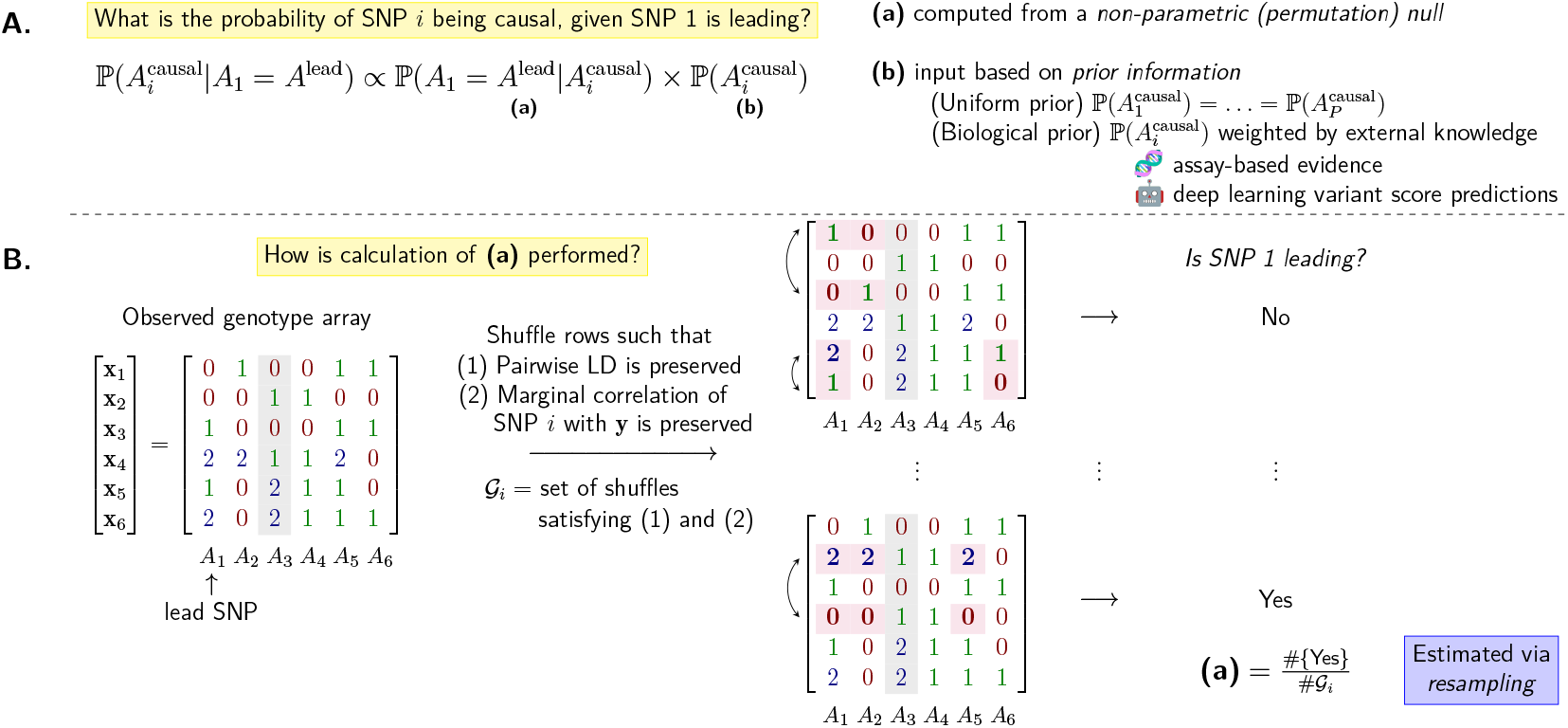
Visual summary of the PICS algorithm described in Probabilistic Identification of Causal SNPs. **A**. Breakdown of the calculation of the probability of a focal SNP *A*_*i*_ being causal. **B**. Illustration of the permutation procedure used to generate the null distribution. An example *N* × *P* genotype array with *N* = *P* = 6 is used, with two valid row shuffles, or permutations, of the original array shown. Entries affected by the shuffle are highlighted, as is the focal SNP (*A*_3_).

By running the permutation procedure across all SNPs in the locus, one obtains 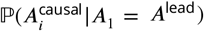 _for each *i*. These posterior probabilities are then normalized so that 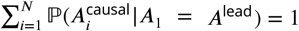._

Finally, the above procedure is performed after restricting the set of all SNPs to only those with correlation magnitude |*r*| *>* 0.5 to the lead SNP.

#### Algorithm

Based on the computation of posterior probabilities described above (eq. (1)), the full PICS algorithm returns putatively causal variants as follows.

The last line of Algorithm 1 returns a list of posterior probability vectors corresponding to each potential set. For our work, we set |*r*| = 0.5, *C* = 3, *R* = 500 and **p**_0_ = (1/*P*, …, 1/*P*) throughout implementations of Algorithm 1.

##### Algorithm 1 PICS

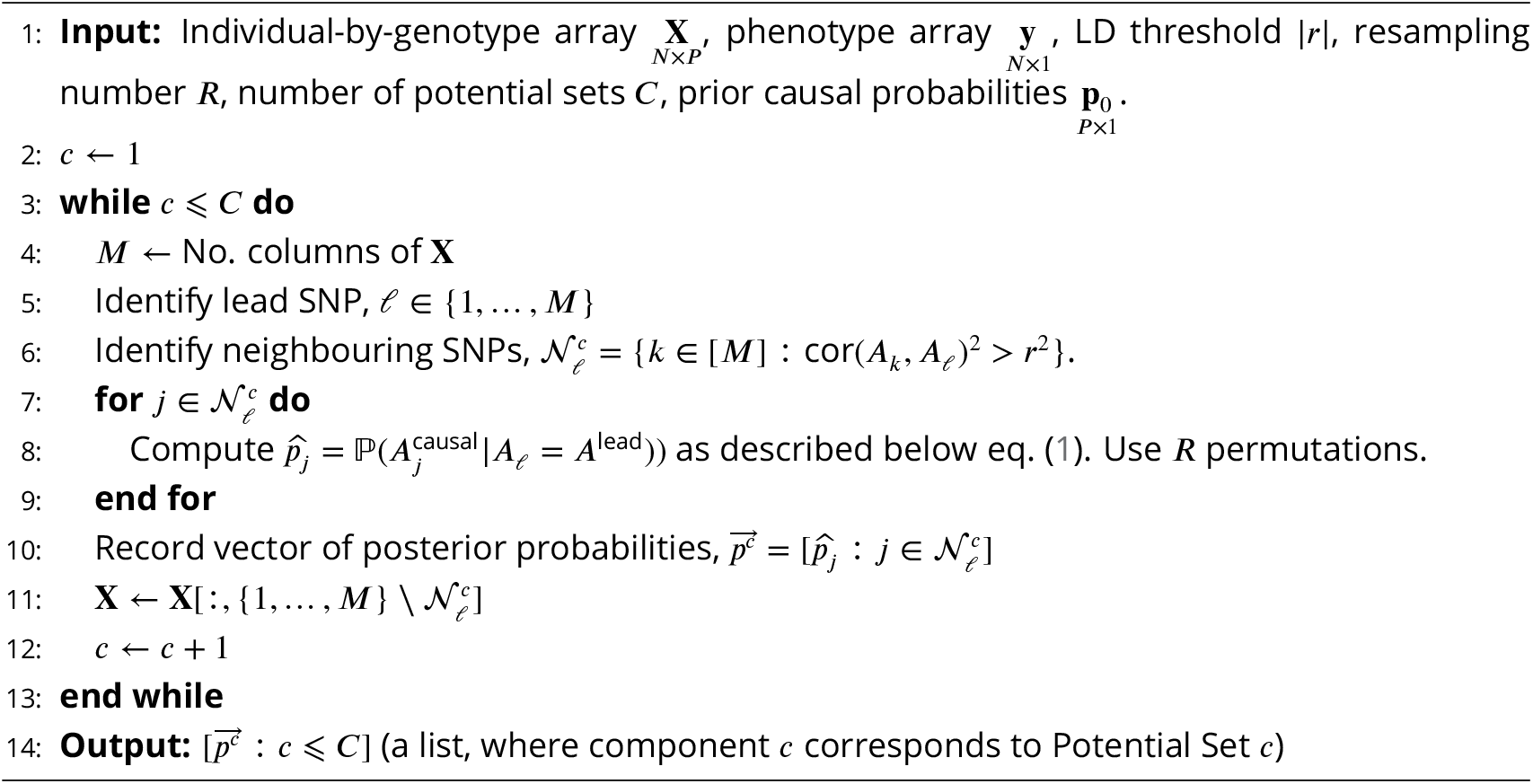

As mentioned in the Introduction, we apply two different approaches to implementing Algorithm 1. One is guided by stability and will be introduced shortly in Incorporating Stability. The other, which we now describe, is based on regressing out confounders, i.e., residualization. In implementing the residualization approach, we regress the top five principal components, obtained from the genotype matrix **X**, from the gene expression phenotype, **y**. We choose five principal components based on the elbow method, an approach described in ***Brown et al. (2018***). This yields residuals **y**^*r*^(Figure 1A), which we use as the input phenotype array in Algorithm 1. We subsequently report the variant with highest posterior probability in each potential set as the putatively causal variant for that potential set. This variant is referred to as the top variant.

We remark that there are other ways of using potential confounders in variable selection, such as inclusion as covariates in a linear model. Because our algorithm explicitly avoids assuming a linear model, we have chosen the residualization approach just described.

### Incorporating Stability

Our stability-guided approach to implementing PICS follows the steps outlined below. Let (**X, y**) be the genotype array and gene expression array. Assume that there are *K* subpopulations making up the dataset, and let *E*_*k*_ denote the set of row indices of **X** corresponding to individuals from subpopulation *k*. (In our present work, *K* = 5. The five subpopulations are Utahns (*N*_CEU_ = 89), Finns (*N*_FIN_ = 92), British (*N*_GBR_ = 86), Toscani (*N*_TSI_ = 91) and Yoruban (*N*_YRI_ = 87).)

1. Run Algorithm 1 on the pair (**X, y**). Obtain list of posterior probability vectors, 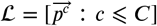. Recall that *C* is the number of potential sets.
2. For each subpopulation *k* = 1,…, *K*, run Algorithm 1 on the pair 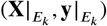. Obtain 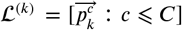 for each *k*.
3. (Stability-guided choice of putatively causal variants) Collect the probability vectors in lists ℒ and 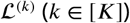. Operationalizing the principle that a stable variant has positive probability across multiple slices, we pick causal variants as follows. For potential set *c*, pick the variants that have (i) positive probability in 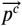 in Step 1, and moreover (ii) have positive probability in the most number of probability vectors; call this set of variants 𝒮_*c*_(note 𝒮_*c*_is a subset of the support of 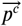 for Potential Set *c*). Among members of 𝒮_*c*_, select the variant that had the highest posterior probability in 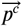.

In other words, the stability-guided approach reports the variant that not only appears with positive probability in the most number of subsets including the pooled sample, but also has the largest probability in the pooled sample. This variant is referred to as the stable variant.

To illustrate the last step, suppose there are *K* = 2 subpopulations, *C* = 1 potential set, and *P* = 5 SNPs. Let the outputs from Steps 1 and 2 be

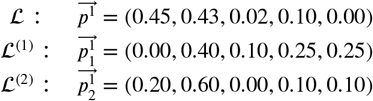

In this example, the second and fourth variants have positive probability in the most number of probability vectors (𝒮_1_ = {*A*_2_, *A*_4_}). Among *A*_2_ and *A*_4_, *A*_2_ has a higher posterior probability (i.e., 0.43) in 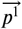, so the stable variant reported is *A*_2_. As a side remark, the posterior probability vector ℒ does not generally agree with the posterior probability vector computed using the residualization approach, because the latter is computed by running Algorithm 1 on the pair (**X, y**^*r*^) rather than (**X, y**), as described earlier.

#### Stability-guided SuSiE

We apply the stability principle to SuSiE (***Wang et al., 2020***) in a similar manner as described above. Concretely, we run SuSiE on the pair (**X, y**), before running it again on each subpopulation *k* = 1,…, *K*. Collecting the posterior inclusion probability (PIP) vectors across each subpopulation and the entire sample, we then keep only variants that have posterior probability at least 1/(no. variants included), owing to SuSiE tending to return nonzero probabilities to all variants included in the fine-mapping. These “good” variants (i.e., variants with PIP at least as large as the prior inclusion probability) are analogous to the “variants with positive probability” in PICS. The remaining steps are identical to stability-guided PICS.

### Simulation Study and Evaluation Details

#### Main Simulation Study

We simulate 2, 400 synthetic gene expression data, which vary by two parameters: *C*, the number of causal variants, and *ϕ*, the proportion of variance in **y** explained by the genotypes **X**. Phenotypes are simulated as follows.

1. For a gene, whenever its canonical transcription start site (TSS) is available, restrict the genotype matrix **X** to only variants lying within 1Mb upstream and downstream. Else, use variants spanning the entire autosome associated with that gene. This ensures that *cis* eQTLs are used in simulation, for the most part.
2. Sample the indices of the *C* causal variants, 𝒞, uniformly at random from {1, …, *p*}, where *p* is the number of variants included in **X** from Step 1.
3. For each *j ∈* 𝒞, independently draw *b*_*j*_ ∼ *N*(0, 0.6^2^) and, for all *j* ∉ 𝒞, set *b*_*j*_ = 0.
4. Set *σ*^2^to achieve the desired proportion of variance explained *ϕ*.
5. Generate the phenotype by drawing **y** ∼ *N*(**Xb**, *σ*^2^**I**_*n*×*n*_).

After gene expression data were simulated, we ran three versions of PICS: the stability-guided version, the residualization version, and an uncorrected version, where no correction for population stratification or stability guidance was applied. To investigate the broader applicability of the stability approach, we also ran two versions of SuSiE (***Wang et al., 2020***), a popular fine-mapping method relying on variational approximation techniques. The first version runs SuSiE on residualized phenotypes, while the second version incorporates stability in selecting the most likely causal variant(s). In both approaches, we rely on default parameters of SuSiE while setting the number of credible sets to return to 3 (i.e., `L=3` in `susieR::susie`).

#### Smaller Simulation Study

To investigate whether non-genetic confounding between ancestries in a multi-ancestry cohort can hinder the performance of fine-mapping algorithms that do not correct for potential confounding by ancestry, we simulate a smaller set of synthetic gene expression data. We select 10 genes from Chromosomes 1, 20, 21 and 22, and repeated the simulation steps as described above, up to Step 4. At Step 4, instead of choosing a single *σ*^2^, we allow *σ*^2^to depend on the ancestry of the individual in order to capture environmental heterogeneity. We specifically consider six scenarios that we classify as (environmental) variance heterogeneity and mean heterogeneity.

- *Variance heterogeneity*. Let ***α***(*t*) = (*α*_TSI_(*t*), *α*_GBR_(*t*), *α*_FIN_(*t*), *α*_CEU_(*t*), *α*_YRI_(*t*)) be a vector of exponentiated proportions, where we define the exponentiated proportion of a subpopulation POP (POP *∈* {TSI, FIN, GBR, CEU, YRI})as 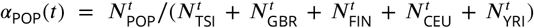. In Step 4, we define a vector of ancestry-specific exogenous variances, ***σ***^**2**^(*t*) = 5*σ*^2^***α***(*t*), where *σ*^2^is calculated as in Step 4 of the original simulation setup. We then generate phenotypes independently in Step 5 by conditioning on the individual’s population membership (POP): *y*_*i*_ ∼ *N*(***x***_*i*_**b, *σ***^**2**^(*t*)(*k*)), where *k* is the index that corresponds to the population membership of individual *i* and ***x***_*i*_ denotes the genotypes of individual *i*. For example, if *t* = 0, ***α***(*t*) = ***α***(0) = (1/5, 1/5, 1/5, 1/5, 1/5) and ***σ***^**2**^(*t*) = (*σ*^2^, *σ*^2^, *σ*^2^, *σ*^2^, *σ*^2^), which reduces to our original simulation setup. The choice of *t >* 0 determines the heterogeneity of the exogenous variance among the subpopulations, with a larger *t* producing higher heterogeneity. Here, we consider four choices of *t*: 8, 16, 128, and 256.
- *Mean heterogeneity*. Step 5 of the original simulation setup is modified to include an ancestry-dependent mean vector. We arbitrarily let GBR be the “focal” population, and generate phenotypes in two ways.
  – “Smooth mean”: After *σ*^2^is calculated in Step 4 of the original simulation setup, define the mean exogenous noise vector as ***μ*** = (*μ*_TSI_, *μ*_GBR_, *μ*_FIN_, *μ*_CEU_, *μ*_YRI_) = (2*σ*, 0, *σ, σ*, 2*σ*). An individual *i* has their phenotype simulated as *y*_*i*_ ∼ *N*(***x***_*i*_**b** + ***μ***(*k*), *σ*^2^), where again *k* is the index that corresponds to the population membership of individual *i*. This effectively increases the gene expression of all but GBR individuals in a positive direction with varying amounts of shift.
  – “Spiked mean”: After *σ*^2^is calculated in Step 4 of the original simulation setup, define ***μ*** = (0, 2*σ*, 0, 0, 0) and simulate individual *i*’s phenotype as *y*_*i*_ ∼ *N*(***x***_*i*_**b** + ***μ***(*k*), *σ*^2^). This effectively increases the gene expression of GBR individuals only in a positive direction, while the other subpopulation individuals share the same zero mean (as in the original simulation setup).

For brevity, we refer to these six scenarios by t=8, t=16, t=128, t=256, |i-3| and i=3.

#### Evaluation

Several metrics have been used to evaluate Bayesian fine-mapping algorithms (see e.g., ***Cui et al., 2024***; ***Yang et al., 2023***; ***Wang et al., 2020***; ***Benner et al., 2016***; ***Wen et al., 2016***; ***Hormozdiari et al., 2015***), many of which compute credible sets and analyze their coverage. A challenge in using credible sets for our study is that the posterior probabilities returned by the PICS algorithm are not posterior inclusion probabilities in a strict sense: they do not measure the frequency with which a variant should be included in a *linear model* (cf. ***Griffin and Steel, 2021***; note PICS does not assume a model relating features to outcome). This complicates defining credible sets and coverage, so we follow ***Mazumder (2020***) instead and evaluate performance of all algorithms by computing the signal recovery probability as a function of the signal to background noise (SNR). By the definition of *ϕ* (see ***Wang et al., 2020*** for details), we may compute SNR directly from *ϕ* as SNR = *ϕ*/(1 − *ϕ*).

### Statistical Comparison Methodology

We rely on 378 external functional annotations to interpret biological significance of our variants, summarized in Table 1. Supplementary File 1 Section S5 provides interpretation for the functional annotations, while Supplementary File 1 Section S6 describes in greater detail how we generate annotations from Enformer predictions.

Our comparison of the top and stable variants is two-fold. First, we evaluate the relative significance of the stable variant against the top variant, by running paired Wilcoxon two-sample tests across all pairs of top and stable variants across all GEUVADIS gene expression phenotypes. We compute one-directional *p*-values in either direction to check for significant depletion or enrichment of the stable variant with respect to a particular annotation. Raw *p*-values are adjusted for false discovery rate control by applying the standard Benjamini-Hochberg (BH) procedure (`R command p.adjust(…,method=‘BH’)`) to *p*-values across all 378 annotations and all potential sets.

To investigate whether various external factors might moderate the differences in functional annotations, we next perform trend tests. Basically, for some external factor *F*, we run a trend test to see if values of *F* are associated with attenuation or augmentation of differences in functional annotation of the top and stable variants. The list of all external factors *F* is provided in Table 2.

For (1) Degree of Stability, we perform a Jonckheere-Terpstra test (R command `clinfun:: jonckheere.test(…,nperm=5000))`. For (2) Population Diversity, (3) Population Differentiation and (6) Degree of Certainty of Causality Using Residualization Approach, we perform a correlation test (R command `cor.test(…))`. For (4) Inclusion of Distal Subpopulations (Top) and (5) Inclusion of Distal Subpopulations (Stable), we perform an unpaired Wilcoxon test (R command `wilcox.test(…))`. Finally, for each moderator, we again compute one-directional *p*-values in either direction, before applying the BH procedure to all *p*-values across annotations and potential sets for that moderator only.

## Data Availability

We provide the following data and scripts on the Github repository https://github.com/songlab-cal/StableFM: fine-mapped variants with their functional annotations and moderator variable quantities, code for reproducing Main Text figures in this manuscript and building our visualization app.

## Author Contributions

AJA, NMI, and YSS designed the study. AJA performed data analyses and developed the interactive visualization app. LCJ developed the interactive visualization app. NMI and YSS supervised the research.

## Acknowledgments

This research is supported in part by an NIH grant R35-GM134922 and grant number CZF2019-002449 from the Chan Zuckerberg Initiative Foundation. We thank Carlos Albors, Gonzalo Benegas, Ryan Chung, Ziyue Gao, Iain Mathieson and Yutong Wang for helpful discussions; members of the Yu Group at Berkeley for feedback on the work; and Aniketh Reddy for help with data processing.

---

**Figure 2—figure supplement 1.**
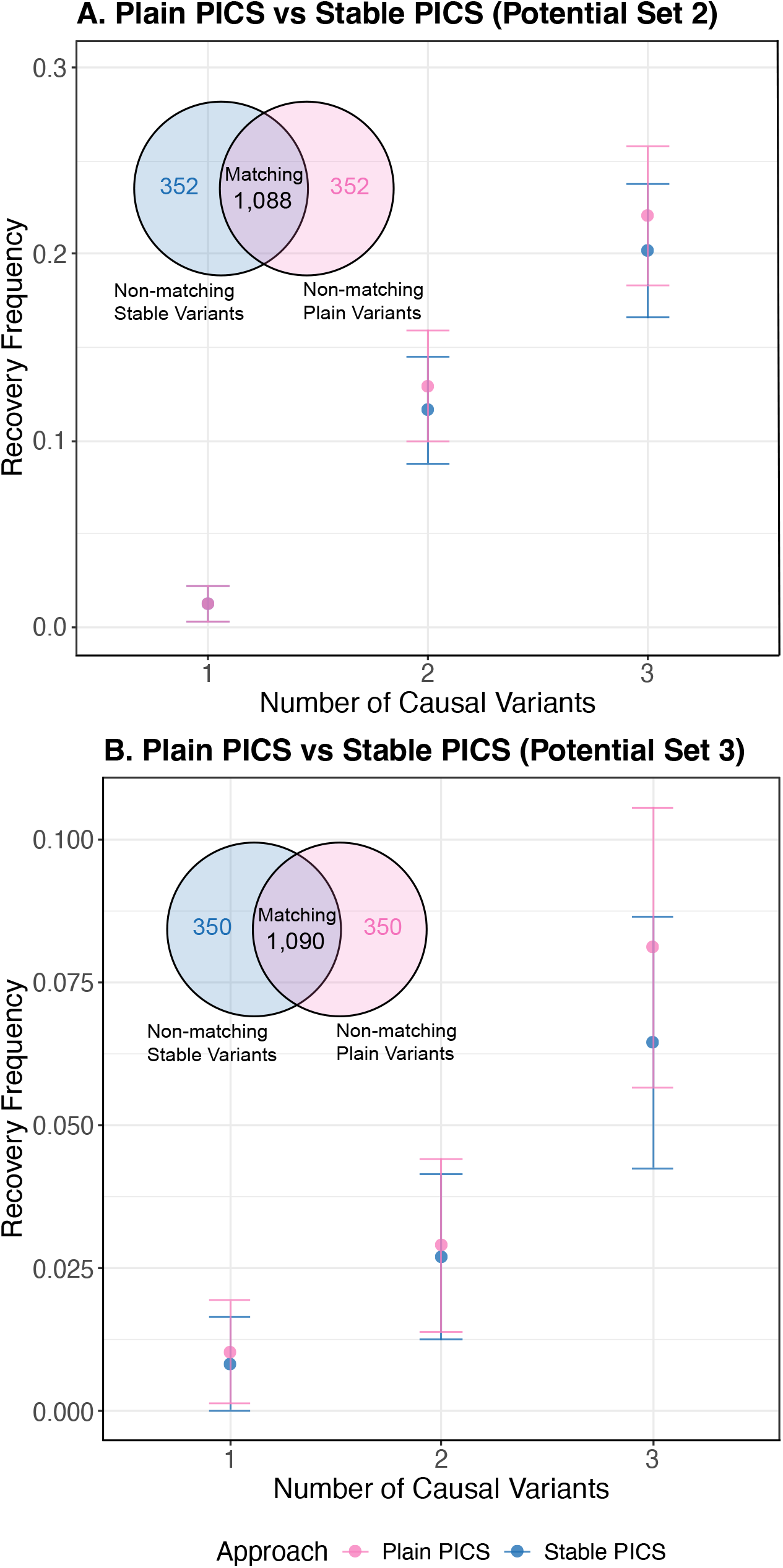
Frequency with which at least one causal variant is recovered in Potential Sets 2 and 3 by Plain PICS and Stable PICS, across 1, 440 simulated gene expression phenotypes. Recovery frequencies are stratified by simulations differing in the number of causal variants, but Venn diagrams report the number of matching and non-matching variants across all simulations.

**Figure 2—figure supplement 2.**
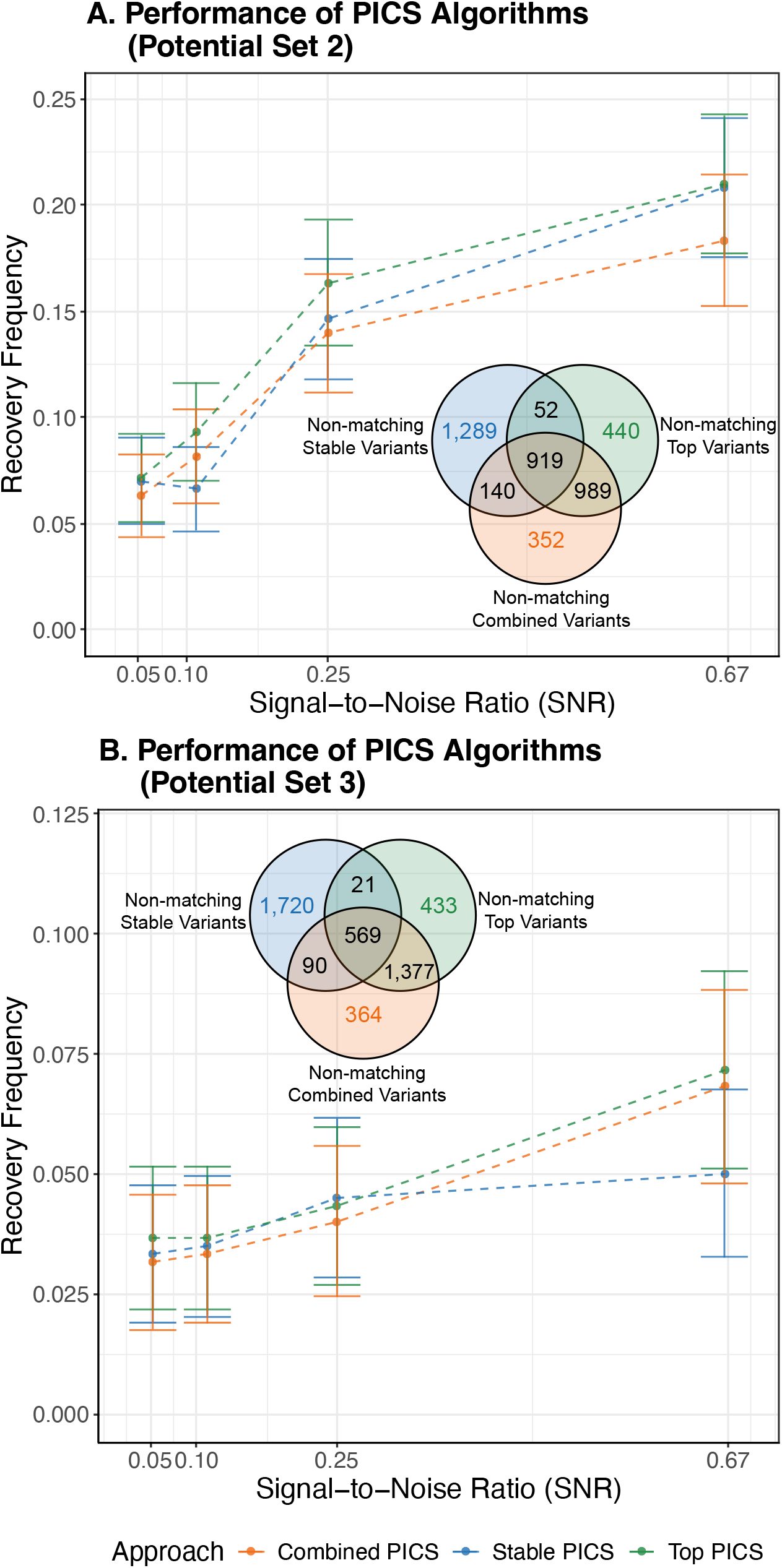
Frequency with which at least one causal variant is recovered in Potential Sets 2 and 3 by Combined PICS, Stable PICS and Top PICS, across 2, 400 simulated gene expression phenotypes. Recovery frequencies are stratified by simulations differing in the signal-to-noise ratio (SNR) parameter *ϕ*, but Venn diagrams report the number of matching and non-matching variants across all simulations.

**Figure 2—figure supplement 3.**
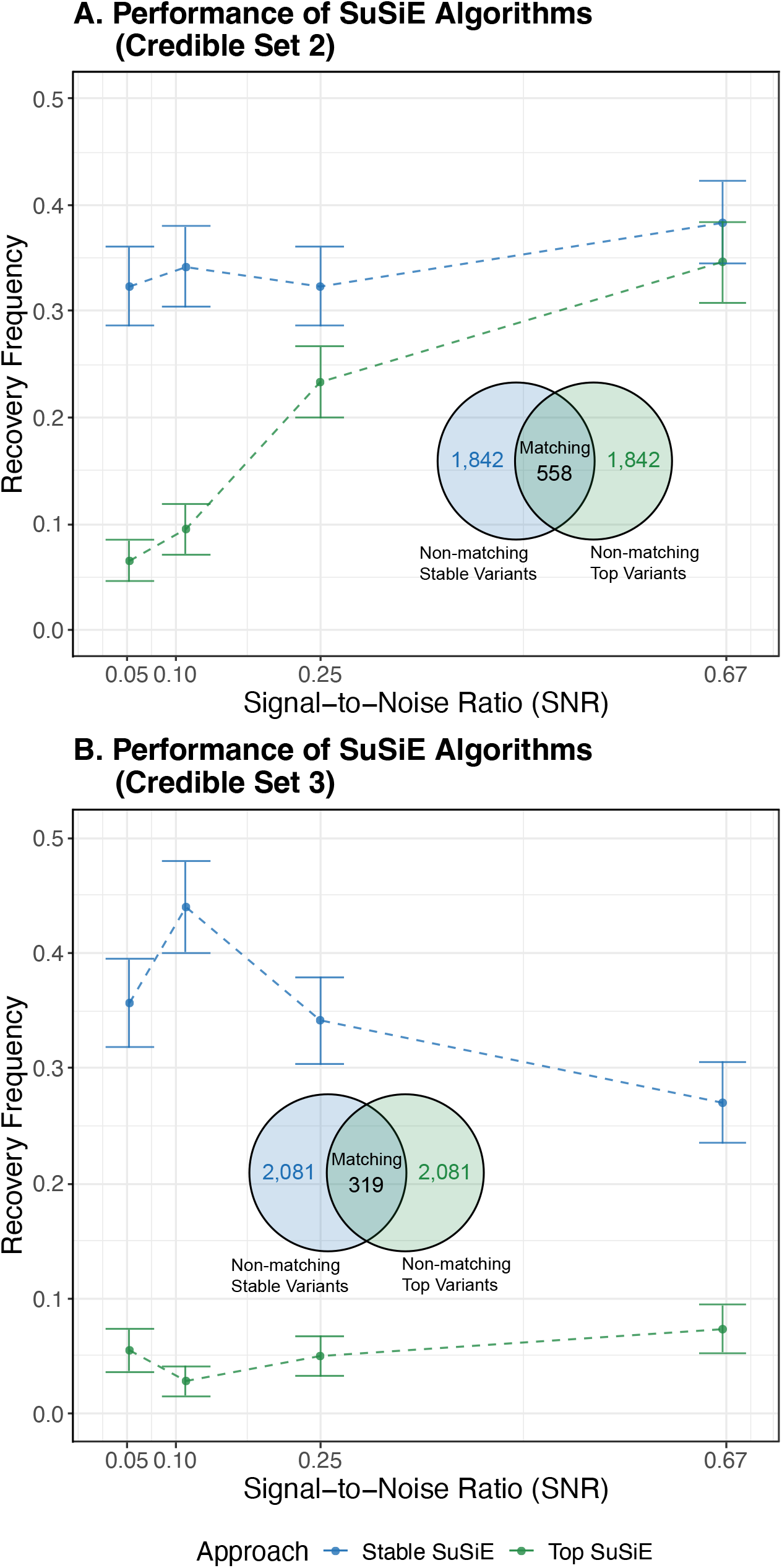
Frequency with which at least one causal variant is recovered in Potential Sets 2 and 3 by Stable SuSiE and Top SuSiE, across 2, 400 simulated gene expression phenotypes. Recovery frequencies are stratified by simulations differing in the signal-to-noise ratio (SNR) parameter *ϕ*, but Venn diagrams report the number of matching and non-matching variants across all simulations.

**Figure 2—figure supplement 4.**
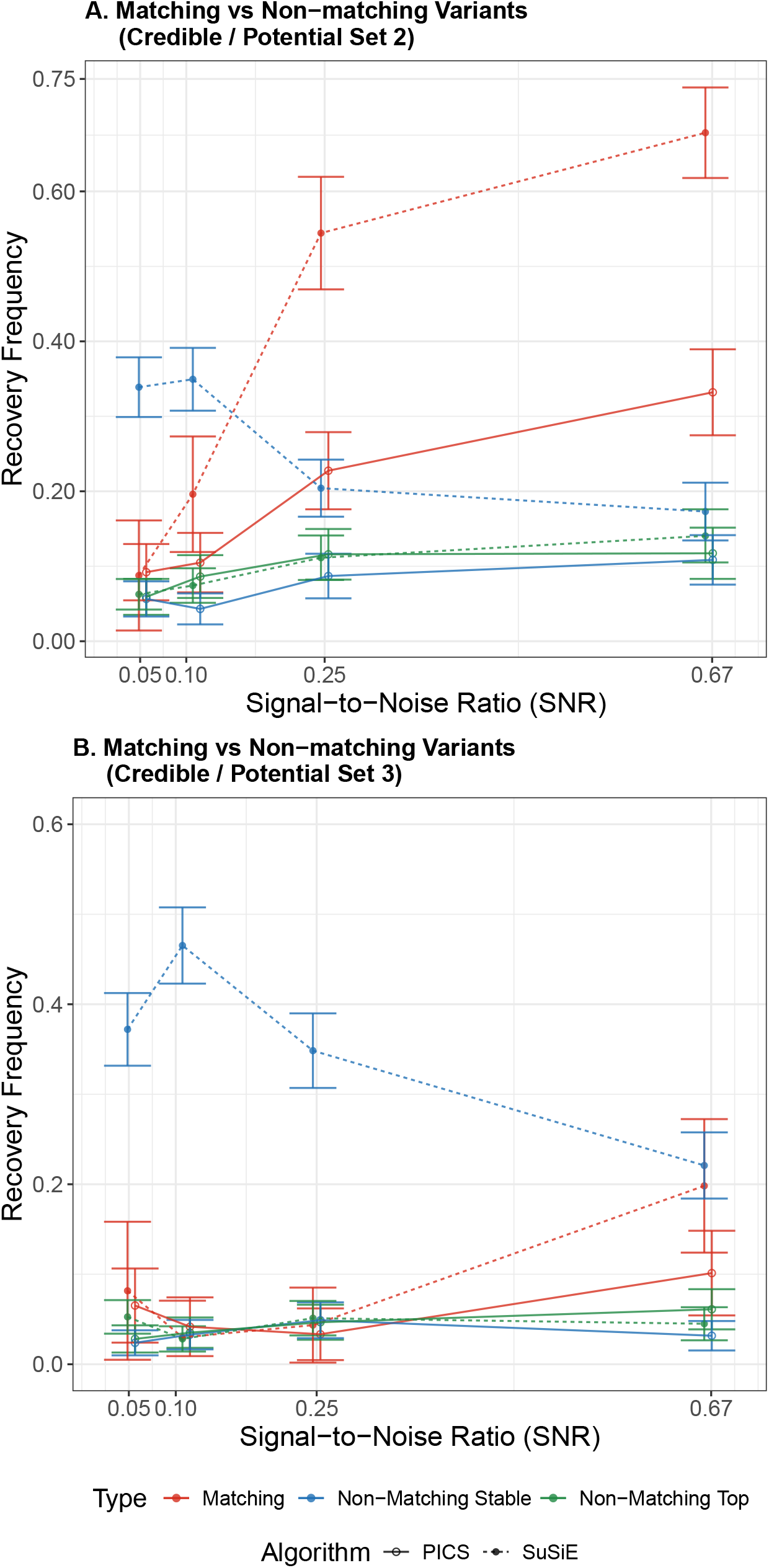
Frequencies with which matching and non-matching variants in the credible or potential set recover a causal variant, obtained from comparing top and stable approaches to an algorithm. Analysis is performed over 2, 400 simulated gene expression phenotypes, and recovery frequencies are stratified by simulations differing in the signal-to-noise ratio (SNR) parameter *ϕ*. **A**. Credible or Potential Set 2. **B**. Credible or Potential Set 3.

**Figure 2—figure supplement 5.**
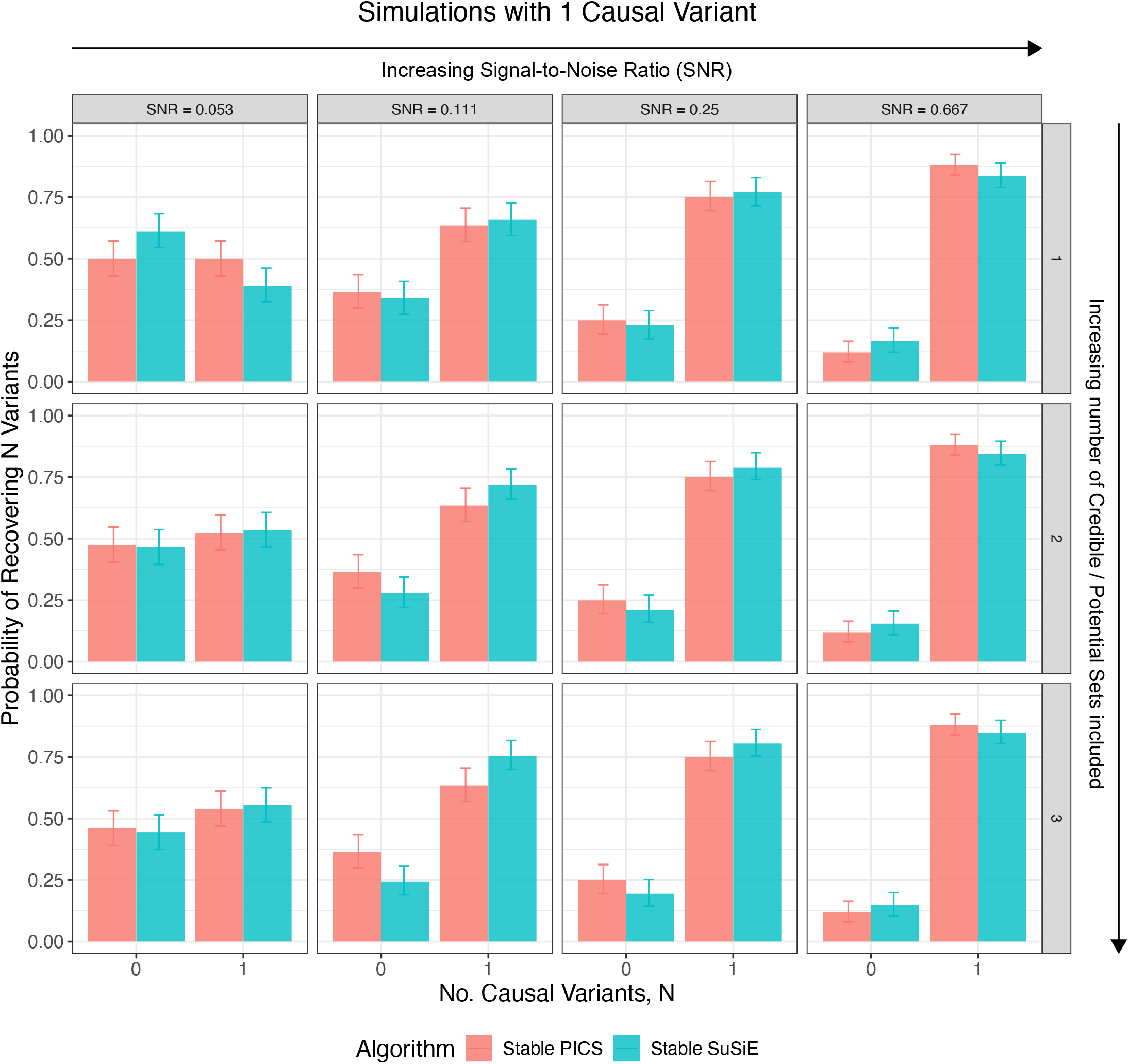
Empirical discrete probability distributions over the number of causal variants (0 or 1) recovered by stability-guided algorithms in simulations with one causal variant, stratified by the SNR parameter used in simulations (increasing SNR from left to right). The impact of including more number of credible or potential sets on the distribution is shown (increasing number of included sets from top to bottom).

**Figure 2—figure supplement 6.**
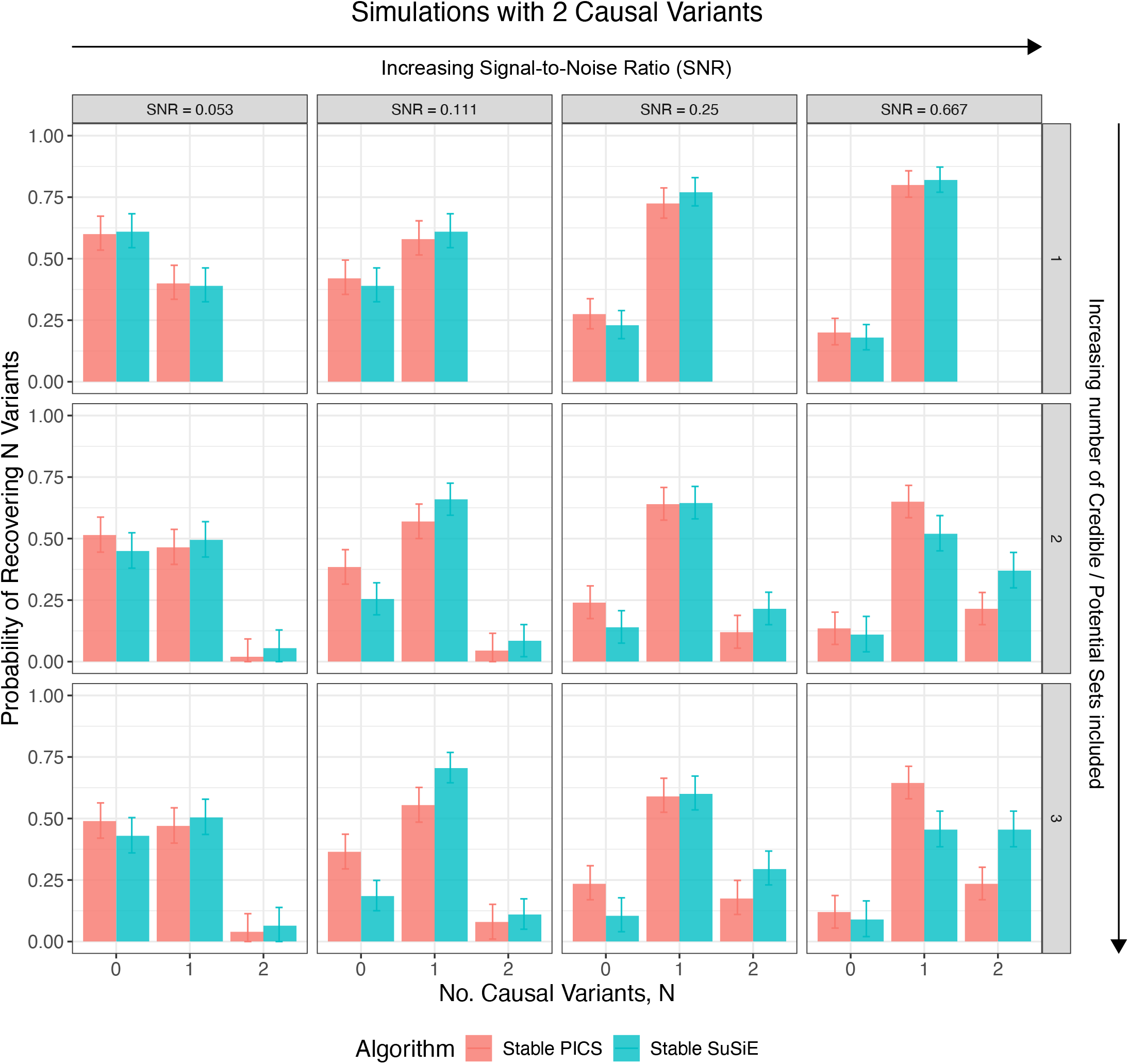
Empirical discrete probability distributions over the number of causal variants (0, 1 or 2) recovered by stability-guided algorithms in simulations with two causal variants, stratified by the SNR parameter used in simulations (increasing SNR from left to right). The impact of including more number of credible or potential sets on the distribution is shown (increasing number of included sets from top to bottom).

**Figure 2—figure supplement 7.**
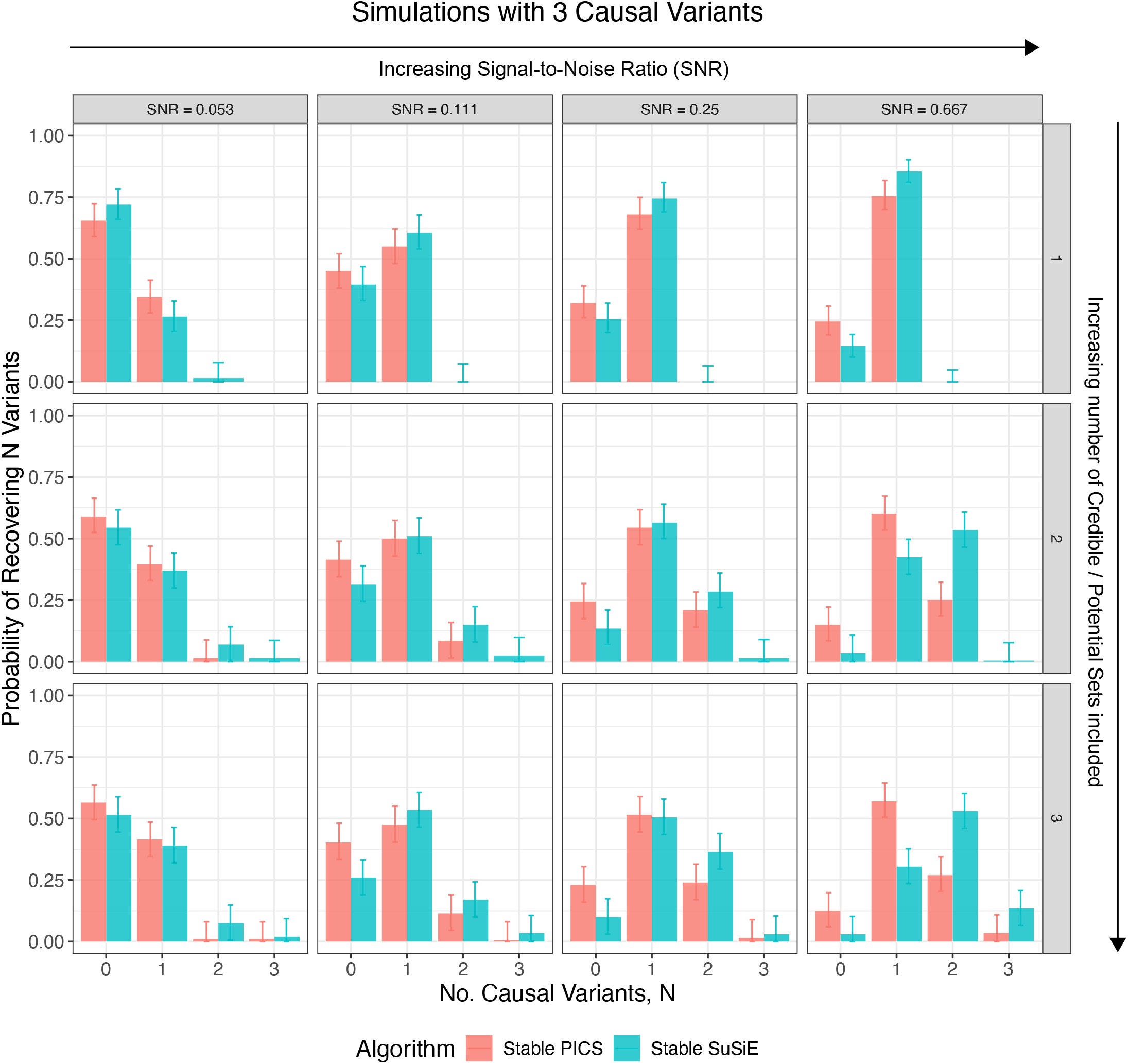
Empirical discrete probability distributions over the number of causal variants (0, 1, 2 or 3) recovered by stability-guided algorithms in simulations with three causal variants, stratified by the SNR parameter used in simulations (increasing SNR from left to right). The impact of including more number of credible or potential sets on the distribution is shown (increasing number of included sets from top to bottom).

**Figure 2—figure supplement 8.**
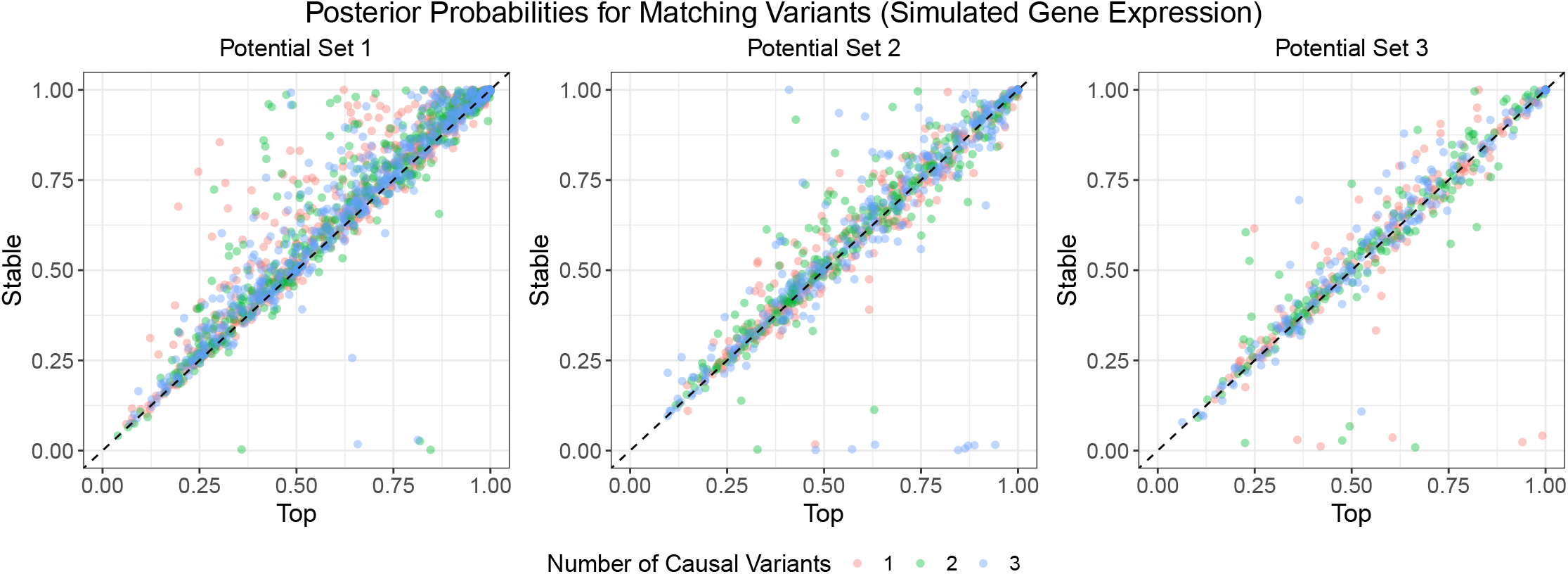
Posterior probabilities of matching top and stable variants across 2, 400 simulated gene expression phenotypes. Points are coloured by the number of causal variants, *S ∈* {1, 2, 3}, set in simulations.

**Figure 2—figure supplement 9.**
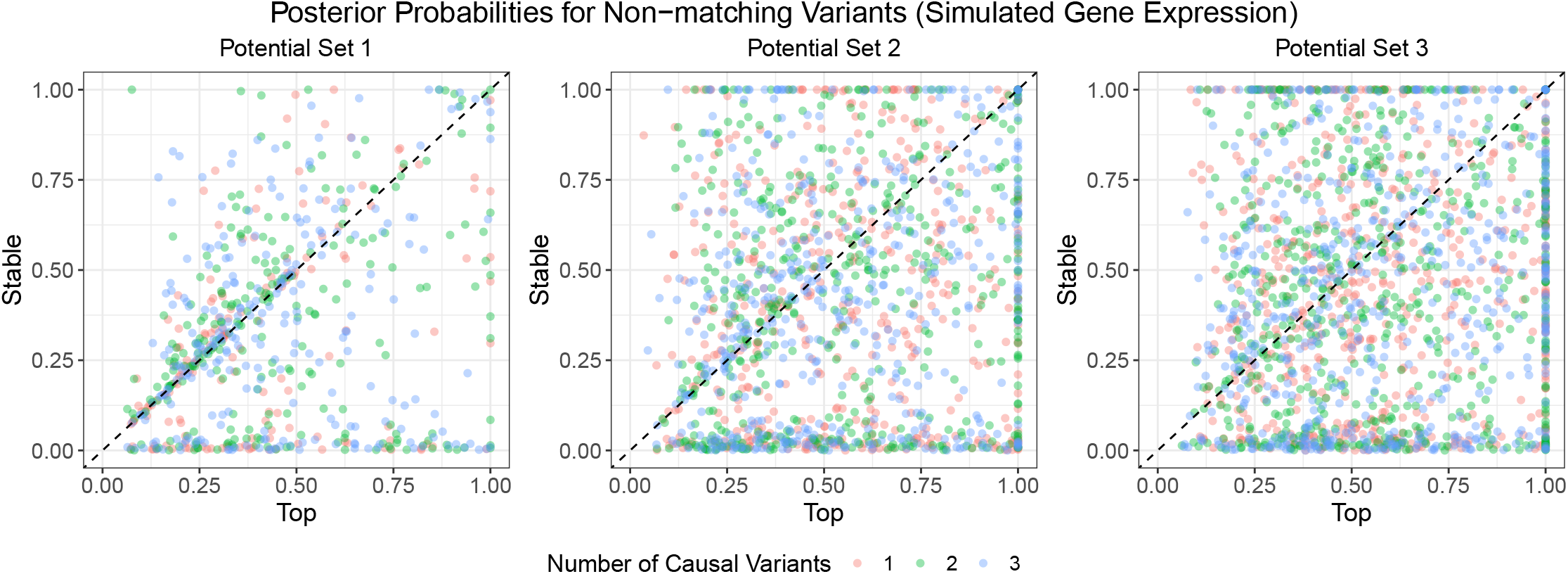
Posterior probabilities of non-matching top and stable variants across 2, 400 simulated gene expression phenotypes. Points are coloured by the number of causal variants, *S ∈* {1, 2, 3}, set in simulations.

**Figure 2—figure supplement 10.**
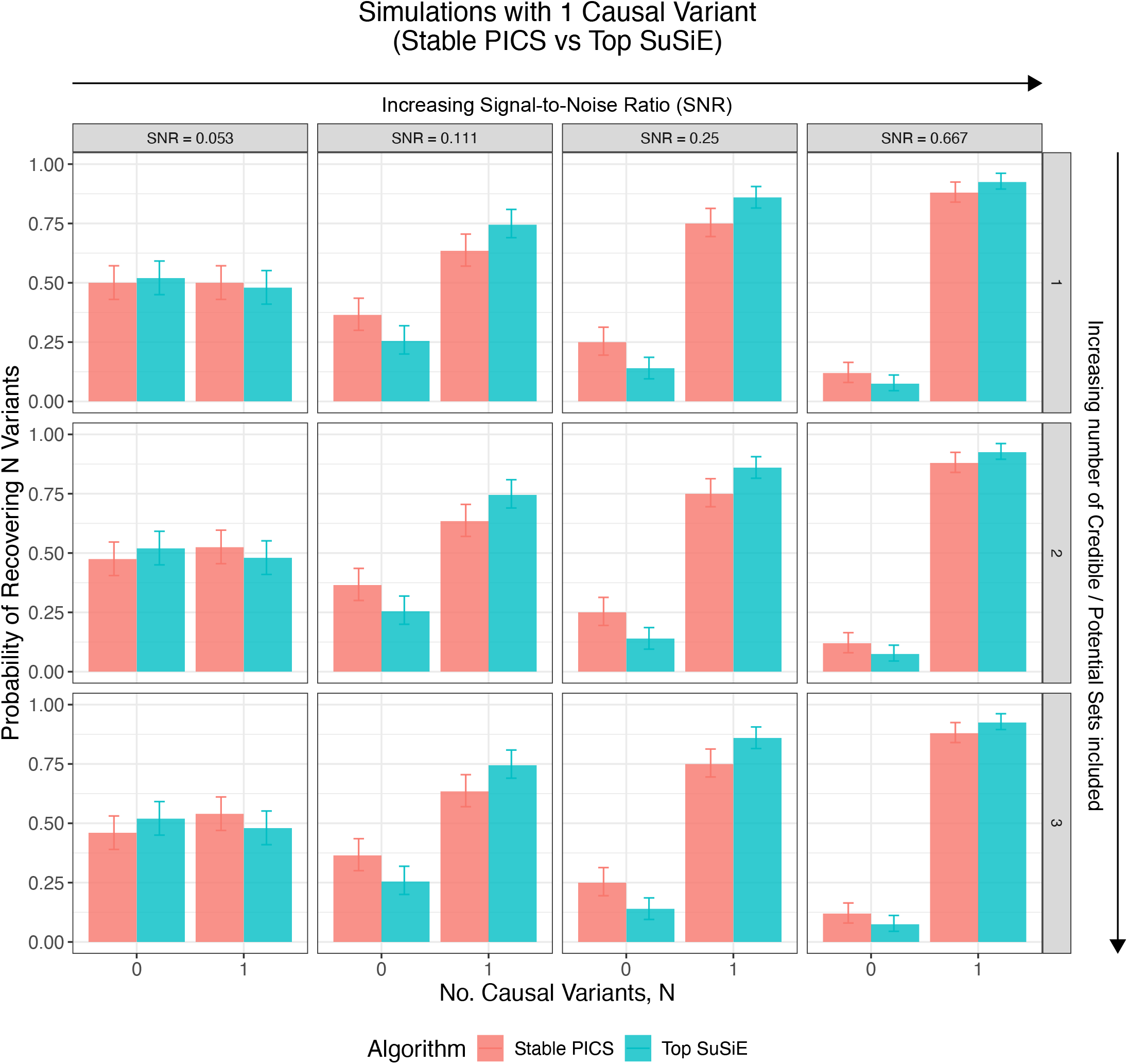
Empirical discrete probability distributions over the number of causal variants (0 or 1) recovered by Stable PICS or Top SuSiE in simulations with one causal variant, stratified by the SNR parameter used in simulations (increasing SNR from left to right). The impact of including more number of credible or potential sets on the distribution is shown (increasing number of included sets from top to bottom).

**Figure 2—figure supplement 11.**
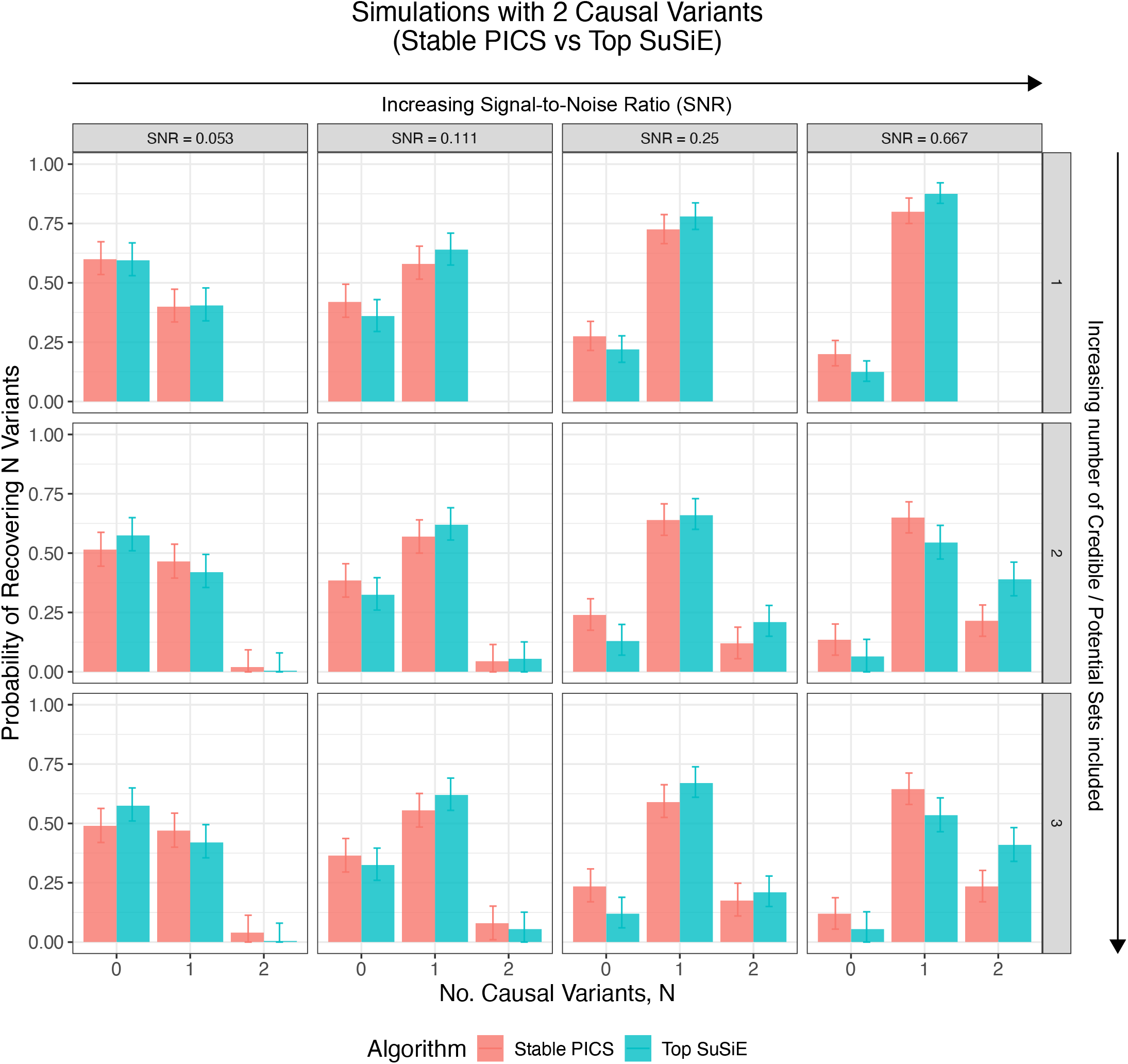
Empirical discrete probability distributions over the number of causal variants (0, 1 or 2) recovered by Stable PICS or Top SuSiE in simulations with two causal variants, stratified by the SNR parameter used in simulations (increasing SNR from left to right). The impact of including more number of credible or potential sets on the distribution is shown (increasing number of included sets from top to bottom).

**Figure 2—figure supplement 12.**
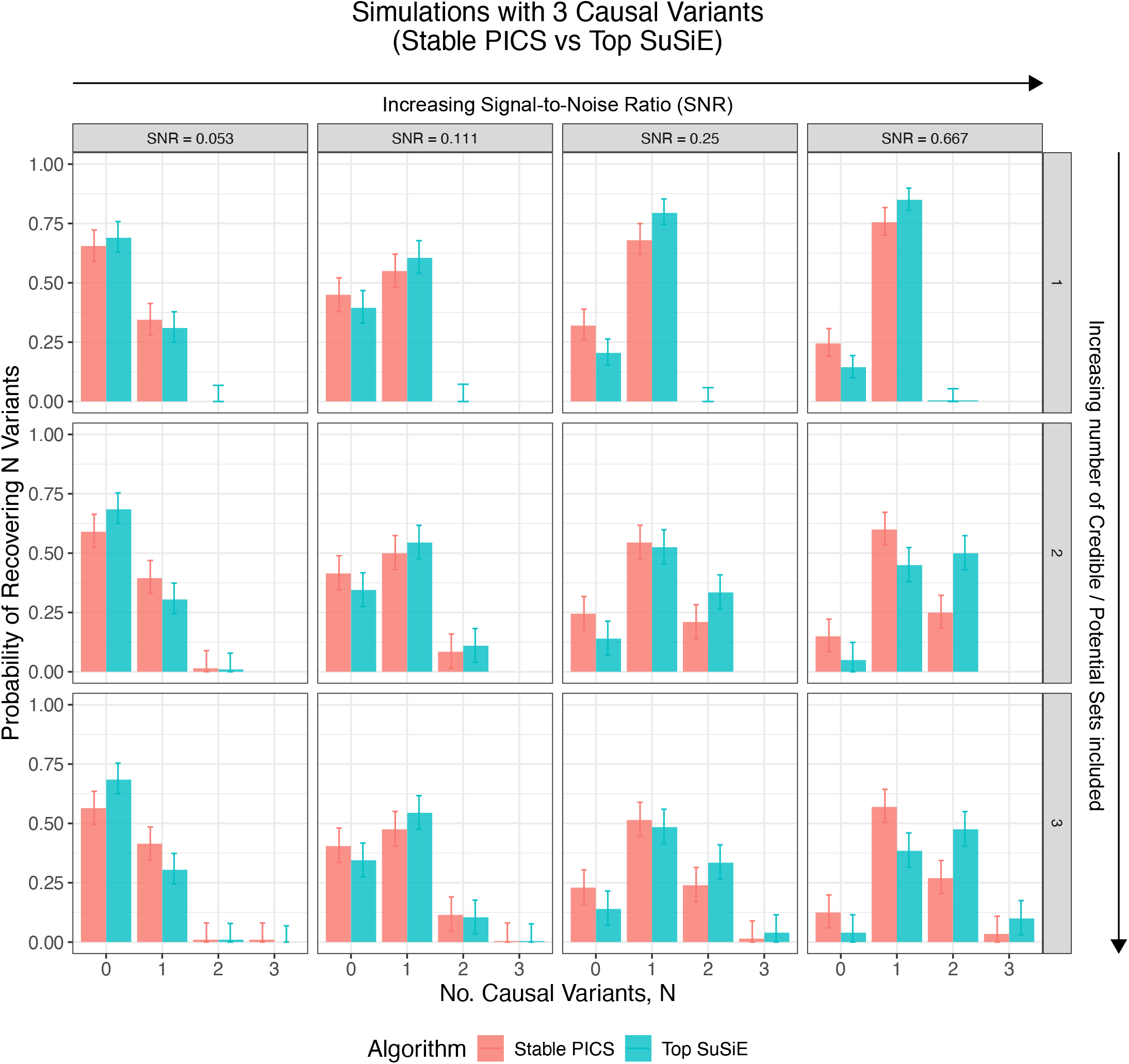
Empirical discrete probability distributions over the number of causal variants (0, 1, 2 or 3) recovered by Stable PICS or Top SuSiE in simulations with three causal variants, stratified by the SNR parameter used in simulations (increasing SNR from left to right). The impact of including more number of credible or potential sets on the distribution is shown (increasing number of included sets from top to bottom).

**Figure 2—figure supplement 13.**
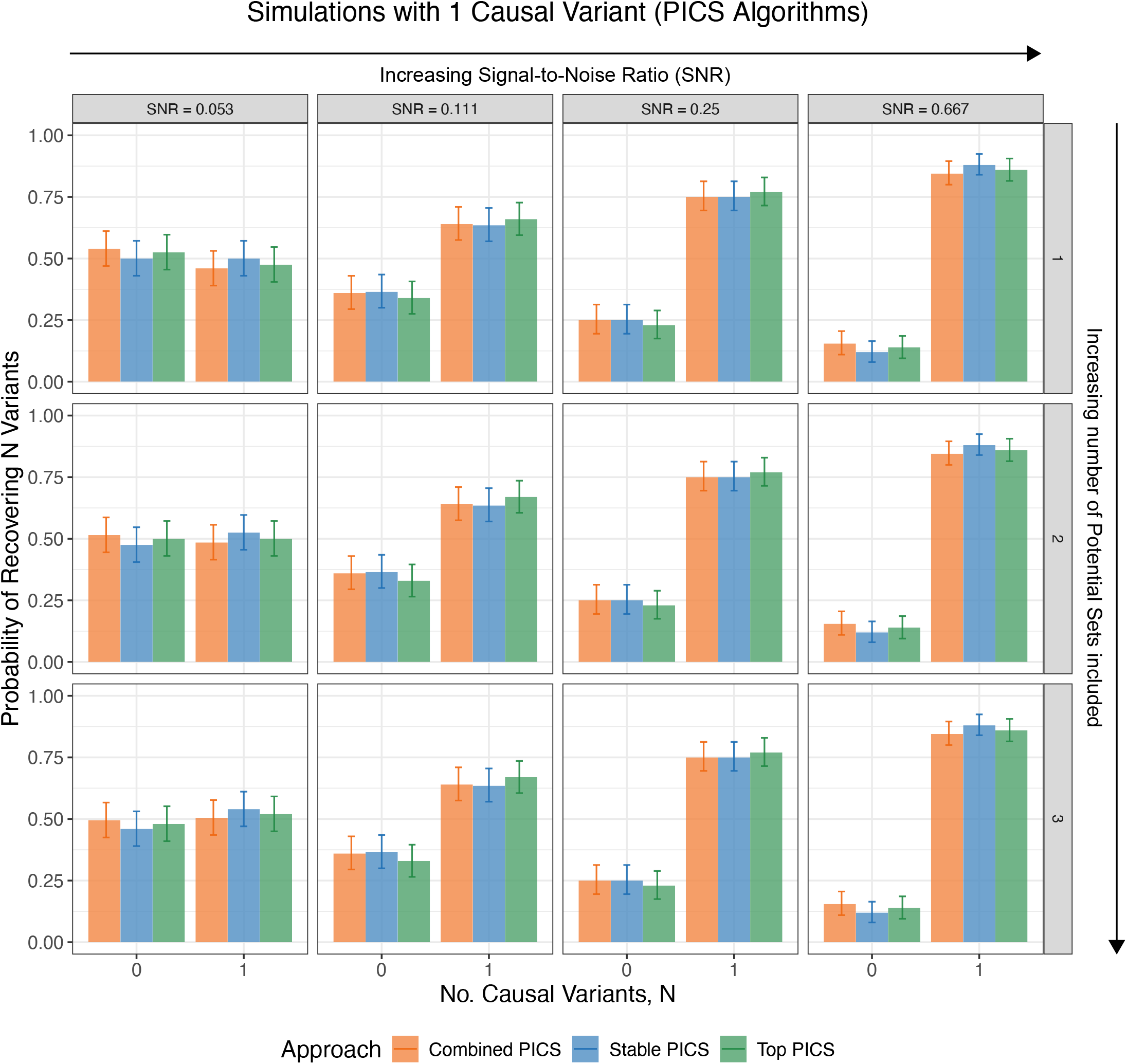
Empirical discrete probability distributions over the number of causal variants (0 or 1) recovered by PICS algorithms in simulations with one causal variant, stratified by the SNR parameter used in simulations (increasing SNR from left to right). The impact of including more number of potential sets on the distribution is shown (increasing number of included sets from top to bottom).

**Figure 2—figure supplement 14.**
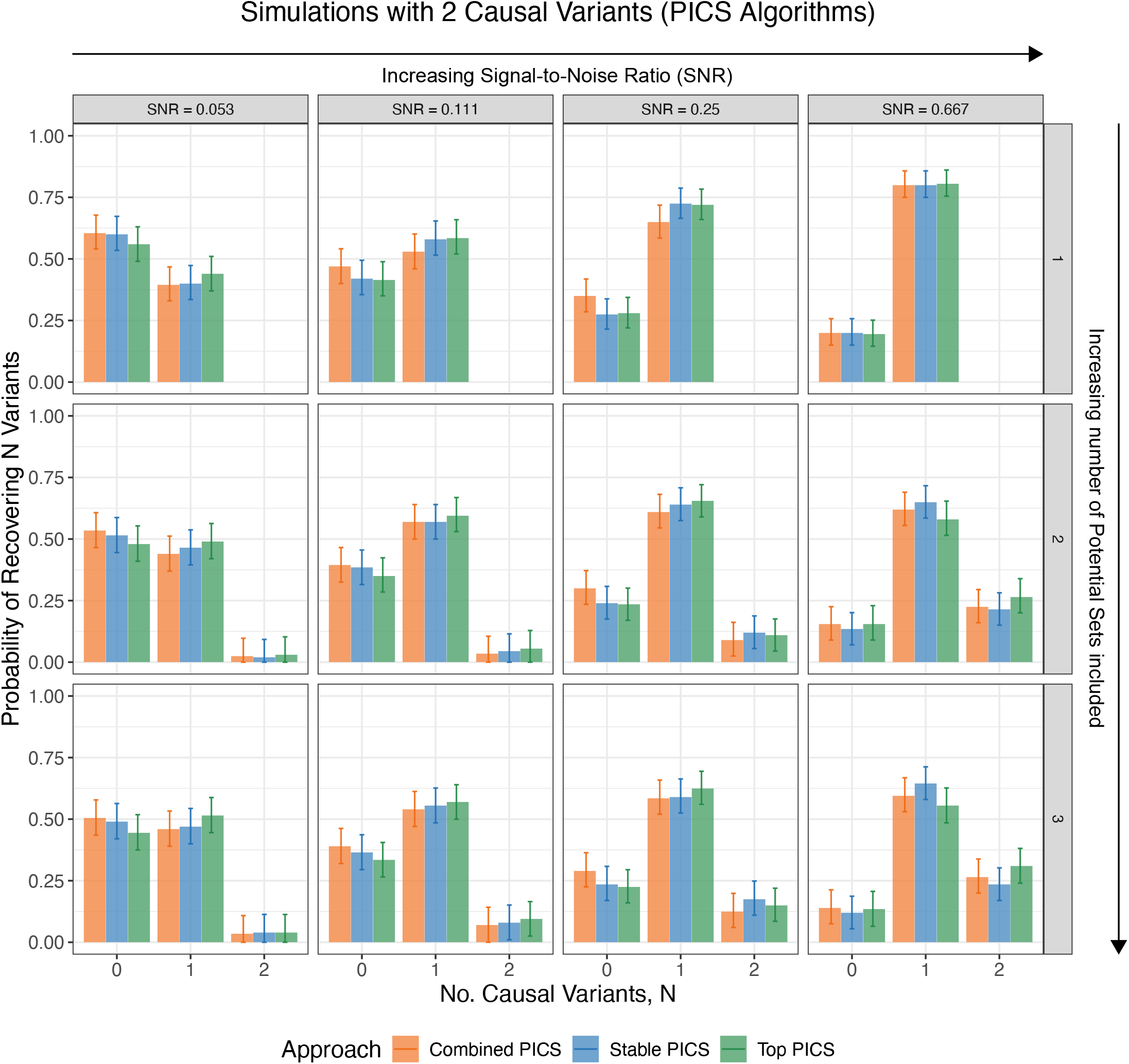
Empirical discrete probability distributions over the number of causal variants (0, 1 or 2) recovered by PICS algorithms in simulations with two causal variants, stratified by the SNR parameter used in simulations (increasing SNR from left to right). The impact of including more number of potential sets on the distribution is shown (increasing number of included sets from top to bottom).

**Figure 2—figure supplement 15.**
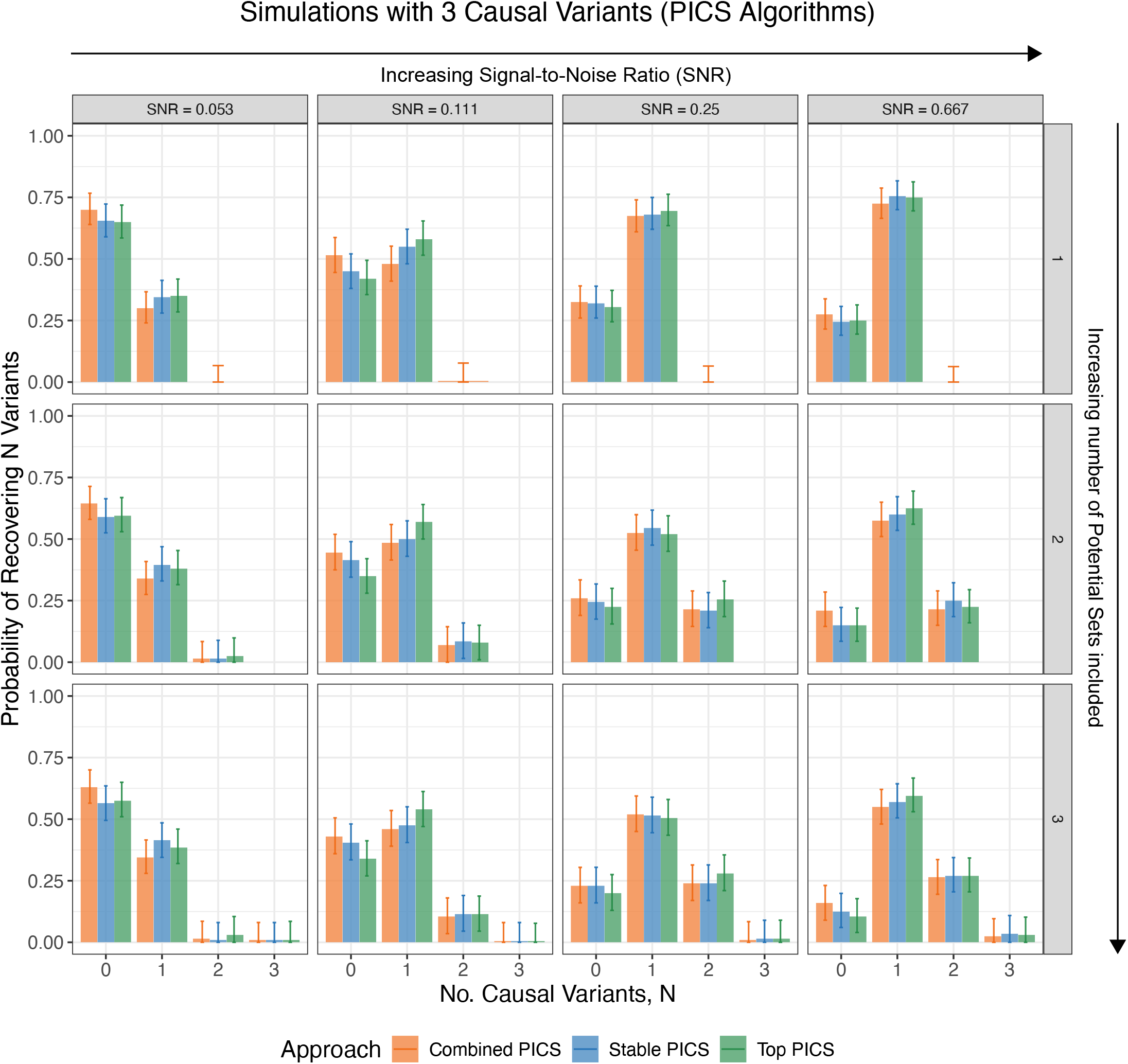
Empirical discrete probability distributions over the number of causal variants (0, 1, 2 or 3) recovered by PICS algorithms in simulations with three causal variants, stratified by the SNR parameter used in simulations (increasing SNR from left to right). The impact of including more number of potential sets on the distribution is shown (increasing number of included sets from top to bottom).

**Figure 2—figure supplement 16.**
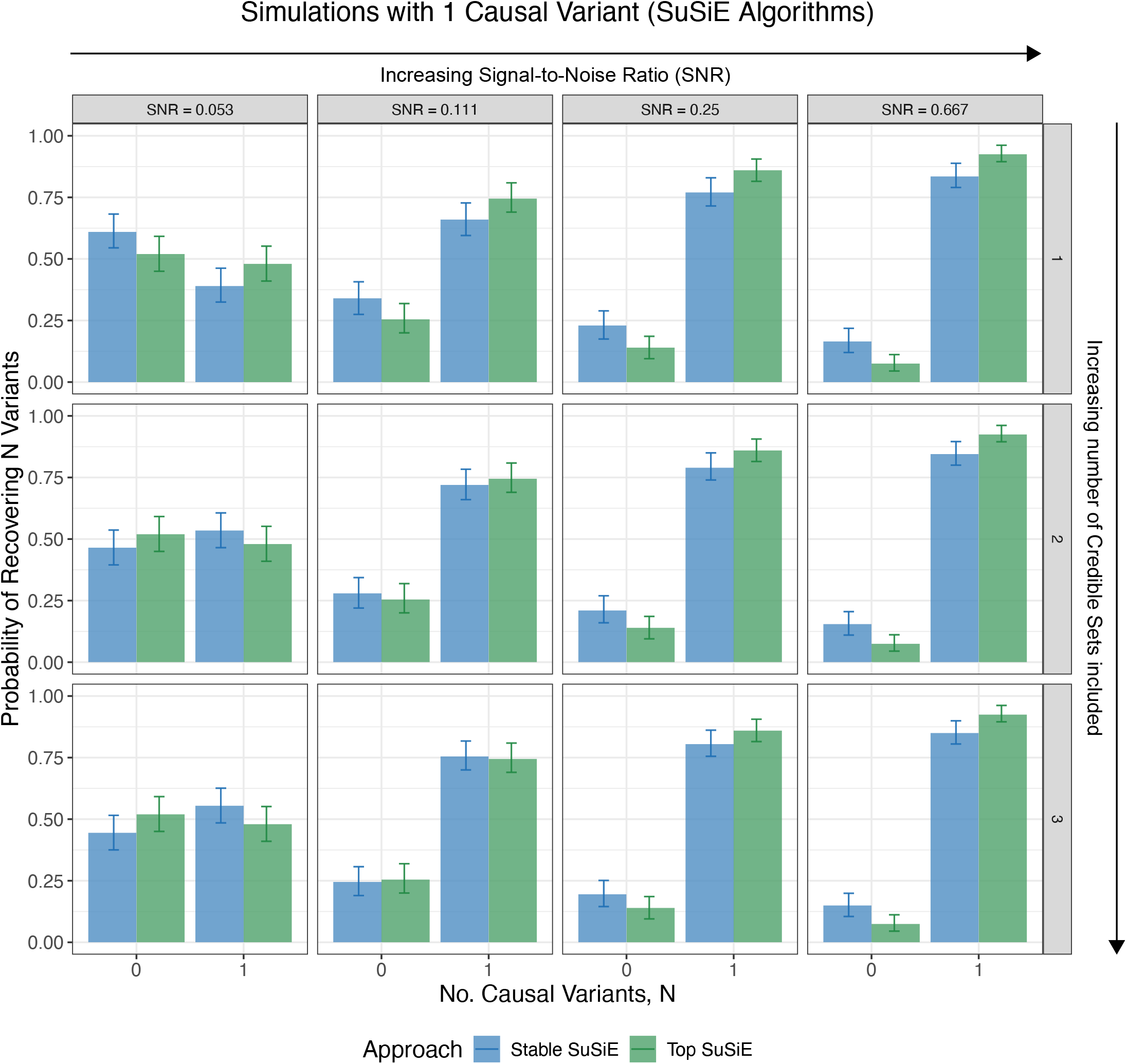
Empirical discrete probability distributions over the number of causal variants (0 or 1) recovered by SuSiE algorithms in simulations with one causal variant, stratified by the SNR parameter used in simulations (increasing SNR from left to right). The impact of including more number of credible sets on the distribution is shown (increasing number of included sets from top to bottom).

**Figure 2—figure supplement 17.**
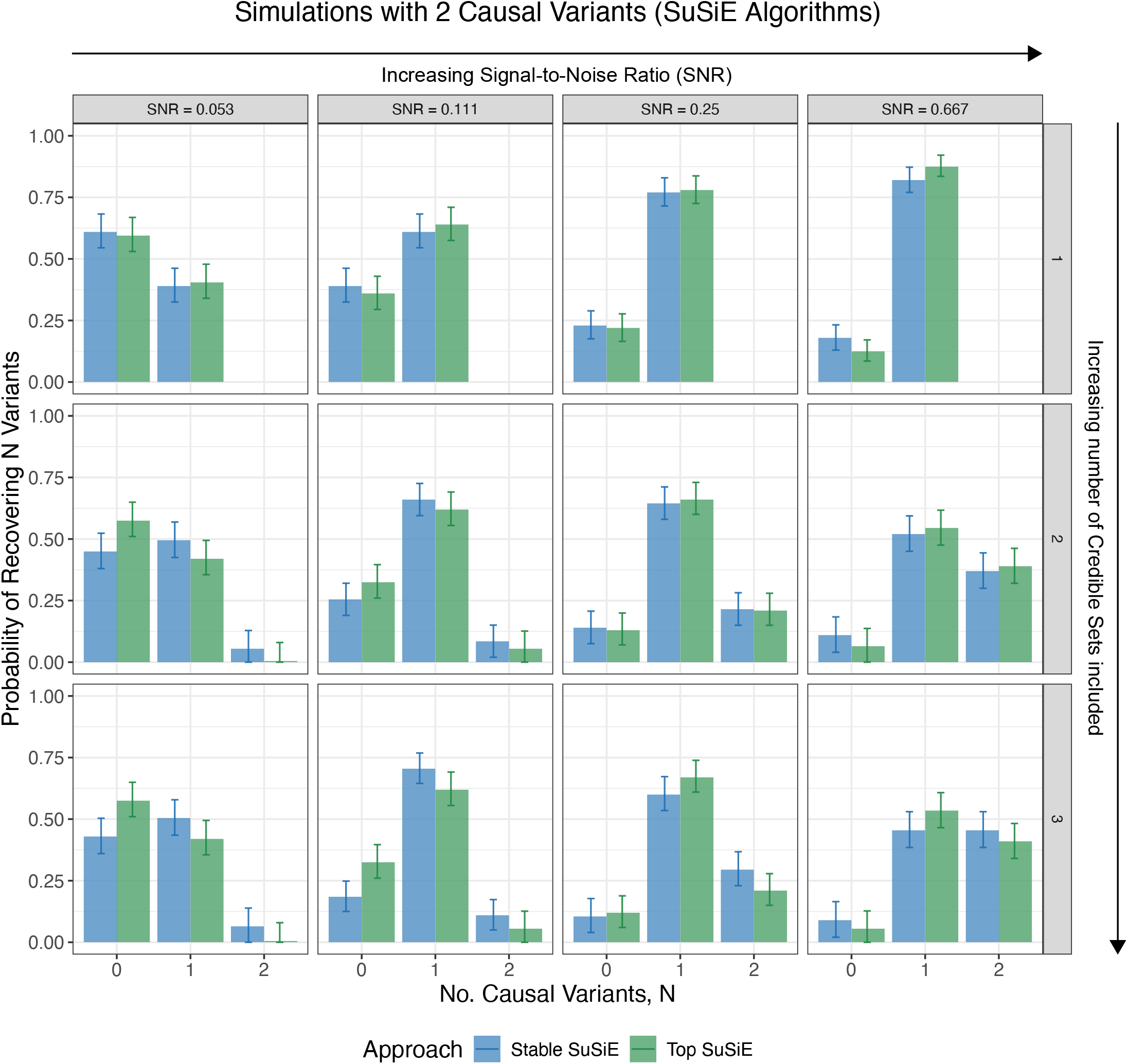
Empirical discrete probability distributions over the number of causal variants (0, 1 or 2) recovered by SuSiE algorithms in simulations with two causal variants, stratified by the SNR parameter used in simulations (increasing SNR from left to right). The impact of including more number of credible sets on the distribution is shown (increasing number of included sets from top to bottom).

**Figure 2—figure supplement 18.**
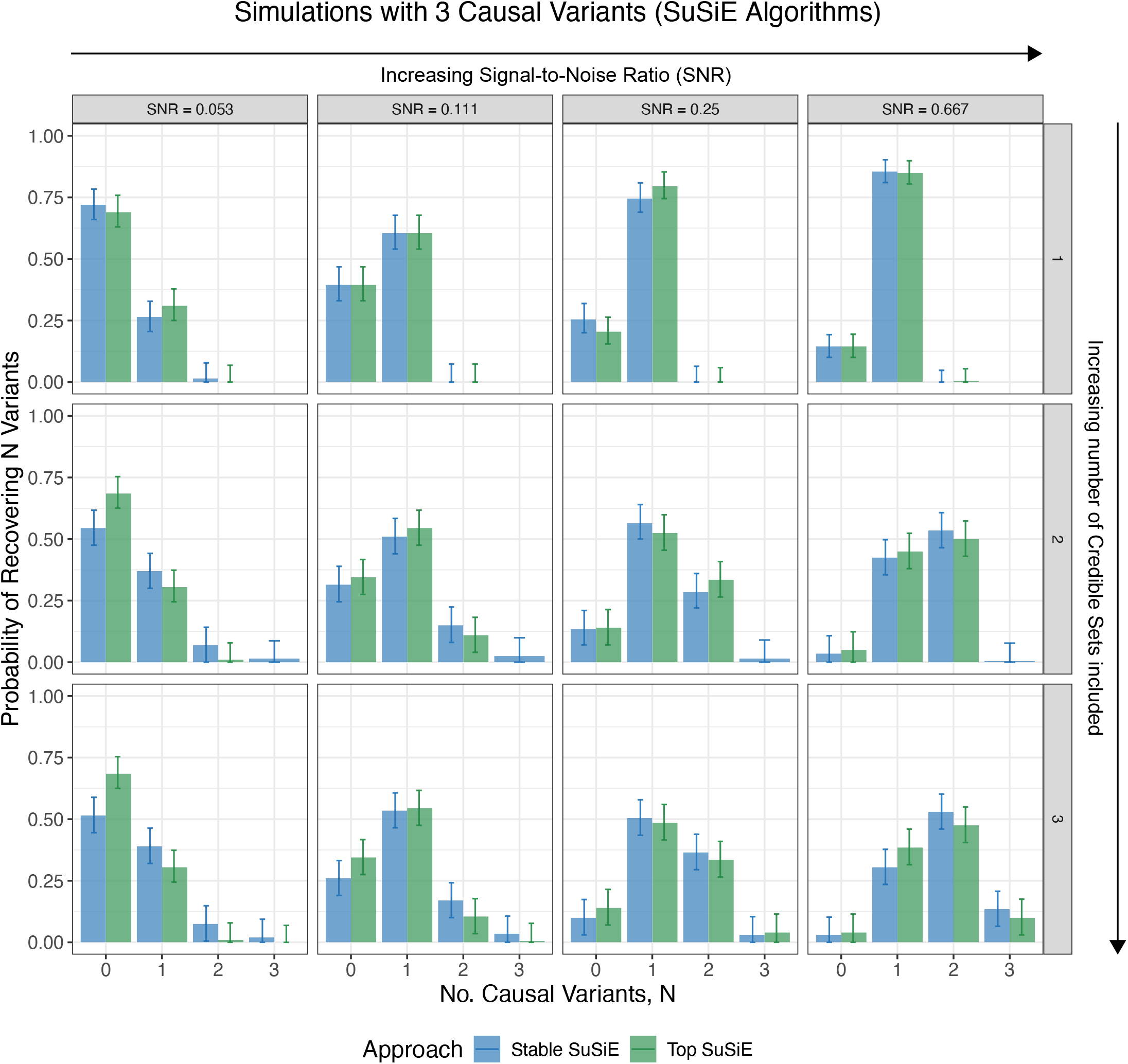
Empirical discrete probability distributions over the number of causal variants (0, 1, 2 or 3) recovered by SuSiE algorithms in simulations with three causal variants, stratified by the SNR parameter used in simulations (increasing SNR from left to right). The impact of including more number of credible sets on the distribution is shown (increasing number of included sets from top to bottom).

**Figure 2—figure supplement 19.**
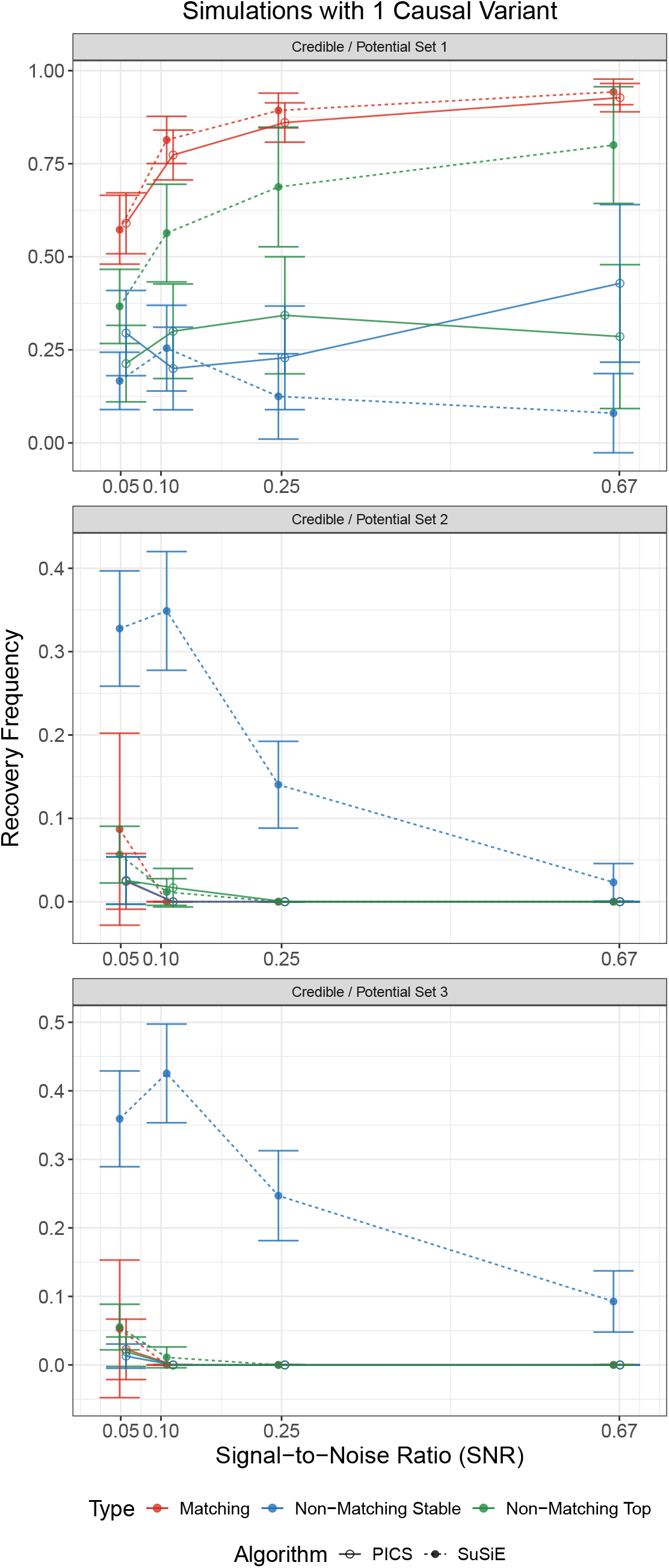
Frequency with which the causal variant is recovered by a matching variant, non-matching top variant or non-matching stable variant in simulations involving one causal variant. Recovery frequencies are stratified by simulations differing in the signal-to-noise ratio (SNR) parameter *ϕ*.

**Figure 2—figure supplement 20.**
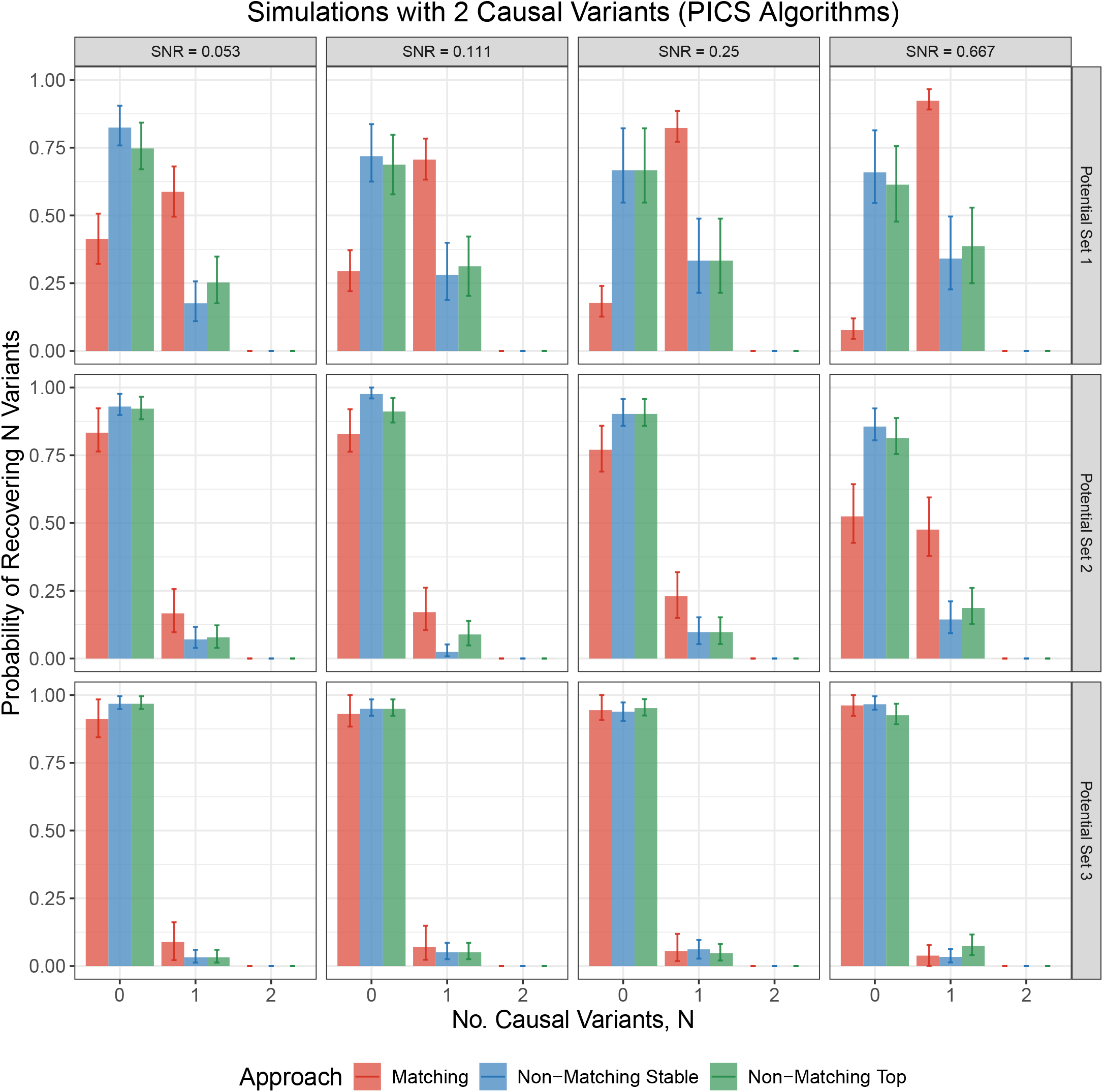
Empirical discrete probability distributions over the number of causal variants (0, 1 or 2) recovered by matching Top and Stable PICS variants, non-matching Top PICS variants and non-matching Stable PICS variants, in simulations involving two causal variants. Empirical distributions are stratified by simulations differing in the signal-to-noise ratio (SNR) parameter *ϕ* (increasing SNR from left to right).

**Figure 2—figure supplement 21.**
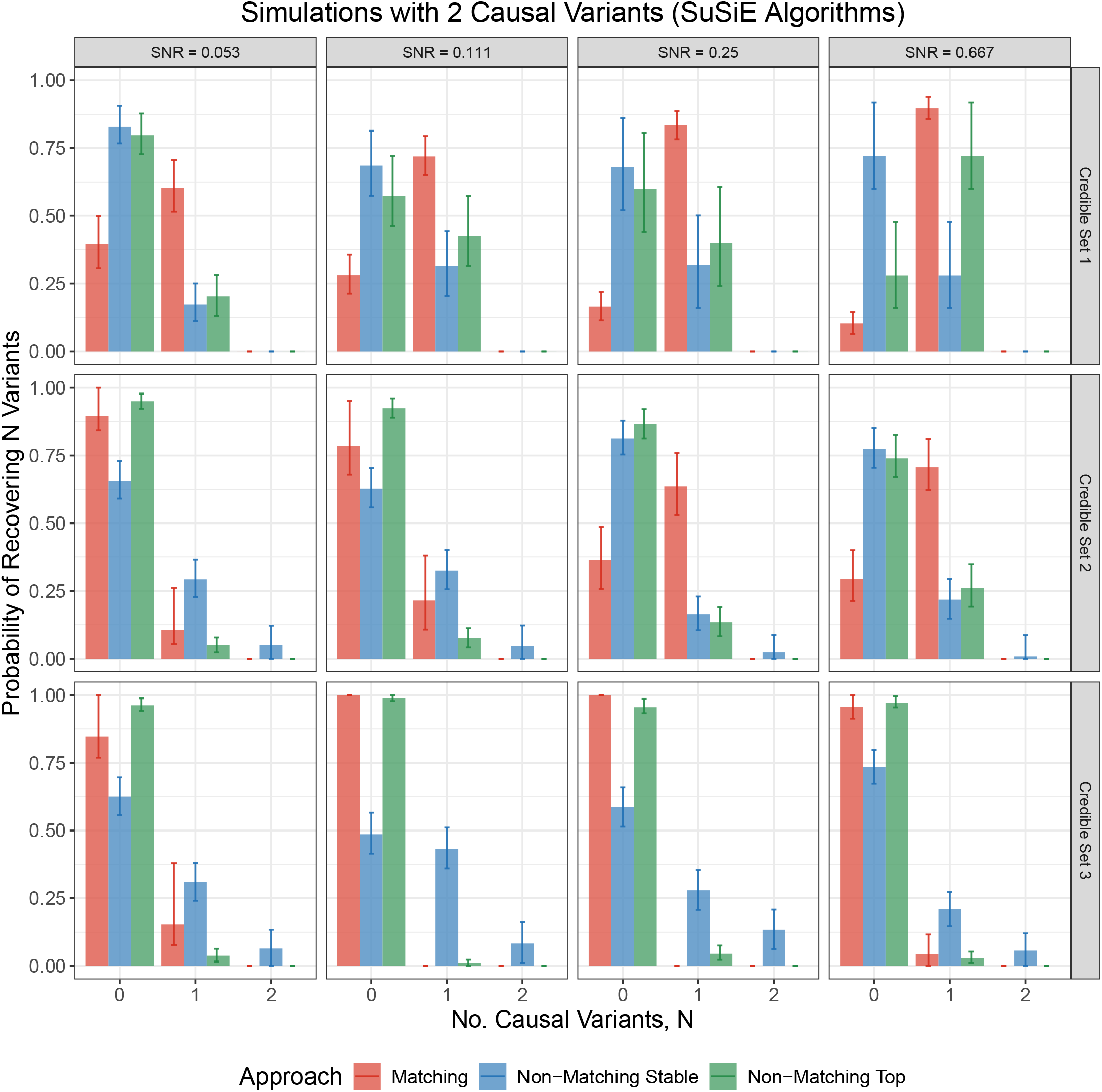
Empirical discrete probability distributions over the number of causal variants (0, 1 or 2) recovered by matching Top and Stable SuSiE variants, non-matching Top SuSiE variants and non-matching Stable SuSiE variants, in simulations involving two causal variants. Empirical distributions are stratified by simulations differing in the signal-to-noise ratio (SNR) parameter *ϕ* (increasing SNR from left to right).

**Figure 2—figure supplement 22.**
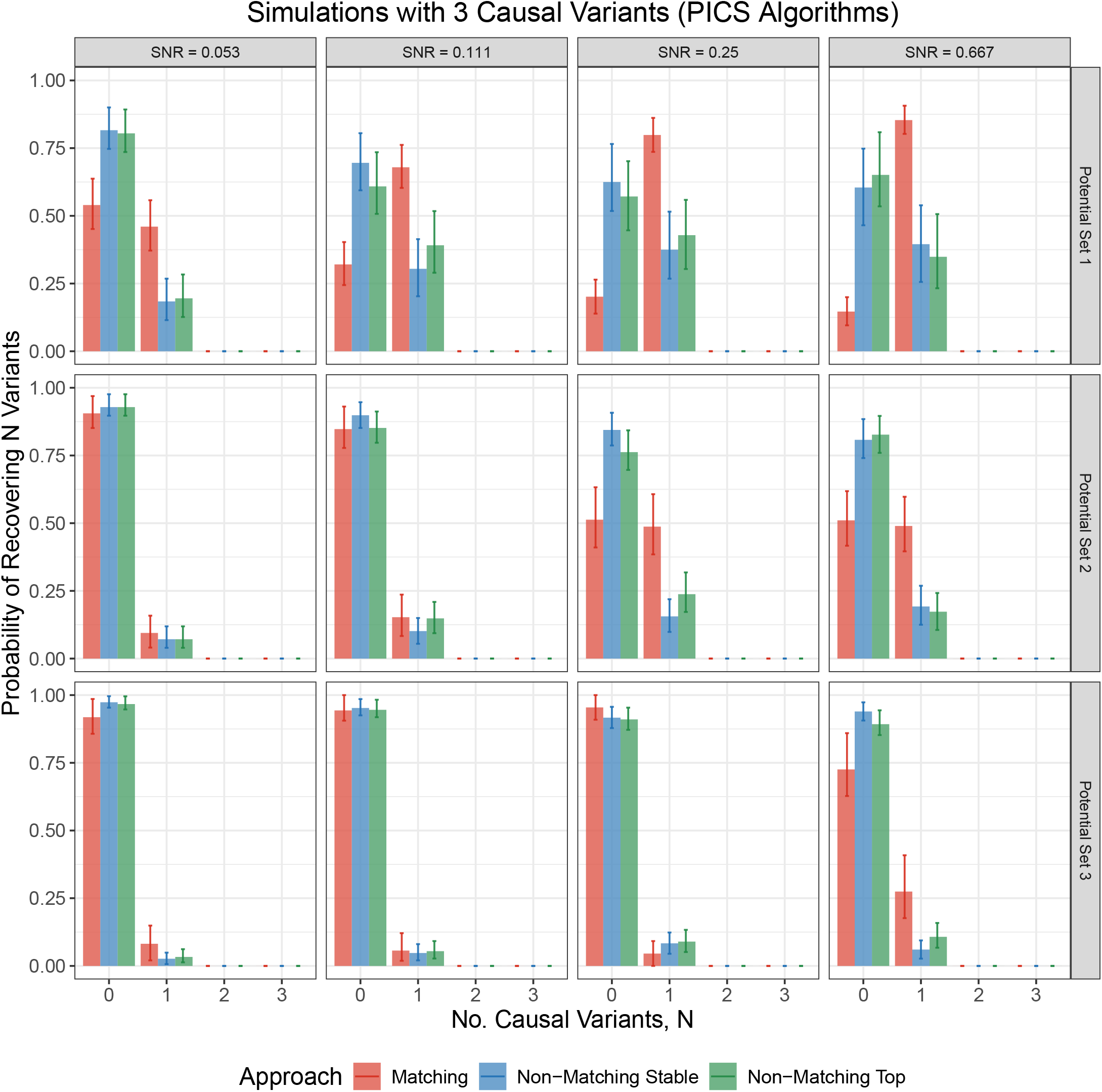
Empirical discrete probability distributions over the number of causal variants (0, 1, 2 or 3) recovered by matching Top and Stable PICS variants, non-matching Top PICS variants and non-matching Stable PICS variants, in simulations involving three causal variants. Empirical distributions are stratified by simulations differing in the signal-to-noise ratio (SNR) parameter *ϕ* (increasing SNR from left to right).

**Figure 2—figure supplement 23.**
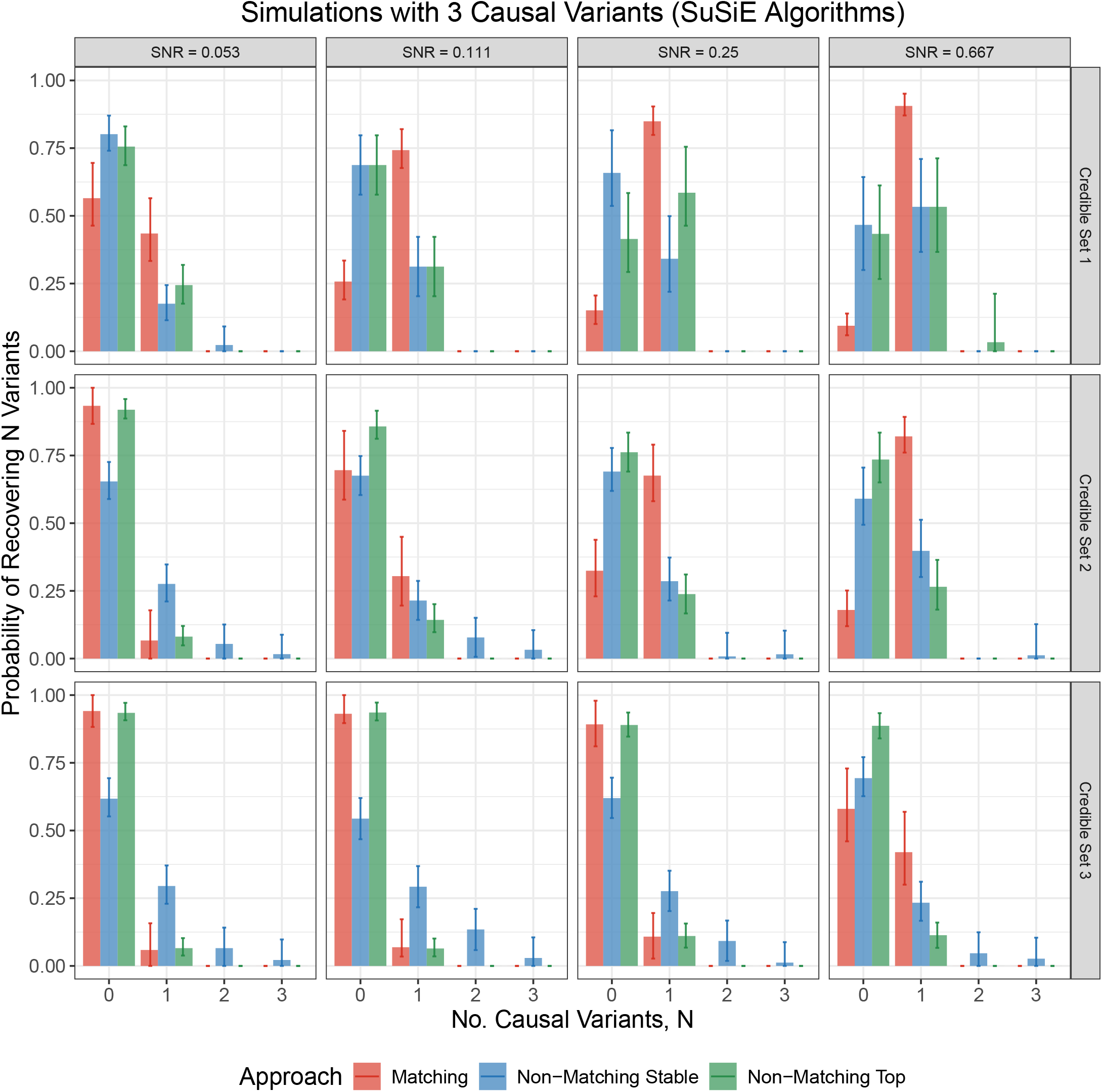
Empirical discrete probability distributions over the number of causal variants (0, 1, 2 or 3) recovered by matching Top and Stable SuSiE variants, non-matching Top SuSiE variants and non-matching Stable SuSiE variants, in simulations involving three causal variants. Empirical distributions are stratified by simulations differing in the signal-to-noise ratio (SNR) parameter *ϕ* (increasing SNR from left to right).

**Figure 2—figure supplement 24.**
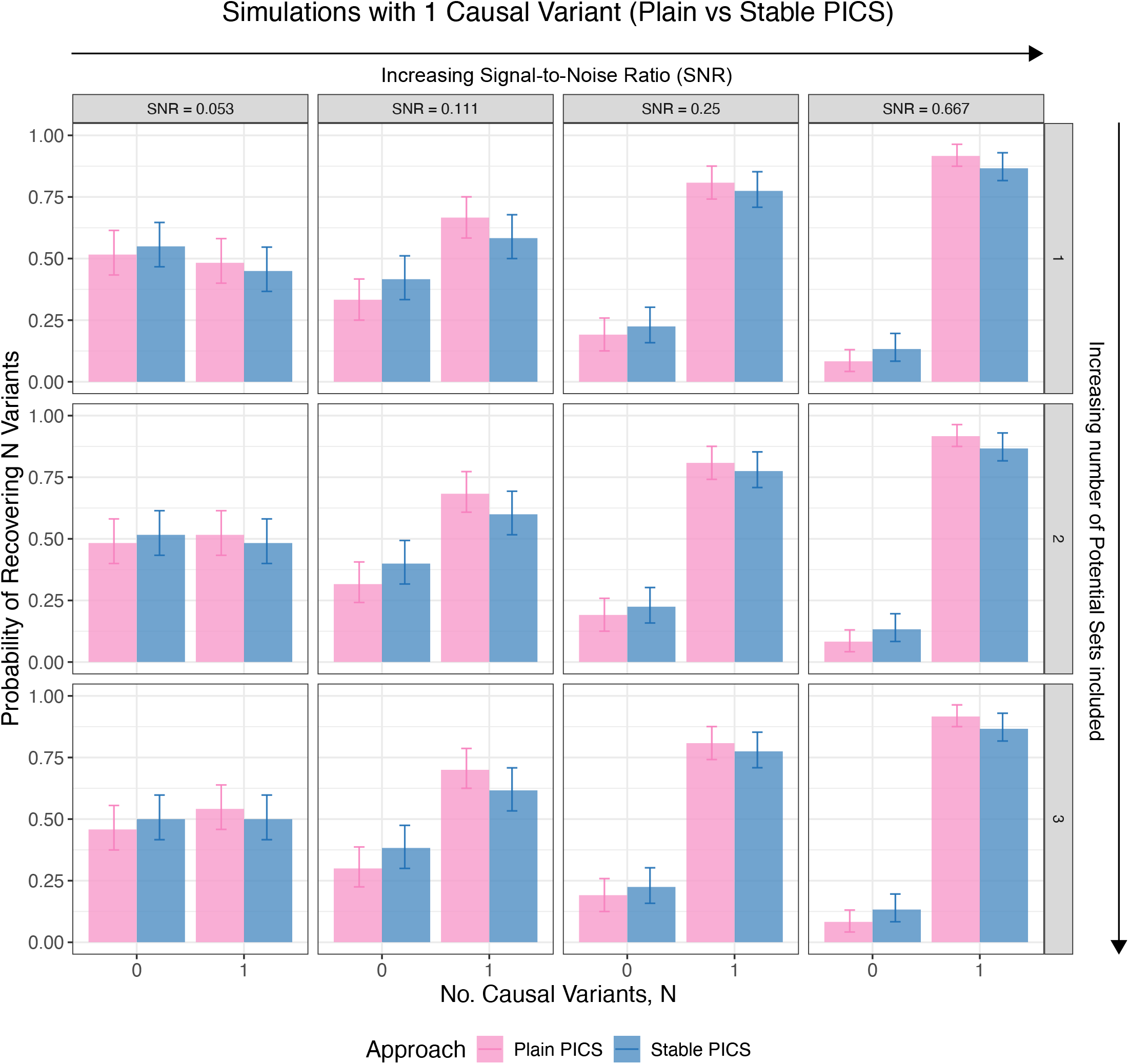
Empirical discrete probability distributions over the number of causal variants (0 or 1) recovered by Plain or Stable PICS in simulations with one causal variant and environmental heterogeneity, stratified by the SNR parameter used in simulations (increasing SNR from left to right). The impact of including more number of potential sets on the distribution is shown (increasing number of included sets from top to bottom).

**Figure 2—figure supplement 25.**
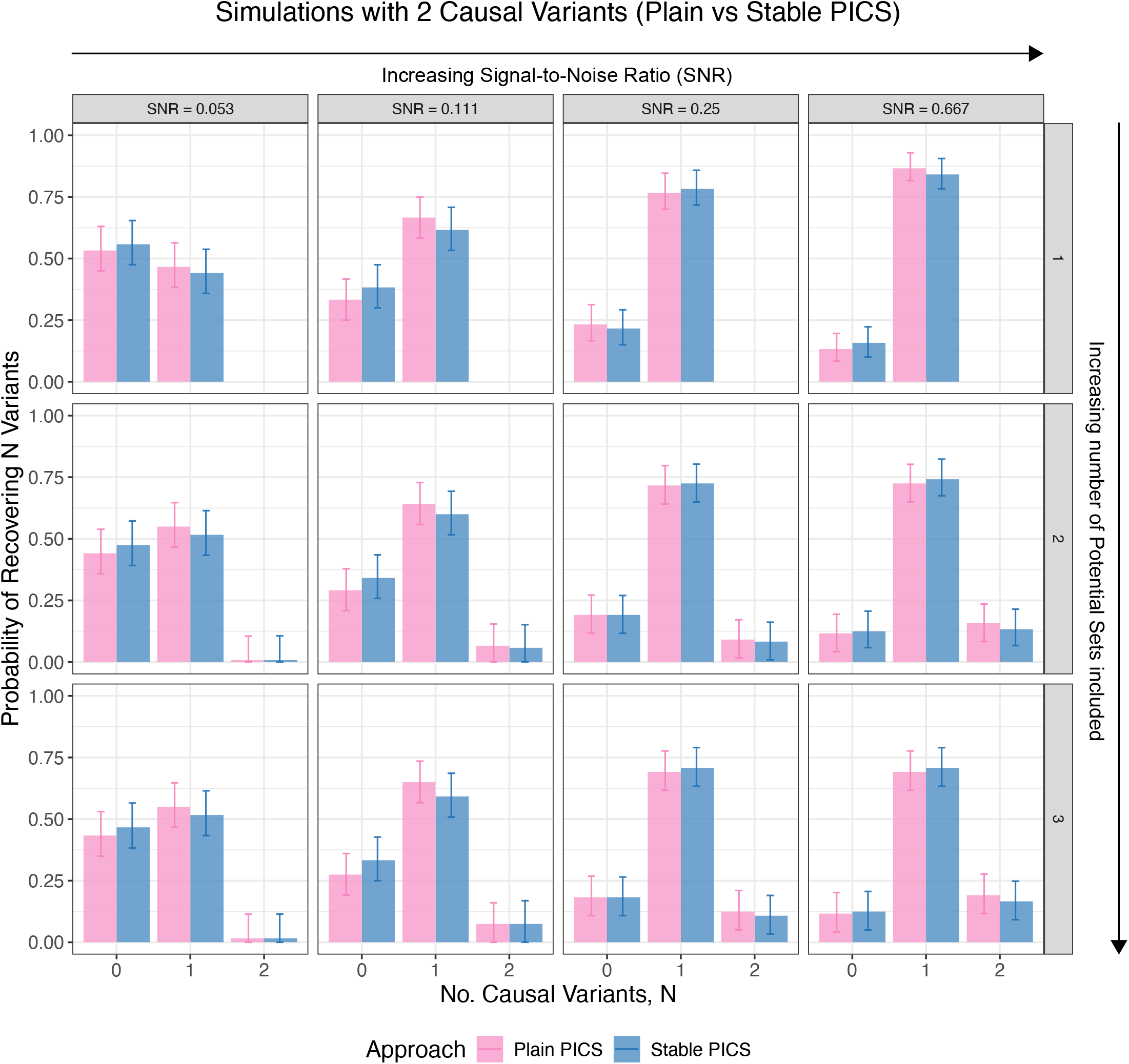
Empirical discrete probability distributions over the number of causal variants (0, 1 or 2) recovered by Plain or Stable PICS in simulations with two causal variants and environmental heterogeneity, stratified by the SNR parameter used in simulations (increasing SNR from left to right). The impact of including more number of potential sets on the distribution is shown (increasing number of included sets from top to bottom).

**Figure 2—figure supplement 26.**
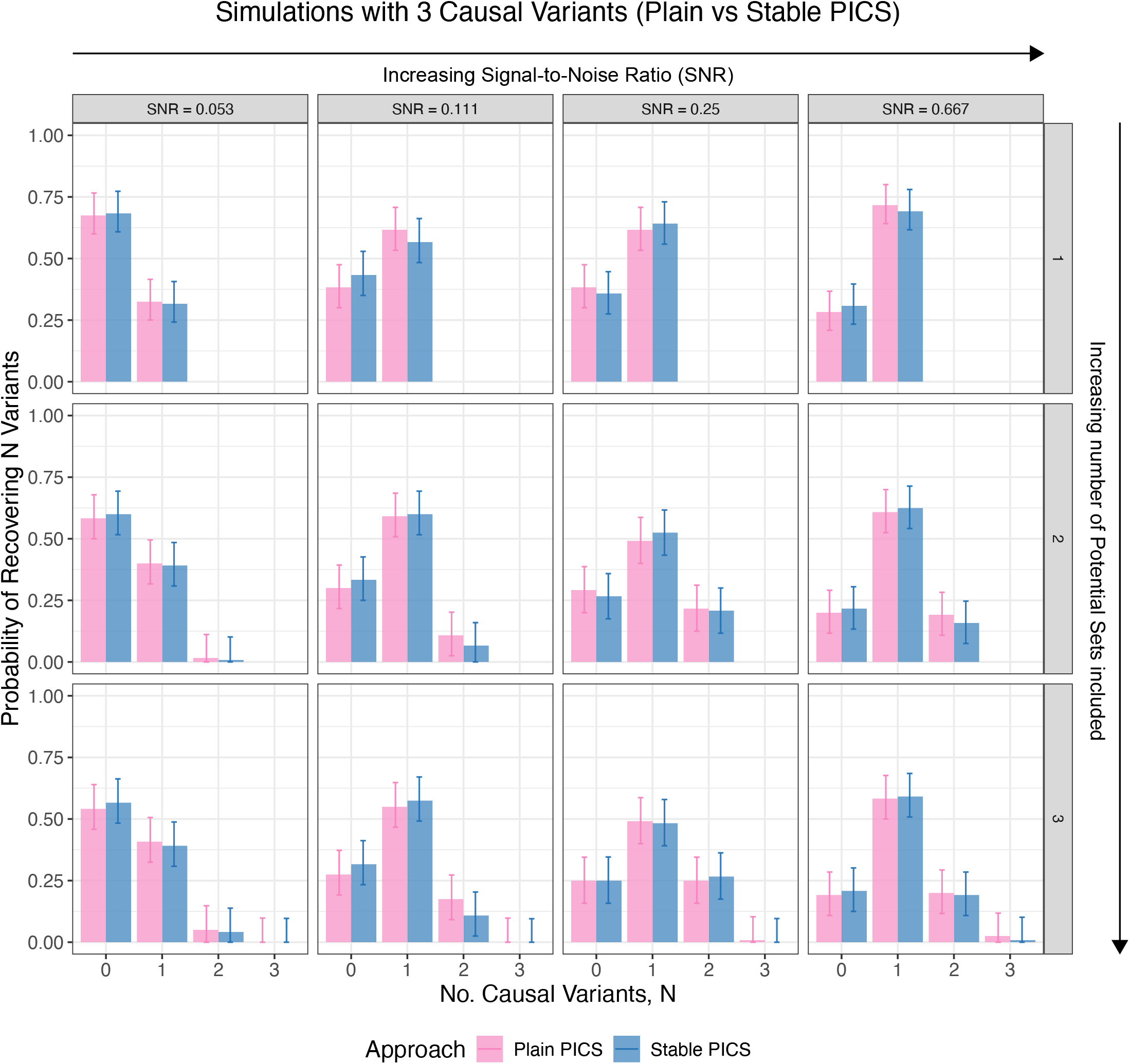
Empirical discrete probability distributions over the number of causal variants (0, 1, 2 or 3) recovered by Plain or Stable PICS in simulations with three causal variants and environmental heterogeneity, stratified by the SNR parameter used in simulations (increasing SNR from left to right). The impact of including more number of potential sets on the distribution is shown (increasing number of included sets from top to bottom).

**Figure 2—figure supplement 27.**
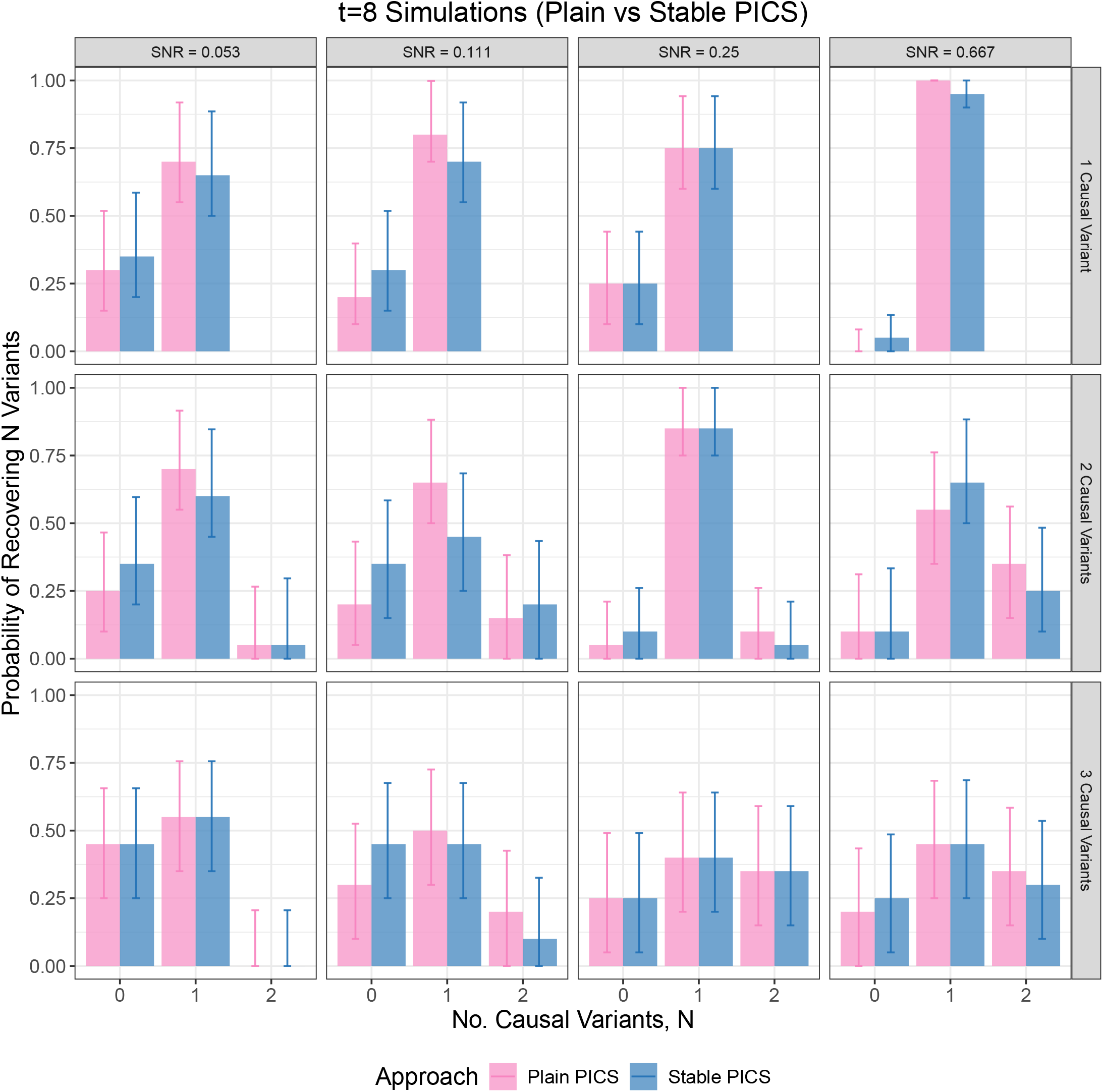
Empirical discrete probability distributions over the number of causal variants (0, 1, 2 or 3) recovered by Plain or Stable PICS in “*t* = 8” simulations involving environmental heterogeneity (variance shift scenario), stratified by the SNR parameter used in simulations (increasing SNR from left to right). Each row reports the distribution for simulations with a specific number of causal variants (1, 2 or 3), and we use all three potential sets to compute the number of causal variants recovered in each case.

**Figure 2—figure supplement 28.**
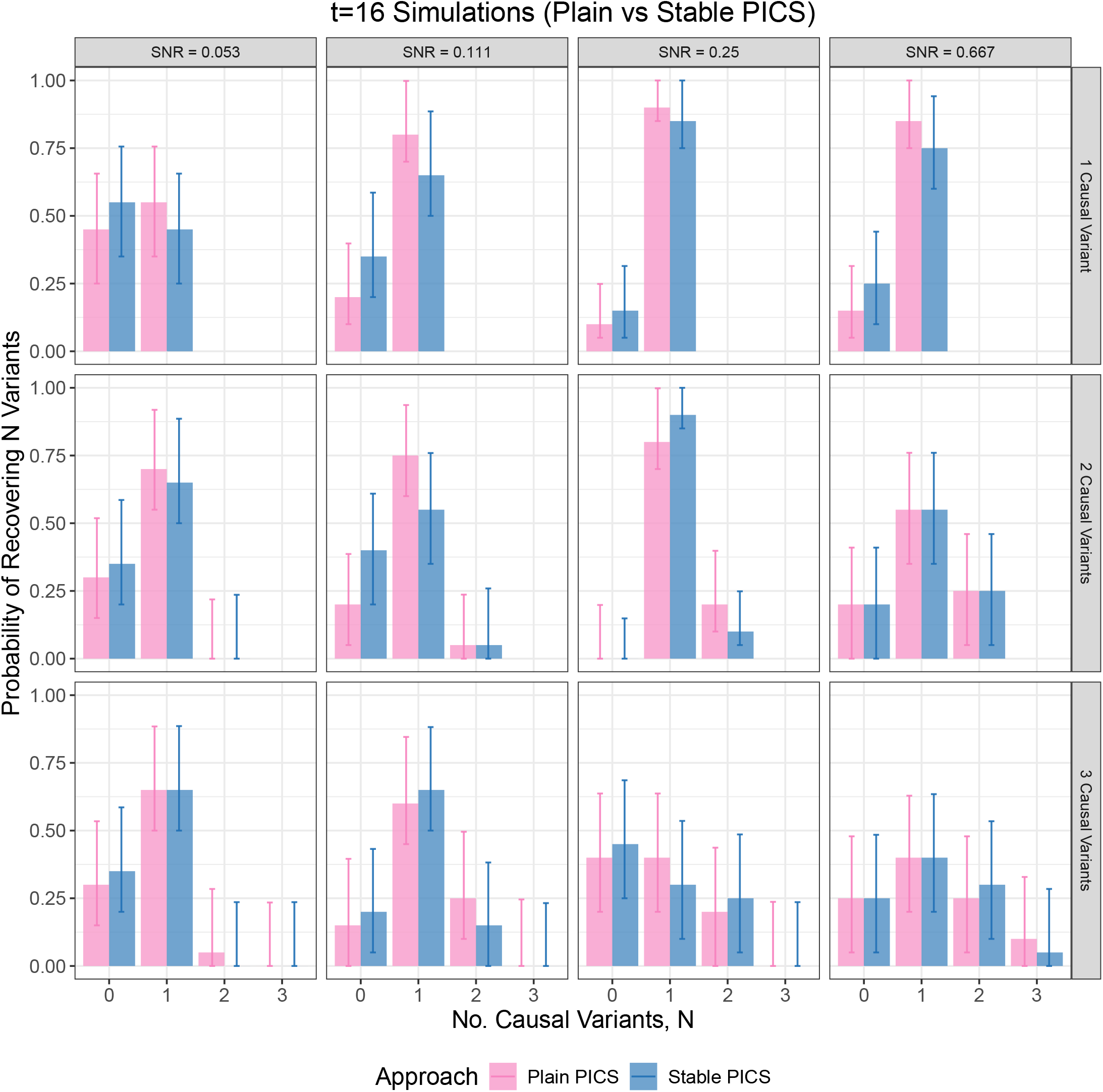
Empirical discrete probability distributions over the number of causal variants (0, 1, 2 or 3) recovered by Plain or Stable PICS in “*t* = 16” simulations involving environmental heterogeneity (variance shift scenario), stratified by the SNR parameter used in simulations (increasing SNR from left to right). Each row reports the distribution for simulations with a specific number of causal variants (1, 2 or 3), and we use all three potential sets to compute the number of causal variants recovered in each case.

**Figure 2—figure supplement 29.**
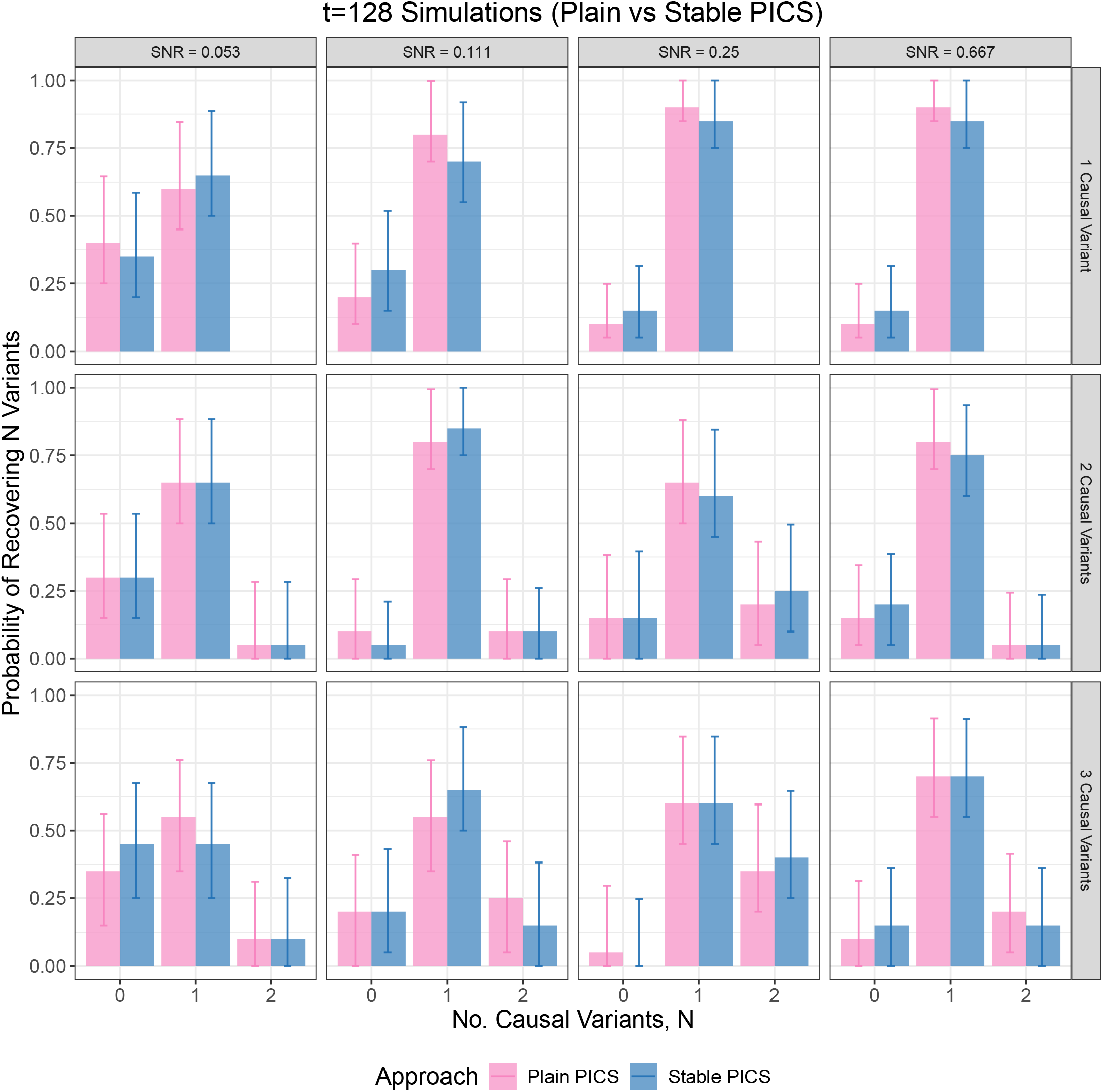
Empirical discrete probability distributions over the number of causal variants (0, 1, 2 or 3) recovered by Plain or Stable PICS in “*t* = 128” simulations involving environmental heterogeneity (variance shift scenario), stratified by the SNR parameter used in simulations (increasing SNR from left to right). Each row reports the distribution for simulations with a specific number of causal variants (1, 2 or 3), and we use all three potential sets to compute the number of causal variants recovered in each case.

**Figure 2—figure supplement 30.**
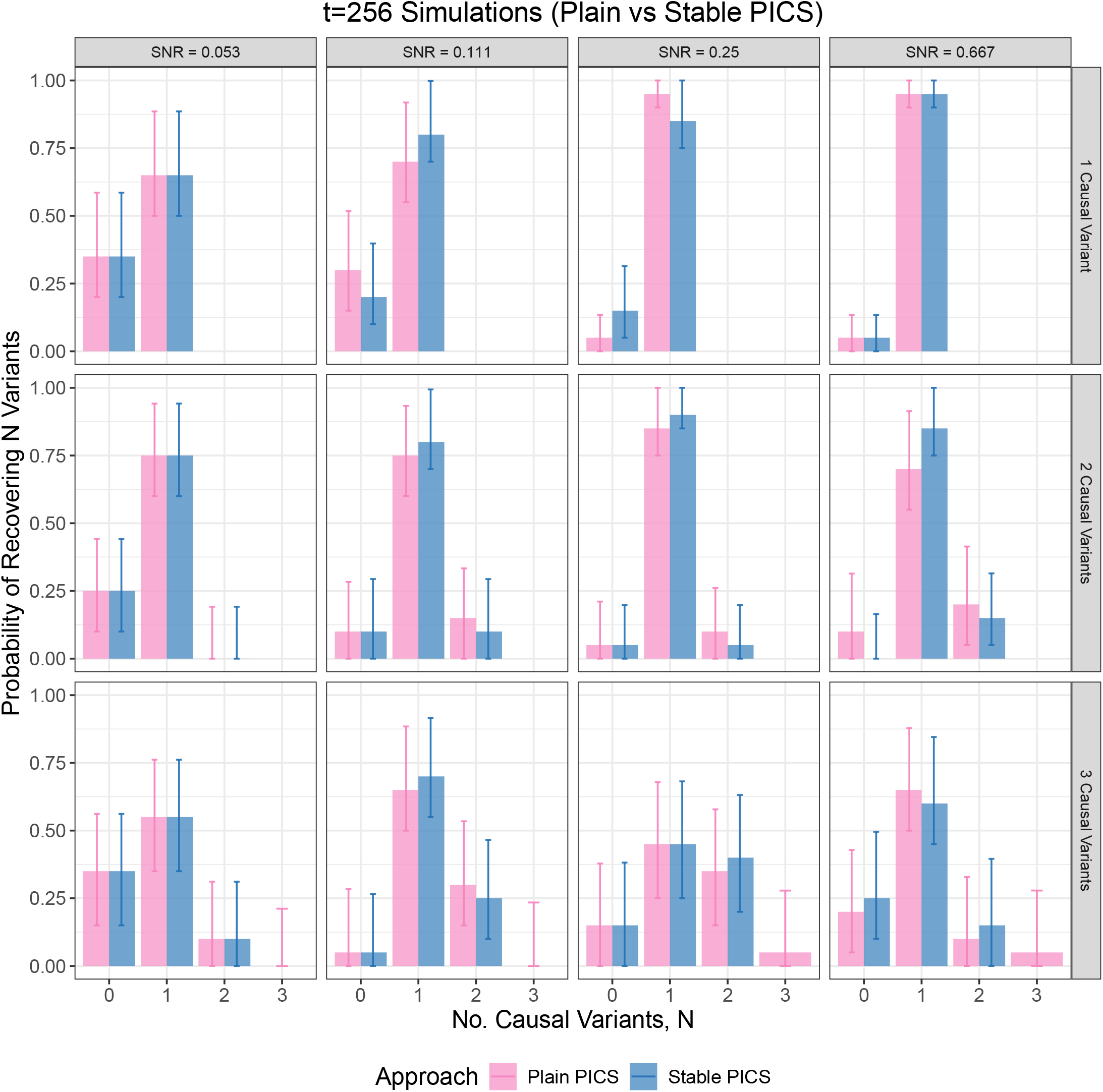
Empirical discrete probability distributions over the number of causal variants (0, 1, 2 or 3) recovered by Plain or Stable PICS in “*t* = 256” simulations involving environmental heterogeneity (variance shift scenario), stratified by the SNR parameter used in simulations (increasing SNR from left to right). Each row reports the distribution for simulations with a specific number of causal variants (1, 2 or 3), and we use all three potential sets to compute the number of causal variants recovered in each case.

**Figure 2—figure supplement 31.**
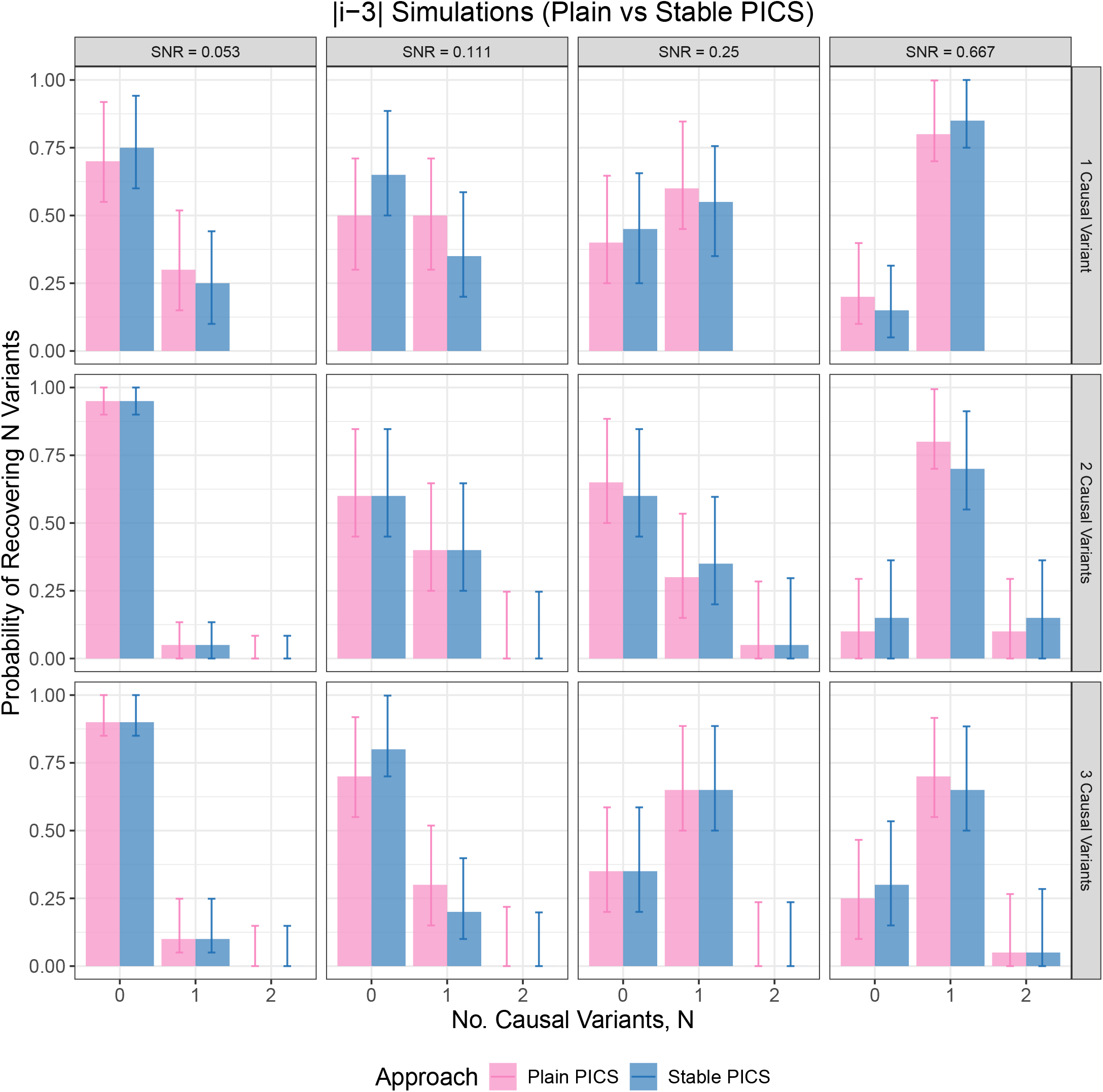
Empirical discrete probability distributions over the number of causal variants (0, 1, 2 or 3) recovered by Plain or Stable PICS in “|*i*−3|” simulations involving environmental heterogeneity (mean shift scenario), stratified by the SNR parameter used in simulations (increasing SNR from left to right). Each row reports the distribution for simulations with a specific number of causal variants (1, 2 or 3), and we use all three potential sets to compute the number of causal variants recovered in each case.

**Figure 2—figure supplement 32.**
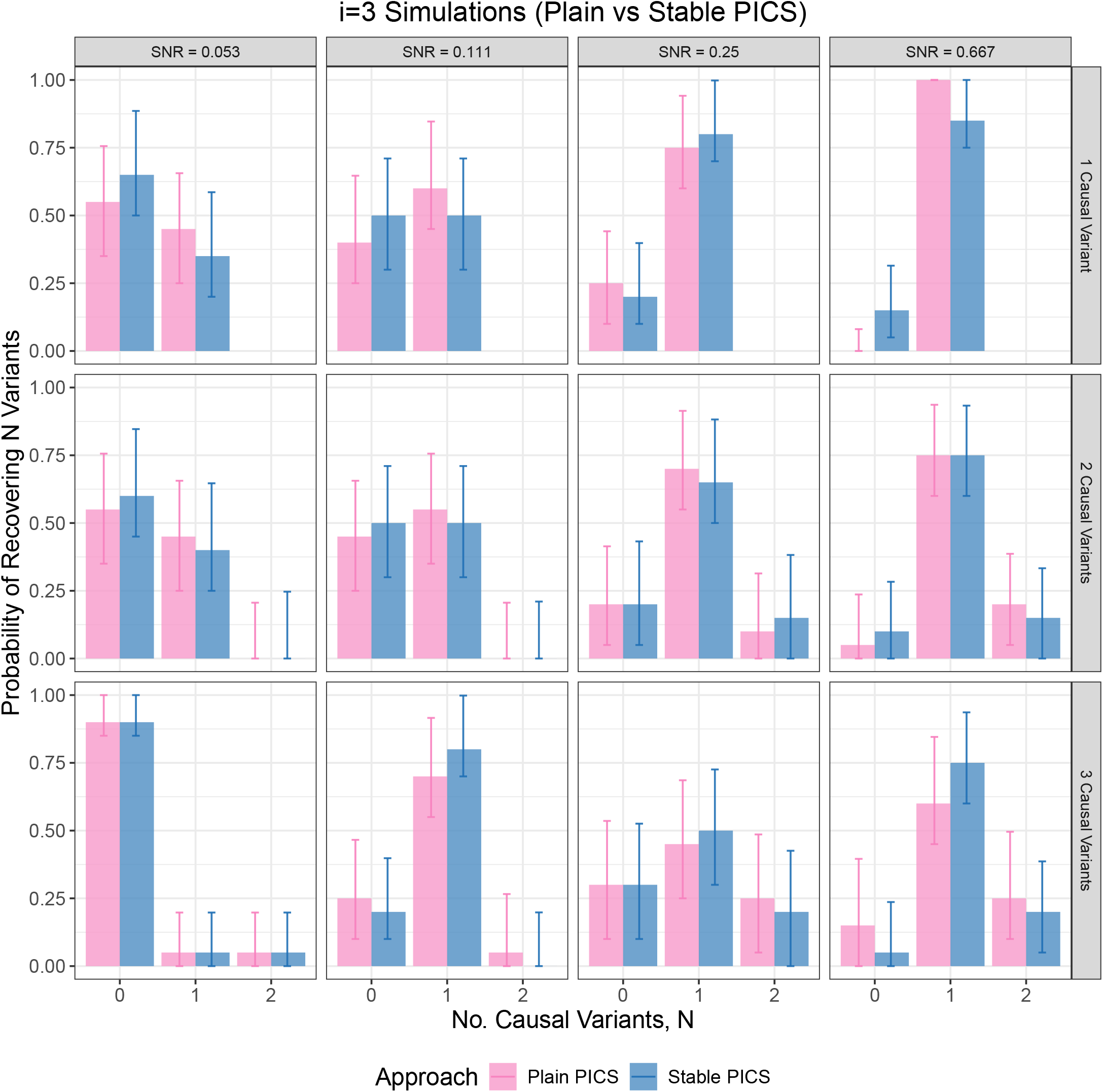
Empirical discrete probability distributions over the number of causal variants (0, 1, 2 or 3) recovered by Plain or Stable PICS in “*i* = 3” simulations involving environmental heterogeneity (mean shift scenario), stratified by the SNR parameter used in simulations (increasing SNR from left to right). Each row reports the distribution for simulations with a specific number of causal variants (1, 2 or 3), and we use all three potential sets to compute the number of causal variants recovered in each case.

**Figure 3—figure supplement 1.**
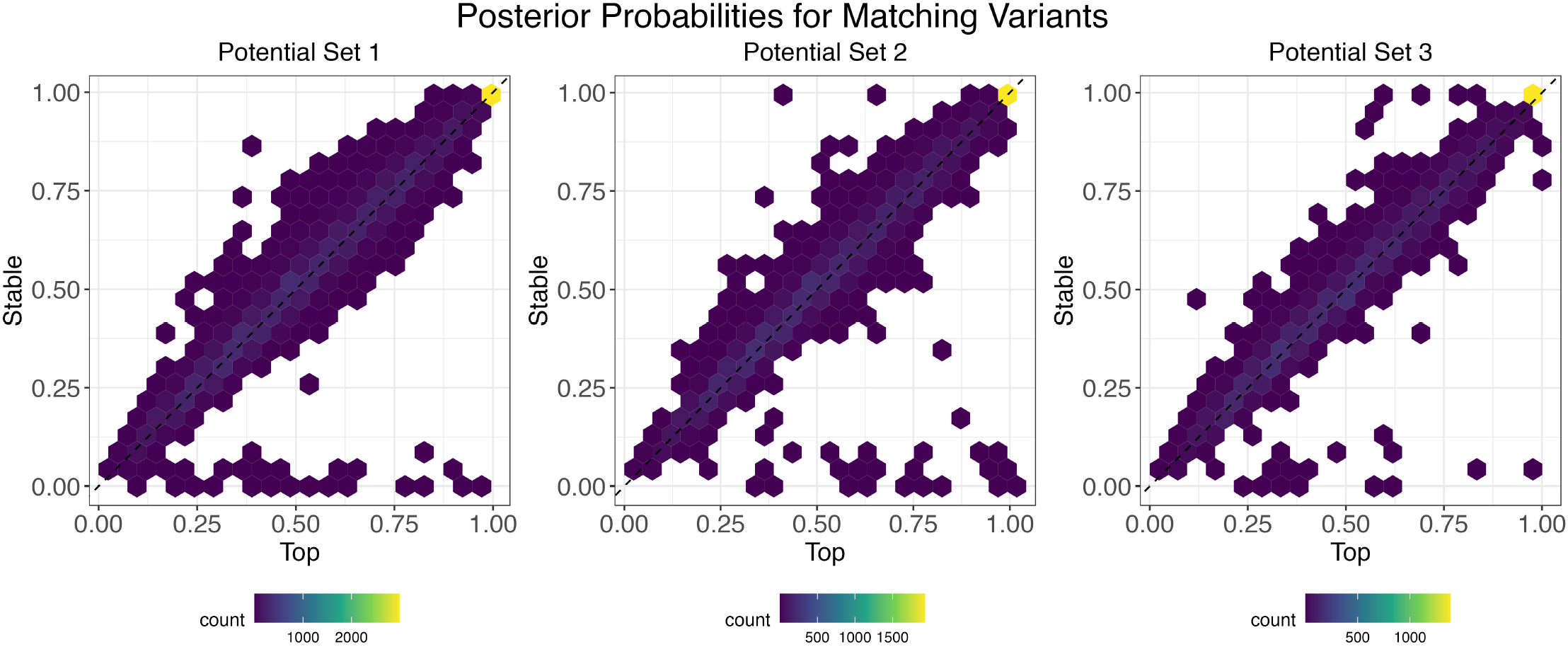
Pair density plot of posterior probabilities of the top variant and the stable variant, in case they match.

**Figure 3—figure supplement 2.**
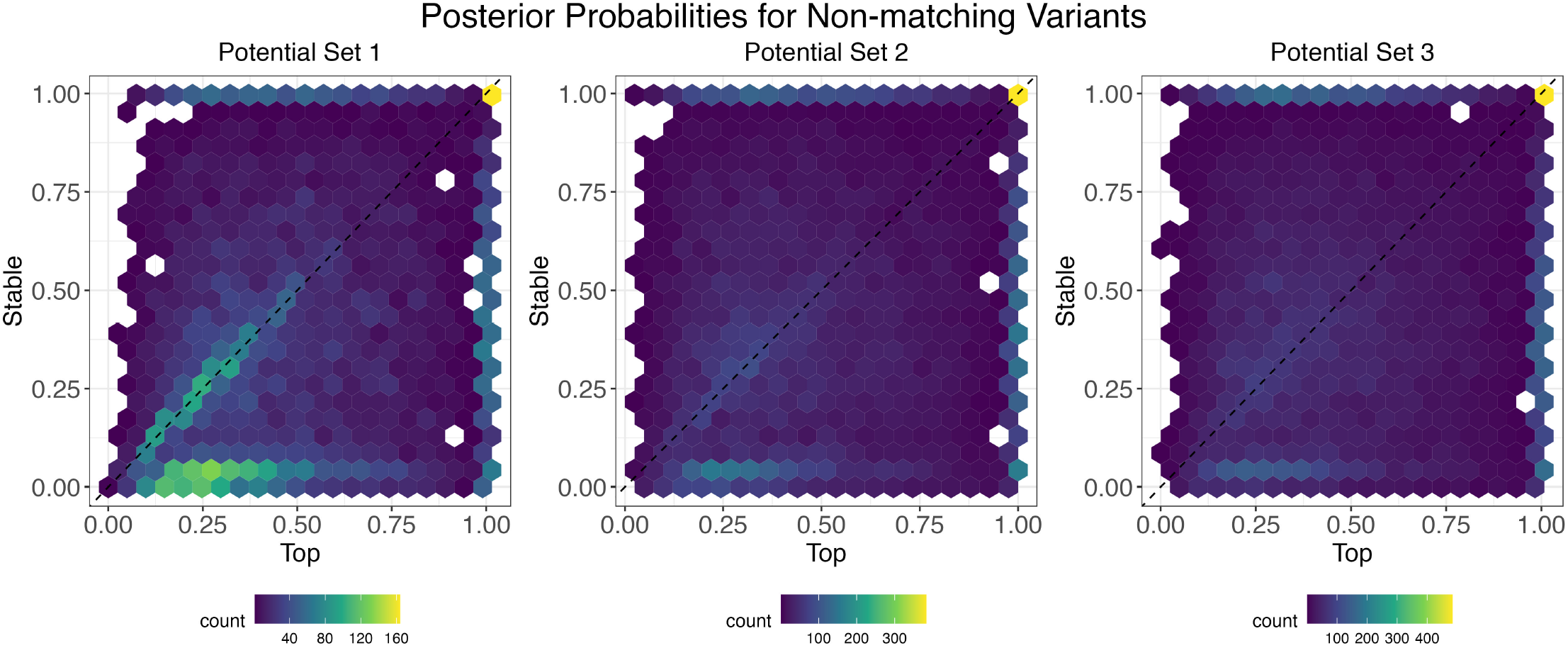
Pair density plot of posterior probabilities of the top variant and the stable variant, in case they do not match.

## Footnotes

1 https://www.ebi.ac.uk/biostudies/files/E-GEUV-1/E-GEUV-1/analysis_results/GeuvadisRNASeqAnalysisFiles_README.txt

## Notes

### Competing Interest Statement

The authors have declared no competing interest.

### Summary of Updates

The revised version includes additional simulation studies and analysis of another fine-mapping method.

## References

1000 Genomes Project Consortium, et al. A global reference for human genetic variation. Nature. 2015; 526(7571):68.

Abdellaoui A, Dolan CV, Verweij KJ, Nivard MG. Gene–environment correlations across geographic regions affect genome-wide association studies. Nature Genetics. 2022; 54(9):1345–1354.

Adzhubei IA, Schmidt S, Peshkin L, Ramensky VE, Gerasimova A, Bork P, Kondrashov AS, Sunyaev SR. A method and server for predicting damaging missense mutations. Nature Methods. 2010; 7(4):248–249.

Avsec Ž, Agarwal V, Visentin D, Ledsam JR, Grabska-Barwinska A, Taylor KR, Assael Y, Jumper J, Kohli P, Kelley DR. Effective gene expression prediction from sequence by integrating long-range interactions. Nature Methods. 2021; 18(10):1196–1203.

Balasubramanian S, Fu Y, Pawashe M, McGillivray P, Jin M, Liu J, Karczewski KJ, MacArthur DG, Gerstein M. Using ALoFT to determine the impact of putative loss-of-function variants in protein-coding genes. Nature Communications. 2017; 8(1):1–11.

Barbadilla-Martínez L, Klaassen N, van Steensel B, de Ridder J. Predicting gene expression from DNA sequence using deep learning models. Nature Reviews Genetics. 2025; p. 1–15.

Basu S, Kumbier K, Brown JB, Yu B. Iterative random forests to discover predictive and stable high-order interactions. Proceedings of the National Academy of Sciences. 2018; 115(8):1943–1948.

Benegas G, Ye C, Albors C, Li JC, Song YS. Genomic language models: opportunities and challenges. Trends in Genetics. 2025;.

Benner C, Spencer CC, Havulinna AS, Salomaa V, Ripatti S, Pirinen M. FINEMAP: efficient variable selection using summary data from genome-wide association studies. Bioinformatics. 2016; 32(10):1493–1501.

Bousquet O, Elisseeff A. Stability and generalization. The Journal of Machine Learning Research. 2002; 2:499–526.

Brown BC, Bray NL, Pachter L. Expression reflects population structure. PLoS Genetics. 2018; 14(12):e1007841.

Cavazos TB, Witte JS. Inclusion of variants discovered from diverse populations improves polygenic risk score transferability. Human Genetics and Genomics Advances. 2021; 2(1):100017.

Cui R, Elzur RA, Kanai M, Ulirsch JC, Weissbrod O, Daly MJ, Neale BM, Fan Z, Finucane HK. Improving fine-mapping by modeling infinitesimal effects. Nature Genetics. 2024; 56(1):162–169.

Cunningham F, Allen JE, Allen J, Alvarez-Jarreta J, Amode MR, Armean IM, Austine-Orimoloye O, Azov AG, Barnes I, Bennett R, et al. Ensembl 2022. Nucleic Acids Research. 2022; 50(D1):D988–D995.

Davenport EE, Amariuta T, Gutierrez-Arcelus M, Slowikowski K, Westra HJ, Luo Y, Shen C, Rao DA, Zhang Y, Pearson S, et al. Discovering in vivo cytokine-eQTL interactions from a lupus clinical trial. Genome Biology. 2018; 19:1–15.

Davydov EV, Goode DL, Sirota M, Cooper GM, Sidow A, Batzoglou S. Identifying a high fraction of the human genome to be under selective constraint using GERP++. PLoS Computational Biology. 2010; 6(12):e1001025.

Efron B, Tibshirani RJ. An Introduction to the Bootstrap. CRC press; 1994.

Farh KKH, Marson A, Zhu J, Kleinewietfeld M, Housley WJ, Beik S, Shoresh N, Whitton H, Ryan RJ, Shishkin AA, et al. Genetic and epigenetic fine mapping of causal autoimmune disease variants. Nature. 2015; 518(7539):337–343.

Favé MJ, Lamaze FC, Soave D, Hodgkinson A, Gauvin H, Bruat V, Grenier JC, Gbeha E, Skead K, Smargiassi A, et al. Gene-by-environment interactions in urban populations modulate risk phenotypes. Nature Communications. 2018; 9(1):1–12.

Fu Y, Liu Z, Lou S, Bedford J, Mu XJ, Yip KY, Khurana E, Gerstein M. FunSeq2: a framework for prioritizing noncoding regulatory variants in cancer. Genome Biology. 2014; 15(10):1–15.

Gao B, Zhou X. MESuSiE enables scalable and powerful multi-ancestry fine-mapping of causal variants in genome-wide association studies. Nature Genetics. 2024; 56(1):170–179.

Gong J, Mei S, Liu C, Xiang Y, Ye Y, Zhang Z, Feng J, Liu R, Diao L, Guo AY, et al. PancanQTL: systematic identification of cis-eQTLs and trans-eQTLs in 33 cancer types. Nucleic Acids Research. 2018; 46(D1):D971–D976.

Griffin JE, Steel MF. Adaptive computational methods for Bayesian variable selection. In: Handbook of Bayesian Variable Selection Chapman and Hall/CRC; 2021. p. 109–130.

Han B, Eskin E. Interpreting meta-analyses of genome-wide association studies. PLoS Genetics. 2012; 8(3):e1002555.

Hormozdiari F, Kichaev G, Yang WY, Pasaniuc B, Eskin E. Identification of causal genes for complex traits. Bioinformatics. 2015; 31(12):i206–i213.

Hu S, Ferreira LA, Shi S, Hellenthal G, Marchini J, Lawson DJ, Myers SR. Fine-scale population structure and widespread conservation of genetic effect sizes between human groups across traits. Nature Genetics. 2025; 57(2):379–389.

Huang C, Shuai RW, Baokar P, Chung R, Rastogi R, Kathail P, Ioannidis NM. Personal transcriptome variation is poorly explained by current genomic deep learning models. Nature Genetics. 2023; 55(12):2056–2059.

Huang YF, Gulko B, Siepel A. Fast, scalable prediction of deleterious noncoding variants from functional and population genomic data. Nature Genetics. 2017; 49(4):618–624.

Ioannidis NM, Davis JR, DeGorter MK, Larson NB, McDonnell SK, French AJ, Battle AJ, Hastie TJ, Thibodeau SN, Montgomery SB, et al. FIRE: Functional inference of genetic variants that regulate gene expression. Bioinformatics. 2017; 33(24):3895–3901.

Karollus A, Mauermeier T, Gagneur J. Current sequence-based models capture gene expression determinants in promoters but mostly ignore distal enhancers. Genome Biology. 2023; 24(1):56.

Katsonis P, Wilhelm K, Williams A, Lichtarge O. Genome interpretation using in silico predictors of variant impact. Human Genetics. 2022; 141(10):1549–1577.

Keys KL, Mak AC, White MJ, Eckalbar WL, Dahl AW, Mefford J, Mikhaylova AV, Contreras MG, Elhawary JR, Eng C, et al. On the cross-population generalizability of gene expression prediction models. PLoS Genetics. 2020; 16(8):e1008927.

Kheradpour P, Kellis M. Systematic discovery and characterization of regulatory motifs in ENCODE TF binding experiments. Nucleic Acids Research. 2014; 42(5):2976–2987.

LaPierre N, Taraszka K, Huang H, He R, Hormozdiari F, Eskin E. Identifying causal variants by fine mapping across multiple studies. PLoS Genetics. 2021; 17(9):e1009733.

Lappalainen T, Sammeth M, Friedländer MR t Hoen PA, Monlong J, Rivas MA, Gonzalez-Porta M, Kurbatova N, Griebel T, Ferreira PG, et al. Transcriptome and genome sequencing uncovers functional variation in humans. Nature. 2013; 501(7468):506–511.

Li S, Sesia M, Romano Y, Candès E, Sabatti C. Searching for robust associations with a multi-environment knockoff filter. Biometrika. 2022; 109(3):611–629.

Liang Y, Pividori M, Manichaikul A, Palmer AA, Cox NJ, Wheeler HE, Im HK. Polygenic transcriptome risk scores (PTRS) can improve portability of polygenic risk scores across ancestries. Genome Biology. 2022; 23(1):1–18.

Lim C, Yu B. Estimation stability with cross-validation (ESCV). Journal of Computational and Graphical Statistics. 2016; 25(2):464–492.

Livesey BJ, Badonyi M, Dias M, Frazer J, Kumar S, Lindorff-Larsen K, McCandlish DM, Orenbuch R, Shearer CA, Muffley L, et al. Guidelines for releasing a variant effect predictor. Genome Biology. 2025; 26(1):97.

Lu Z, Gopalan S, Yuan D, Conti DV, Pasaniuc B, Gusev A, Mancuso N. Multi-ancestry fine-mapping improves precision to identify causal genes in transcriptome-wide association studies. The American Journal of Human Genetics. 2022; 109(8):1388–1404.

Lu Z, Wang X, Carr M, Kim A, Gazal S, Mohammadi P, Wu L, Pirruccello J, Kachuri L, Gusev A, et al. Improved multiancestry fine-mapping identifies cis-regulatory variants underlying molecular traits and disease risk. Nature Genetics. 2025; p. 1–9.

Márquez-Luna C, Loh PR, Consortium SATDS, Consortium STD, Price AL. Multiethnic polygenic risk scores improve risk prediction in diverse populations. Genetic Epidemiology. 2017; 41(8):811–823.

Mazumder R. Discussion of “Best Subset, Forward Stepwise or Lasso? Analysis and Recommendations Based on Extensive Comparisons”. Statistical Science. 2020; 35(4):602–608.

McVicker G, Gordon D, Davis C, Green P. Widespread genomic signatures of natural selection in hominid evolution. PLoS Genetics. 2009; 5(5):e1000471.

Morris AP. Transethnic meta-analysis of genomewide association studies. Genetic Epidemiology. 2011; 35(8):809–822.

Mostafavi H, Harpak A, Agarwal I, Conley D, Pritchard JK, Przeworski M. Variable prediction accuracy of poly-genic scores within an ancestry group. Elife. 2020; 9:e48376.

Ng PC, Henikoff S. SIFT: Predicting amino acid changes that affect protein function. Nucleic Acids Research. 2003; 31(13):3812–3814.

Pollard KS, Hubisz MJ, Rosenbloom KR, Siepel A. Detection of nonneutral substitution rates on mammalian phylogenies. Genome Research. 2010; 20(1):110–121.

Rentzsch P, Witten D, Cooper GM, Shendure J, Kircher M. CADD: predicting the deleteriousness of variants throughout the human genome. Nucleic Acids Research. 2019; 47(D1):D886–D894.

Rogers MF, Shihab HA, Mort M, Cooper DN, Gaunt TR, Campbell C. FATHMM-XF: accurate prediction of pathogenic point mutations via extended features. Bioinformatics. 2018; 34(3):511–513.

Sasse A, Ng B, Spiro AE, Tasaki S, Bennett DA, Gaiteri C, De Jager PL, Chikina M, Mostafavi S. Benchmarking of deep neural networks for predicting personal gene expression from DNA sequence highlights shortcomings. Nature Genetics. 2023; 55(12):2060–2064.

Schaid DJ, Chen W, Larson NB. From genome-wide associations to candidate causal variants by statistical fine-mapping. Nature Reviews Genetics. 2018; 19(8):491–504.

Shi H, Gazal S, Kanai M, Koch EM, Schoech AP, Siewert KM, Kim SS, Luo Y, Amariuta T, Huang H, et al. Population-specific causal disease effect sizes in functionally important regions impacted by selection. Nature Commu nications. 2021; 12(1):1098.

Taylor KE, Ansel KM, Marson A, Criswell LA, Farh KKH. PICS2: next-generation fine mapping via probabilistic identification of causal SNPs. Bioinformatics. 2021; 37(18):3004–3007.

Turley P, Martin AR, Goldman G, Li H, Kanai M, Walters RK, Jala JB, Lin K, Millwood IY, Carey CE, et al. Multi-ancestry meta-analysis yields novel genetic discoveries and ancestry-specific associations. BioRxiv. 2021; p. 2021–04.

Wang G, Sarkar A, Carbonetto P, Stephens M. A simple new approach to variable selection in regression, with application to genetic fine mapping. Journal of the Royal Statistical Society Series B: Statistical Methodology. 2020; 82(5):1273–1300.

Wang QS, Kelley DR, Ulirsch J, Kanai M, Sadhuka S, Cui R, Albors C, Cheng N, Okada Y, Aguet F, et al. Leveraging supervised learning for functionally informed fine-mapping of cis-eQTLs identifies an additional 20,913 putative causal eQTLs. Nature Communications. 2021; 12(1):1–11.

Wen X, Lee Y, Luca F, Pique-Regi R. Efficient integrative multi-SNP association analysis via deterministic approximation of posteriors. The American Journal of Human Genetics. 2016; 98(6):1114–1129.

Wen X, Luca F, Pique-Regi R. Cross-population joint analysis of eQTLs: fine mapping and functional annotation. PLoS Genetics. 2015; 11(4):e1005176.

Willer CJ, Li Y, Abecasis GR. METAL: fast and efficient meta-analysis of genomewide association scans. Bioinformatics. 2010; 26(17):2190–2191.

Yang Z, Wang C, Liu L, Khan A, Lee A, Vardarajan B, Mayeux R, Kiryluk K, Ionita-Laza I. CARMA is a new Bayesian model for fine-mapping in genome-wide association meta-analyses. Nature Genetics. 2023; 55(6):1057– 1065.

Yu B, Kumbier K. Veridical data science. Proceedings of the National Academy of Sciences. 2020; 117(8):3920– 3929.

Yuan K, Longchamps RJ, Pardiñas AF, Yu M, Chen TT, Lin SC, Chen Y, Lam M, Liu R, Xia Y, et al. Fine-mapping across diverse ancestries drives the discovery of putative causal variants underlying human complex traits and diseases. Nature Genetics. 2024; 56(9):1841–1850.

Zaidi AA, Mathieson I. Demographic history mediates the effect of stratification on polygenic scores. Elife. 2020; 9:e61548.

Zhou H, Arapoglou T, Li X, Li Z, Zheng X, Moore J, Asok A, Kumar S, Blue EE, Buyske S, et al. FAVOR: functional annotation of variants online resource and annotator for variation across the human genome. Nucleic Acids Research. 2023; 51(D1):D1300–D1311.

